# Network reciprocity reshaped by environmental knowledge and feedback

**DOI:** 10.64898/2026.05.15.725397

**Authors:** Amit Basak, Maria Kleshnina, Supratim Sengupta

## Abstract

Cooperative interactions often unfold in environments that are shaped by collective behavior, yet how knowledge about such changing environments feeds back into evolutionary dynamics remains poorly understood. While network reciprocity explains how spatial structure enables clusters of cooperators to emerge and grow under certain conditions, it typically ignores how individuals respond to environmental change. Here, we integrate stochastic environmental feedback with network reciprocity to examine how knowledge about environmental state shapes the evolution of cooperation in structured populations. We compare regimes in which individuals either condition their behavior on the current state or remain unaware of it. Under weak selection, we derive a simple condition showing that cooperation is favored when the benefit-to-cost ratio exceeds a modified classic reciprocity threshold accounting for the effect of environmental transitions and state knowledge. Environmental shifts can either promote or hinder cooperation depending on accessibility and fidelity of state knowledge. Counterintuitively, greater knowledge does not universally enhance cooperation: for certain transition rules, state awareness raises the critical threshold for cooperation, a phenomenon we term a “knowledge curse”. Our results reveal that, in an ever-changing environment, cooperation in structured populations emerges from a subtle interplay between environmental feedback and information availability.

---

Persistence of cooperation – paying an individual cost to benefit others – across animal kingdoms remains a central question in evolutionary biology. Decades of research have identified mechanisms such as kin selection [1], direct [2, 3, 4] and indirect reciprocity [5, 6, 7], and network reciprocity [8, 9, 10] that promote cooperation under favorable conditions [11, 12, 13]. Yet most theoretical work assumes static payoff structures or focuses on well-mixed populations, leaving the interplay between ecological environmental dynamics and shifting individual incentives comparatively unexplored [14, 15, 16]. In natural systems, however, interactions unfold within structured populations where social ties and spatial proximity shape strategic behavior [17, 18, 19, 20]. Here, we investigate how environmental feedback and knowledge of the environmental state influence the evolution of cooperation on networks, thereby linking ecological dynamics with network reciprocity.

Network reciprocity provides one of the simplest mechanisms for the evolution of cooperation: unlike direct or indirect reciprocity, it does not rely on repeated interactions, memory, or reputation [11]. Here, population structure enables cooperators to form clusters, thereby preventing cooperators from exploitation through defectors by restricting the interaction opportunity. In its canonical form, cooperation is favored when the benefit-to-cost ratio of an altruistic act exceeds the average number of neighbors [9]. This structural condition makes network reciprocity applicable even to organisms with limited cognitive capacity, including unicellular microbes [21, 22, 23]. However, the same condition implies that increasing connectivity suppresses cooperation, as a higher average degree raises the threshold for the sustainability of cooperation. This prediction appears at odds with densely populated microbial biofilms, where cooperation is frequently observed despite intense local competition [24]. Motivated by this tension and in line with the most recent literature, we relax the standard assumption of a fixed payoff structure and instead allow the environment to dynamically shift interaction incentives, examining how this feedback alters the emergence of cooperation on networks.

Many natural and social systems exhibit feedback between collective behavior and environmental state — from epidemic control [25, 26, 27] and resource management [28, 29, 30] to climate mitigation [31, 32, 33] and microbial community dynamics [34, 35, 36, 37, 38]. In such settings, strategic incentives are not fixed: individual actions reshape the environment, which in turn modifies future payoffs [14, 15, 39]. These coupled dynamics are naturally captured by stochastic games [16, 40, 41, 42], where interactions unfold over time and environmental states (games) evolve in response to behavior and chance, thereby linking present actions to future social dilemmas.

Despite their relevance, stochastic games have only recently been combined with network reciprocity [18, 43]. These studies demonstrate that environmental state transitions can promote cooperation by effectively lowering the critical benefit-to-cost threshold. However, individuals in these models do not condition their strategies on the environmental state; while payoffs depend on the environment, the state itself remains hidden. In many real-world social dilemmas, by contrast, awareness of environmental degradation can critically shape behavior. For example, in the context of climate change, recognition of deteriorating environmental conditions can spur the large-scale cooperative action needed to avert catastrophic outcomes and severe collective losses [44, 45, 46]. Consistent with this intuition, recent work in well-mixed populations shows that informational assumptions can fundamentally alter the conditions for cooperation, sometimes yielding the counterintuitive result that ignorance improves cooperation rate [47]. More broadly, economic theory has long emphasized that information can both enhance and hinder [48] coordination and collective outcomes [49, 50, 51], and studies of partially observable stochastic games demonstrate that incomplete information may, in some cases, improve decision-making [52, 53]. Yet how environmental information interacts with network reciprocity remains unexplored.

We investigate how environmental state uncertainty in stochastic games shapes the evolution of cooperation on networks. We systematically analyze how environmental transition structure influences cooperative dynamics and uncover a fundamental tradeoff between environmental feedback and the sustainability of cooperation in the population of individuals who do not condition their behavior on the state of the environment. We further compare two informational regimes: in a ‘complete knowledge’ setting, individuals observe the environmental state and adopt state-dependent strategies, whereas in a setting ‘without knowledge’, they follow state-independent strategies despite stochastic state transitions. To isolate the role of state knowledge, we consider two environmental states differing in the benefit of cooperation and embed these interactions in structured populations represented by regular graphs.

Under weak selection, we derive a simple condition showing that cooperation is favored when the benefit-to-cost ratio exceeds 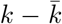, where *k* is the average number of neighbors of an individual and 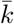 captures how game transitions and information availability modify the classical network reciprocity threshold. We show that environmental dynamics can either promote or hinder cooperation depending on whether individuals observe the environmental state. Strikingly, greater knowledge does not universally enhance cooperation: for some transition rules, access to state knowledge raises the critical threshold, a phenomenon we term a “knowledge curse”. Extending the analysis to noisy information, we find that imperfect state awareness can decisively shape cooperation, particularly in highly connected networks. Our comprehensive analysis of environmental transitions reveals that cooperation in structured populations is jointly shaped by environmental feedback and information availability.

## Results

### Model

We study evolutionary dynamics with environmental state transitions in a finite but large population of *N* individuals arranged on an undirected, unweighted regular graph of degree *k*. The population structure is described by an adjacency matrix 𝒲= (*w*_*ij*_), where *w*_*ij*_ = 1 when two individuals *i* and *j* are connected, and *w*_*ij*_ = 0 otherwise. We introduce environmental state transitions in the simplest setting in which each pair of connected individuals can be in one of two possible states. Depending on the state, they play one of the two Prisoner’s Dilemma (PD) games. In each state *i*, players can choose to either cooperate (*C*) or defect (*D*); cooperators incur a cost *c* to confer a benefit *b*_*i*_ to the other player, whereas defection requires no cost and yields no benefit (Fig. 1a). We assume that state 1 is more beneficial, i.e., *b*_1_ *> b*_2_ *> c* = 1. Although we focus on the PD game to model social dilemmas in each state, the framework extends to general pairwise interactions (see **Supplementary Information**). At each time step, players interact with all their neighbors, and the game played along different edges may differ depending on the local environmental (game) state (Fig. 1b). States along an edge evolve, and the transitions between states are fully characterized by a transition vector

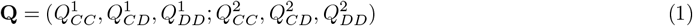

where each entry 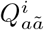 represents the probability that an edge remains in (for *i* = 1) or switches to (for *i* = 2) the more beneficial state (game 1) in the next time step. This probability depends on the current state *i* and the actions *a, ã* ∈ {*C, D*} of the players connected by the edge in that state. We assume that transition probabilities depend only on the number of cooperators on an edge, and not on their identities, such that 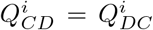. Under this symmetry, there are 2^6^ = 64 distinct deterministic 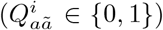 transition vectors. In the main text, we focus primarily on this class of vectors.

**Figure 1.**
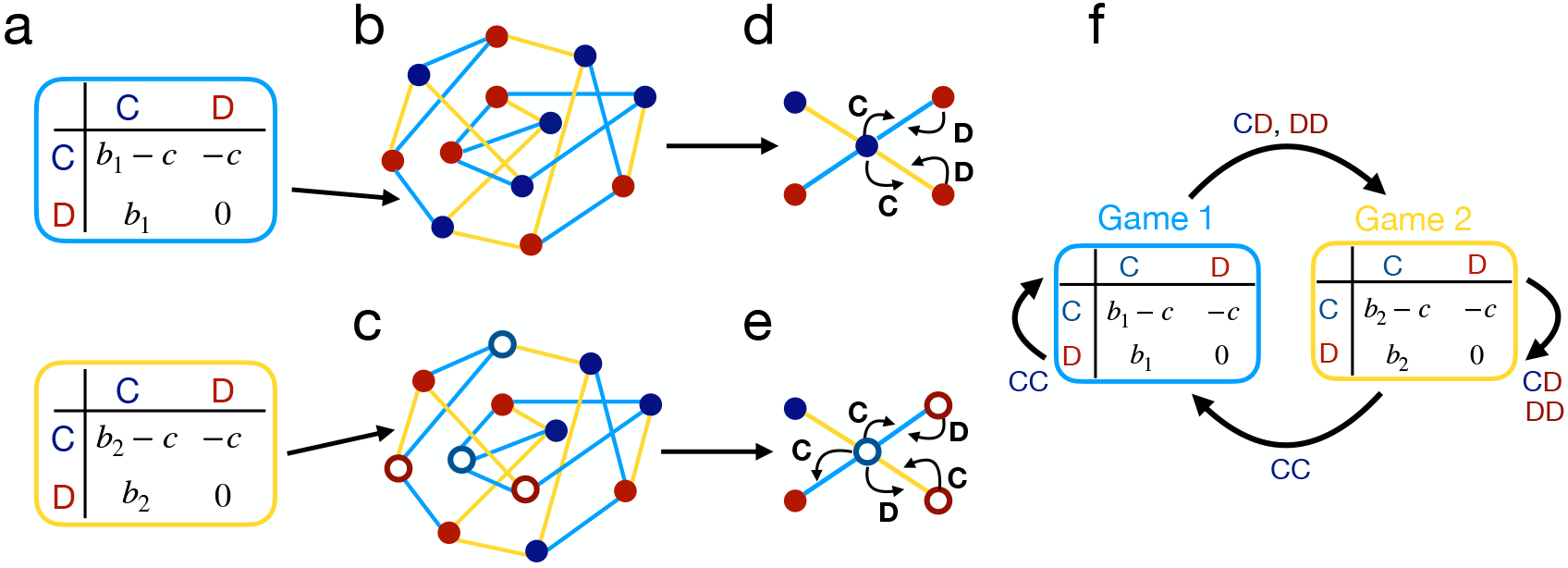
Schematic of game transitions on graphs with and without knowledge of the environmental states. **a** We study game transitions between 2 states represented by two distinct payoff matrices. **b** Each individual is represented by a node on the graph, and exhibits state-independent behavior in the absence of knowledge of the current state, blue (“cooperate” across all states) or red (“defect” across all states), during interactions with neighbors. **c** When individuals are aware of the current state, they can adopt state-dependent behavior, empty blue circle (cooperate in game 1 and defect in game 2) or empty red circle (defect in game 1 and cooperate in game 2), when interacting with neighbors. Games played in interactions with different neighbors can be distinct, represented by the color of the edges; blue edges indicate game 1, and yellow edges denote game 2. **d** An unconditional cooperator (defector) cooperates (defects) in all states, whereas **e** a conditional cooperator (empty blue circle or empty red circle) cooperates depending on the state an edge currently is in. At each time step, individuals interact with their neighbors and accumulate payoffs. After all interactions and payoff accumulations, games are updated along each edge based on both players’ actions and the game they played in the previous interaction. **f** Example of a game transition rule, where only mutual cooperation leads to game 1 and all other actions lead to game 2.

To investigate how state knowledge shapes the evolution of cooperative behavior under state dynamics, we consider two settings: players either observe the current state (Fig. 1c) or lack such information (Fig. 1d). We analyze each setting in detail below.

#### Game transitions with complete state knowledge

When individuals observe the current environmental state prior to the interaction, their strategies can be state-dependent (Fig. 1e), choosing distinct actions in different states. For transitions between two states, a state-dependent strategy is represented as **s**_**d**_ = (*a*_1_, *a*_2_), where *a*_1_, *a*_2_ ∈ { *C, D* } are actions in state 1 and state 2, respectively. In our model, interactions occur only once per evolutionary time step, so strategies do not depend on past actions, and we primarily focus on pure (deterministic) strategies. Thus, the set of pure state-dependent strategies consists of four elements (*n* = 4): { (*C, C*), (*C, D*), (*D, C*), (*D, D*) }. At each time step, players can adopt one of these four possible strategies. For notational simplicity, we refer to these as *CC, CD, DC*, and *DD* strategies for the rest of this article.

In the complete knowledge setting, the population state is given by (*p*_*CC*_, *p*_*CD*_, *p*_*DC*_, *p*_*DD*_), where *p*_**s**_ is the fraction of strategy **s** ∈ { *CC, CD, DC, DD* } present in the population. In the absence of mutations (*µ* = 0), the dynamics admits four absorbing states: *CC* fixation (1, 0, 0, 0); *CD* fixation (0, 1, 0, 0); *DC* fixation (0, 0, 1, 0); and *DD* fixation (0, 0, 0, 1). We quantify the expected level of cooperation by

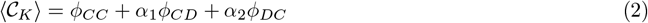

where *ϕ*_**s**_ is the fixation probability of strategy **s** and *α*_1_ (*α*_2_) denote the probability of an edge being in state 1 (state 2) after strategy *CD* (*DC*) has been fixed. Since *CD* (*DC*) cooperates only in state 1 (state 2), the contribution of these strategies to the cooperation level depends on how transition rules shape the long-term distribution of environmental states. For instance, if transitions ensure that all edges occupy state 2 after *CD* fixation (*α*_1_ = 0), the population effectively behaves as defectors. Such cases are excluded when evaluating the cooperation level in Eq. (2). The quantity ⟨𝒞_*K*_⟩ thus represents the probability that a randomly selected interaction is cooperative after the system has reached equilibrium in the complete knowledge scenario.

#### Game transitions without state knowledge

When individuals lack access to environmental state knowledge, or choose to disregard it, they adopt state-independent strategies, using the same action across all states. A pure, state-independent strategy is represented by a single action *a*, written as **s**_**I**_ = (*a*), where *a* ∈ {*C, D* }. A state-independent strategy corresponds to the special case of a state-dependent strategy with *a*_1_ = *a*_2_ = *a*.

In the no knowledge setting, the state of the population is described by a tuple (*p*_*C*_, *p*_*D*_), where *p*_**s**_ is the fraction of strategy **s** ∈ {*C, D*} present in the population. The expected level of cooperation is quantified by

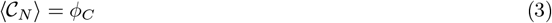

where *ϕ*_*C*_ is the fixation probability of cooperators.

#### Evolutionary dynamics under death-birth updating

At each time step, individuals accumulate pay-offs from interactions with all their neighbors. The accumulated payoff (*π*) determines individual fitness *f* = 1 − *δ* + *δπ*, where *δ* ≥ 0 represents selection strength. After payoff accumulation, an individual *j* is randomly selected for death and replaced by one of her neighbors *l* with probability proportional to *l*’s fitness *f*_*l*_. Following this death-birth update step, the games played along edges are updated according to the transition vector **Q** given by Eq. (1) (Fig. 1f). The individual occupying the vacated site inherits the environmental states resulting from the previous occupant’s interactions.

#### Value of environmental knowledge

We study the evolutionary dynamics in structured populations numerically in the absence of mutations and under weak selection (*µ* = 0, *δ* ≪ 1). For both the complete-knowledge and no knowledge settings, we compute the critical benefit-to-cost ratio 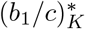 and 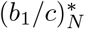, respectively, above which cooperation level (given by Eqs. (2), (3), respectively) exceeds its neutral abundance, 𝒞_*n*_ = 0.5. To quantify the value of state knowledge, we define

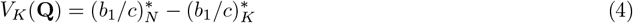

such that a positive value indicates that access to environmental information lowers the threshold for cooperation, whereas a negative value reflects a knowledge curse.

#### Specification for numerical simulations

In numerical simulations, we vary the benefit of the more beneficial state (*b*_1_), while keeping the difference between the benefits of the two states (Δ*b* = *b*_1_ − *b*_2_) fixed (see **Supplementary Information**, Fig. S3 for variable Δ*b*). The population is structured as a random regular graph of degree *k* and initialized in an unbiased state with all strategies equally represented. Accordingly, the initial values for strategy frequencies are (1*/*4, 1*/*4, 1*/*4, 1*/*4) in the complete knowledge settings, and (1*/*2, 1*/*2) in the setting without knowledge. This ensures a consistent comparison between the two informational regimes.

### General rule for the evolution of cooperation

The expected level of cooperation in both scenarios with (Eq. (2)) and without state knowledge (Eq. (3)) is determined by the fixation probabilities of different cooperative strategies. We use analytically derived expressions for the fixation probabilities in the rare-mutation (*µ* → 0) and weak-selection (*δ* ≪ 1) limit to identify conditions for the emergence of cooperation in each case.

Let *ϕ*_*C*_(*y*) denote the fixation probability of cooperators starting from an initial fraction *p*_*C*_(*t* = 0) = *y*, and *ϕ*_*D*_(*y*) be the similar fixation probability for defectors. In the absence of state knowledge, cooperation is favored over defection ( ⟨𝒞 _*N*_ ⟩*>* 0.5) if *ϕ*_*C*_(*y*) *> ϕ*_*D*_(*y*). Using the relation *ϕ*_*D*_(*y*) = 1 − *ϕ*_*C*_(1 − *y*), this condition reduces to *ϕ*_*C*_(*y* = 1*/*2) *>* 1*/*2. We use this condition to numerically determine the critical benefit-to-cost ratio 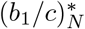. To obtain an analytical approximation under weak selection, we instead compare the fixation probabilities of a single cooperator and a single defector, denoted by *ρ*_*C*_ = *ϕ*_*C*_(1*/N* ) and *ρ*_*D*_ = *ϕ*_*D*_(1*/N* ), respectively. The condition *ρ*_*C*_ *> ρ*_*D*_ then determines the critical ratio 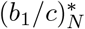 [54].

When actions are conditioned on state knowledge, a strategy **s** is favored by selection if the fixation probability of **s** exceeds the neutral abundance. Since the population is initialized in the state with an equal fraction of all strategies (*p*_**s**_(*t* = 0) = 1*/n*), this condition becomes *ϕ*_**s**_(1*/n*) *>* 1*/n*. We use this condition to numerically determine the threshold 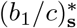 above which a state-dependent strategy **s** is favored by selection. For analytical results in the rare-mutation and weak-selection limit, we consider pairwise competition between **s** and each alternative strategy **s**′ ∈ { *CC, CD, DC, DD* }. The condition for **s** to be favored is then ∑_**s**_*′ ρ*_**ss**_*′* − *ρ*_**s**_*′*_**s**_ *>* 0 [55], where *ρ*_**ss**_*′* denotes the fixation probability of a single **s** individual in a population of (*N* − 1) **s**′ individuals (Details are provided in the **Methods** section).

In the following, we present the results for the basic model. Additional results for alternative game parameters (Fig. S1–S3), evolutionary dynamics with rare mutations (Fig. S4), and stochastic transition vectors (Fig. S5) are provided in the **Supplementary Information**.

### Effect of state knowledge for three different transition rules

We begin by examining how environmental knowledge influences cooperation under three representative deterministic transition rules. Each example captures a “tragedy of the commons”, where mutual cooperation improves the environment by leading to a more beneficial state, while partial defection can sometimes drive it towards a less beneficial state [16].

#### Q_**1**_ = (**1, 0, 0**; **1, 0, 0**)

We first consider a transition rule in which only mutual cooperation maintains the more beneficial state, whereas any defection leads to a less beneficial state (**Q**_**1**_, Fig. 2a). We find that the critical benefit-to-cost ratio (*b*_1_*/c*)^∗^ required to exceed the neutral abundance of cooperation ( 𝒞_*n*_ = 0.5, indicated by the black horizontal line) remains essentially unchanged when we switch between complete and no knowledge scenarios (Fig. 2b), indicating that environmental knowledge does not enhance cooperation under this strict transition rule.

**Figure 2.**
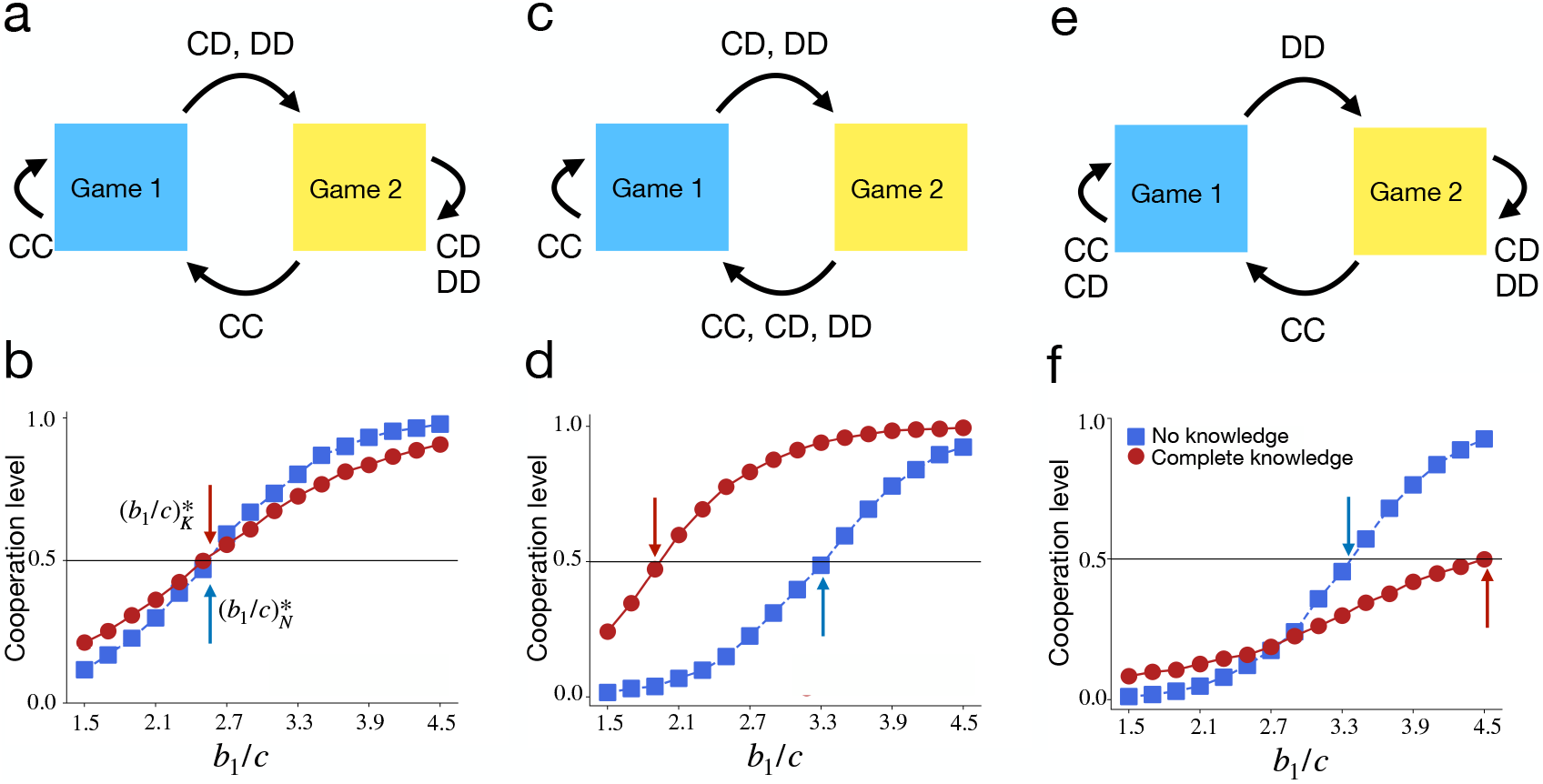
Comparison of complete and no knowledge scenario on the evolution of cooperation with game transitions for three different transition rules. We consider game transitions between two states with deterministic transitions across three different transition rules referred to as **a** natural transition **Q**_**1**_ = (**1, 0, 0**; **1, 0, 0**), **c** timeout rule **Q**_**2**_ = (**1, 0, 0**; **1, 1, 1**), and **e** conditional timeout with conditional return: **Q**_**3**_ = (**1, 1, 0**; **1, 0, 0**). The bottom three panels show the cooperation level under scenarios with state knowledge (red circles) and without state knowledge (blue squares), as a function of the benefit-to-cost ratio in game 1 for each transition rule. **b** For **Q**_**1**_, the critical benefit-to-cost ratio (*b*_1_*/c*)^∗^ for the evolution of cooperation (intersection of dots and the horizontal line) in the complete knowledge scenario is effectively unchanged relative to its value in the no knowledge scenario. **d** Under the timeout rule **Q**_**2**_, knowledge of the environmental states dramatically lowers the (*b*_1_*/c*)^∗^ value compared to the no knowledge baseline. **f** By contrast, for the transition vector **Q**_**3**_, state knowledge increases the (*b*_1_*/c*)^∗^ value beyond the threshold for the no knowledge case, making the evolution of cooperation more difficult when individuals take decisions informed by the current state. The blue arrow indicates the critical threshold 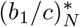 under no knowledge setting, the red arrow denotes the corresponding threshold 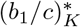 in the complete knowledge scenario. In each simulation, all individuals are initially in state 1. Each simulation runs till one of the strategies fixes in the population. Each point is averaged over 10^4^ independent trials. Other parameters: *N* = 500, *k* = 4, *δ* = 0.01, *b*_2_ = *b*_1_ − 1, *c* = 1.

In the no knowledge setting, cooperation is favored when

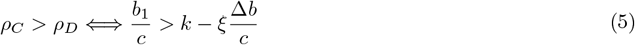

where, *ξ* = (*k* − 1)*/*2 is a positive quantity for the natural transition rule **Q**_**1**_ (Fig. 3a), consistent with previous findings [18]. This shows that environmental transitions already reduce the threshold relative to the static case. Here, *ξ* captures the effect of environmental transitions and depends on the update rule, but not on payoff parameters. When Δ*b* = 0, this reduces to the classical condition *b*_1_*/c > k* [9]. For Δ*b >* 0, environmental transitions lower the cooperation threshold even without state knowledge.

**Figure 3.**
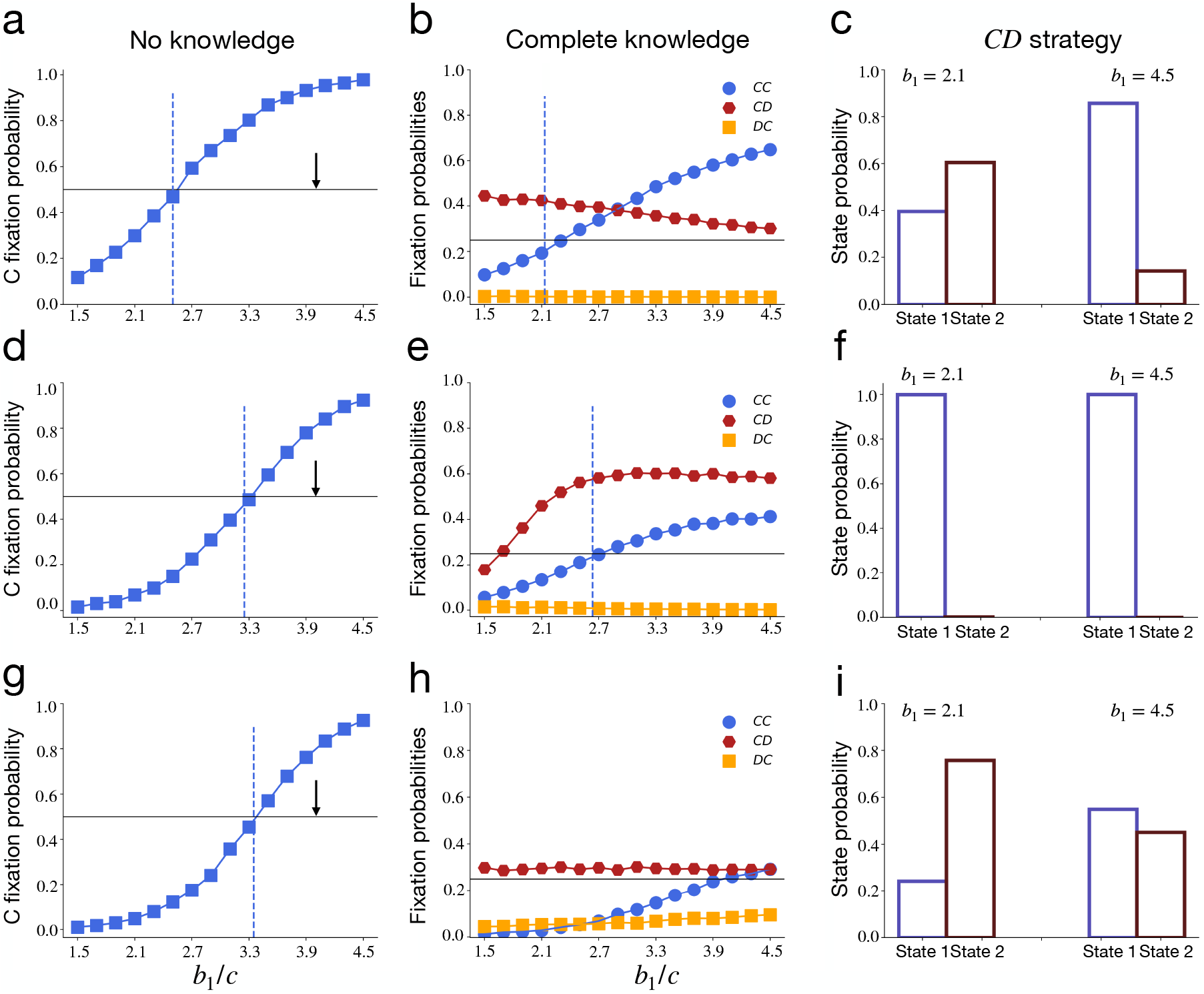
Fixation probability of different cooperative strategies in no knowledge and complete knowledge scenarios for three different transition rules. The first, second and third row corresponds to **(i) Q**_**1**_ = (**1, 0, 0**; **1, 0, 0**), **(ii) Q**_**2**_ = (**1, 0, 0**; **1, 1, 1**) and **(iii) Q**_**3**_ = (**1, 1, 0**; **1, 0, 0**) respectively. **a, d, g:** Fixation probability of cooperators in the no knowledge scenario, and **b, e, h:** Fixation probability of different cooperative strategies (*CC, CD*, and *DC*) for the complete knowledge scenario, as a function of the benefit-to-cost ratio in game 1. **c, f, i** shows the probability of an edge to be in state 1 or state 2 for the leading state-dependent cooperative strategy *CD* for each transition vector. In each figure, dots show simulation results, and the vertical dashed line shows the analytical result. The intersection of the horizontal line and dots gives the critical benefit-to-cost ratio (*b*_1_*/c*)^∗^ for each type of strategy to cross its neutral abundances (1/2 for the system without knowledge and 1/4 in the complete knowledge scenario). The blue vertical dashed line represents the analytical value of 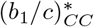 for the strategy *CC* to be favored by selection in both systems with and without state knowledge. The vertical black arrow shows the location of the critical threshold (*b*_1_*/c*)^∗^ for a single game when there are no game transitions. The fixation probability has been calculated as the fraction of times each strategy gets fixed in the population out of 10^4^ trials. Each simulation starts with an equal proportion of available strategies in each case and runs till one of the strategies is fixed in the population. Parameters are the same as in Fig. 2.

In the complete-knowledge setting, both *CC* and *CD* strategies can be favored (Fig. 3b). The condition for a cooperative strategy **s** ∈ {*CC, CD, DC*} to be favored follows

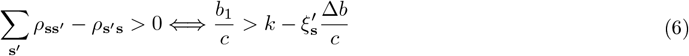

where, depending on the strategy **s** and transition rule (**Q**) under consideration, 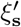 can be positive or negative, and it may depend on the game parameters *b*_1_, *b*_2_, and *c*. For 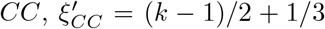, implying a lower threshold compared to the no knowledge setting. However, the *CD* strategy is also strongly favored, and its fixation probability exceeds that of *CC*.

Crucially, the contribution of *CD* to cooperation depends on the equilibrium distribution of environmental states. Figure 3c shows that for low *b*_1_, most edges equilibrate in state 2 (*α*_1_ ≈ 0) after *CD* fixation. Thus, despite being favored by selection, *CD* contributes little to cooperation (see Eq. (2)). As a result, the reduced threshold for *CC* is offset by the dominance of *CD* in the less beneficial state, and the net effect of environmental knowledge is neutral.

#### Q_**2**_ = (**1, 0, 0**; **1, 1, 1**)

We next consider a ‘time-out’ transition rule (Fig. 2c), where defection moves the game to a less beneficial state, but the game automatically returns to the more beneficial state in the following step [47]. In the absence of state knowledge, cooperation is favored when Eq. (5) holds with *ξ* = (*k* − 1)*/*4, which is half the value for **Q**_**1**_. Consequently, the threshold 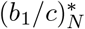 is higher (Fig. 2d).

In contrast, in the complete knowledge setting (Fig. 3e), both the *CC* and *CD* strategies are favored under significantly less restrictive conditions. For 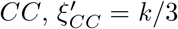 and 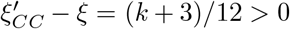. This effect becomes more pronounced for large *k*. The *CD* strategy is even more strongly favored, with a very low critical threshold since for 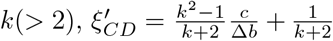 is a large positive quantity when the two games slightly differ from each other (Δ*b >* 0). Figure 3f shows that when *CD* fixates, almost all edges remain in state 1 (*α*_1_ ≃ 1). Thus, *CD* both avoids exploitation in the less beneficial state and promotes recovery to the more beneficial state. The combination of a lower threshold for both *CC* and *CD*, coupled with a high abundance of *CD* in state 1, ensures a strong positive effect of environmental knowledge on cooperation.

#### Q_**3**_ = (**1, 1, 0**; **1, 0, 0**)

Finally, we consider a transition rule (Fig. 2e) where cooperation by at least one player in the more beneficial state maintains that state, but mutual defection leads to a less beneficial state. In the absence of state knowledge, cooperation is favored under Eq. (5) with a positive *ξ* = (*k*^3^ − 2)*/*6*k*^2^, implying that environmental transitions lower the threshold for cooperation (Fig. 3g).

However, when individuals condition their behavior on the state, the threshold for cooperation increases substantially (Fig. 2f). In particular, the coefficient 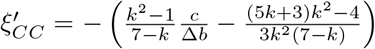 becomes negative, making it significantly harder for *CC* to be favored. Although the *CD* strategy is always favored when *b*_1_ *> b*_2_ (Fig. 3h), its contribution to cooperation depends on the state distribution. Figure 3i shows that for low *b*_1_, most edges equilibrate in state 2 after *CD* fixation, reducing its contribution to the cooperation level. Thus, under **Q**_**3**_, environmental knowledge increases the cooperation threshold and suppresses cooperative outcomes, which can be interpreted as a manifestation of the knowledge curse.

These examples provide two key insights. First, the effect of environmental transitions on cooperation depends strongly on the specific transition rule. Second, environmental knowledge can act as a double-edged sword, sometimes promoting but at other times either hindering or having no effect on cooperation. These three examples demonstrate that the evolution of cooperation in structured populations can be sensitive to both environmental knowledge and feedback manifest through game transitions.

### Comprehensive analysis across all transition rules

To assess the impact of environmental state knowledge on the evolution of cooperation, we first establish how game transitions influence cooperation dynamics in the absence of such knowledge. To this end, we build on the mathematical framework of Su *et al* [18] and extend the analysis to all deterministic transition rules. These results serve as a benchmark for comparison with the complete-knowledge setting, allowing us to identify when access to state knowledge promotes or hinders cooperation.

#### Game transitions without state knowledge across transition rules

In the absence of state knowledge, individuals choose either a pure cooperator (*C*) or defector (*D*) strategy uniformly across all edges, and play the one-shot donation game according to the state they are currently in. In this scenario, Eq. (5) provides the analytical condition for cooperators to be favored by selection over defection under the death-birth update.

For a given transition rule, the value of *ξ* in Eq. (5) determines whether game transitions have a beneficial, detrimental, or no effect on cooperation. A positive value of *ξ* lowers the critical benefit-to-cost ratio (*b*_1_*/c*)^∗^ relative to the baseline case without transitions (*b*_1_*/c*)^∗^ = *k*, whereas a negative value increases this threshold, making cooperation more difficult to evolve. By analyzing all 2^6^ = 64 deterministic transition vectors, we find that *ξ* can be positive, negative, or zero depending on the transition structure (see **Supplementary Information**, section 2, Table 1). In particular, only 14 transition rules yield *ξ >* 0, indicating that such environmental transitions promote cooperation, while the remaining rules either have no effect (*ξ* = 0) or hinder cooperation (*ξ <* 0) when individuals are unaware of the states (Fig. 4a). Below, we classify the transition rules that lead to *ξ >* 0.

**Table 1:** Values of *ξ* for each deterministic transition vector **Q** (first 32 entries in left two columns, remaining 32 in right two columns).

| $\mathbf{Q}$ | $\xi$ | $\mathbf{Q}$ | $\xi$ |
| --- | --- | --- | --- |
| (0,0,0;0,0,0) | -1 | (1,0,0;0,0,0) | -1 |
| (0,0,0;0,0,1) | -1 | (1,0,0;0,0,1) | -1 |
| (0,0,0;0,1,0) | $-\frac{k+3}{4}$ | (1,0,0;0,1,0) | $-\frac{1}{2}$ |
| (0,0,0;0,1,1) | $-\frac{k+3}{4}$ | (1,0,0;0,1,1) | $-\frac{1}{2}$ |
| (0,0,0;1,0,0) | $\frac{k-3}{4}$ | (1,0,0;1,0,0) | $\frac{k-1}{2}$ |
| (0,0,0;1,0,1) | $\frac{k-3}{4}$ | (1,0,0;1,0,1) | $\frac{k-1}{2}$ |
| (0,0,0;1,1,0) | $-\frac{1}{2}$ | (1,0,0;1,1,0) | $\frac{k-1}{4}$ |
| (0,0,0;1,1,1) | $-\frac{1}{2}$ | (1,0,0;1,1,1) | $\frac{k-1}{4}$ |
| (0,0,1;0,0,0) | -1 | (1,0,1;0,0,0) | -1 |
| (0,0,1;0,0,1) | -1 | (1,0,1;0,0,1) | -1 |
| (0,0,1;0,1,0) | $-\frac{k+3}{4}$ | (1,0,1;0,1,0) | $-\frac{1}{2}$ |
| (0,0,1;0,1,1) | $-\frac{k+3}{4}$ | (1,0,1;0,1,1) | $-\frac{1}{2}$ |
| (0,0,1;1,0,0) | $\frac{k-3}{4}$ | (1,0,1;1,0,0) | $\frac{k-1}{2}$ |
| (0,0,1;1,0,1) | $\frac{k-3}{4}$ | (1,0,1;1,0,1) | $\frac{k-1}{2}$ |
| (0,0,1;1,1,0) | $-\frac{1}{2}$ | (1,0,1;1,1,0) | $\frac{k-1}{4}$ |
| (0,0,1;1,1,1) | $-\frac{1}{2}$ | (1,0,1;1,1,1) | $\frac{k-1}{4}$ |
| (0,1,0;0,0,0) | -1 | (1,1,0;0,0,0) | -1 |
| (0,1,0;0,0,1) | $-\frac{k}{12} - 1 + \frac{1}{6k^2}$ | (1,1,0;0,0,1) | $-\frac{1}{2}$ |
| (0,1,0;0,1,0) | $-\frac{k-1}{2}$ | (1,1,0;0,1,0) | 0 |
| (0,1,0;0,1,1) | $-\frac{k-1}{2}$ | (1,1,0;0,1,1) | 0 |
| (0,1,0;1,0,0) | $\frac{k}{12} - \frac{1}{2} - \frac{1}{6k^2}$ | (1,1,0;1,0,0) | $\frac{k^3-2}{6k^2}$ |
| (0,1,0;1,0,1) | $-\frac{1}{2}$ | (1,1,0;1,0,1) | $\frac{k^3-2}{12k^2}$ |
| (0,1,0;1,1,0) | $-\frac{k-1}{4}$ | (1,1,0;1,1,0) | 0 |
| (0,1,0;1,1,1) | $-\frac{k-1}{4}$ | (1,1,0;1,1,1) | 0 |
| (0,1,1;0,0,0) | -1 | (1,1,1;0,0,0) | 0 |
| (0,1,1;0,0,1) | $-\frac{k}{6} - 1 + \frac{1}{3k^2}$ | (1,1,1;0,0,1) | 0 |
| (0,1,1;0,1,0) | $-\frac{k-1}{2}$ | (1,1,1;0,1,0) | 0 |
| (0,1,1;0,1,1) | $-\frac{k-1}{2}$ | (1,1,1;0,1,1) | 0 |
| (0,1,1;1,0,0) | $-\frac{1}{2}$ | (1,1,1;1,0,0) | 0 |
| (0,1,1;1,0,1) | $-\frac{k}{12} - \frac{1}{2} + \frac{1}{6k^2}$ | (1,1,1;1,0,1) | 0 |
| (0,1,1;1,1,0) | $-\frac{k-1}{4}$ | (1,1,1;1,1,0) | 0 |
| (0,1,1;1,1,1) | $-\frac{k-1}{4}$ | (1,1,1;1,1,1) | 0 |

**Figure 4.**
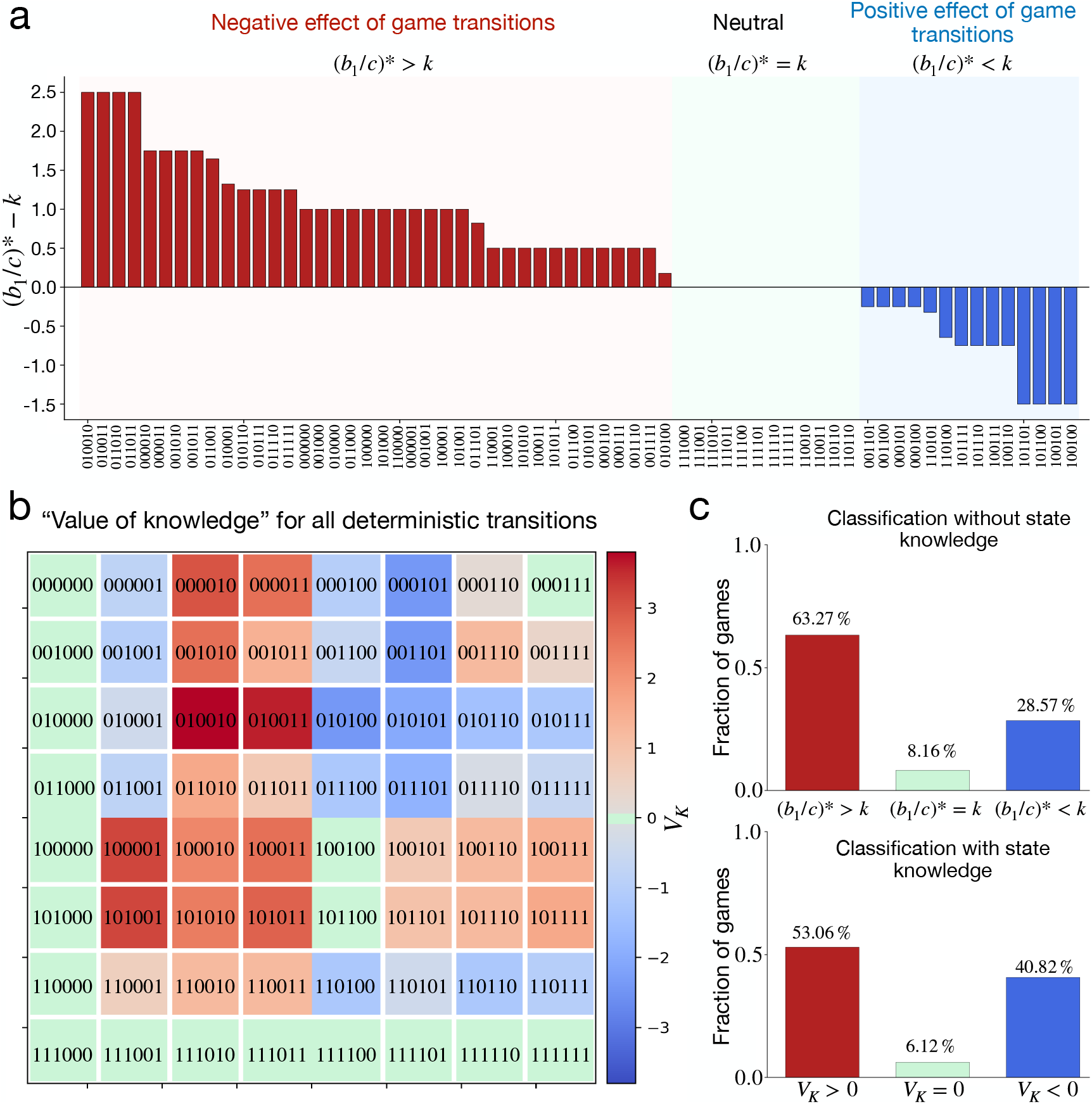
Classification of games without and with state knowledge across all deterministic transition rules. **a** In the absence of state knowledge, game transitions can have a positive, neutral, or negative effect on cooperation depending on the value of (*b*_1_*/c*)^∗^ − *k*. When (*b*_1_*/c*)^∗^ − *k* is a negative quantity, game transitions have a beneficial effect on cooperation, whereas a positive value indicates a detrimental effect on cooperation. The bar diagram shows the respective value of (*b*_1_*/c*)^∗^ − *k* for each of the 64 deterministic transition vectors. **b** When individuals learn about the current state, the value of state knowledge is quantified by comparing the critical benefit-to-cost ratio under both scenarios with and without state knowledge, 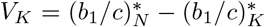. There is a benefit of state knowledge when *V*_*K*_ *>* 0 (denoted by shades of red). When *V*_*K*_ = 0, the game is neutral (denoted by green) and in all other cases, there is a “knowledge curse”, *V*_*K*_ *<* 0 (denoted by shades of blue). The colormap shows the value of knowledge, *V*_*K*_, across all deterministic transition rules. **c** The upper panel shows, the fraction of cases where game transitions in the absence of state knowledge have a positive (blue bar), neutral (green bar), or negative (red bar) effect on cooperation; the lower panel shows the fraction of cases in which state knowledge is beneficial (red bar), detrimental (blue bar), or has no effect (green bar) on the success of cooperation; among the 49 transition vectors with no absorbing states (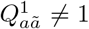 or 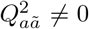 for all *a, ã*). Other parameters are the same as in Fig. 2.

##### First class: mutual cooperation rewarded in both states

In the first class, mutual cooperation leads to a more beneficial state from both environments ( 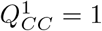 and 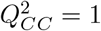). Within this class, we identify four subclasses based on how unilateral cooperation is treated.

i. In the first subclass, unilateral cooperation/defection is penalized in both states (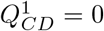 and 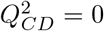), ensuring that unilateral deviation from mutual cooperation leads to a less beneficial state. The corresponding transition vectors are

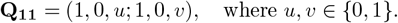

Under this subclass, the beneficial effect of game transitions on cooperation is most pronounced, as the threshold (*b*_1_*/c*)^∗^ is reduced by the largest factor *ξ* = (*k* − 1)*/*2, compared to a single game.
ii. In the second subclass, unilateral defection is penalized in the more beneficial state 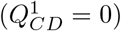, but once an edge is in the less beneficial state, unilateral cooperation allows transition back to the more beneficial state 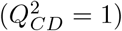. The corresponding transition vectors are

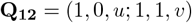

In this case, *ξ* = (*k* − 1)*/*4, which is smaller than in the first subclass by a factor of 1*/*2.
iii. In the third subclass, an edge remains in the more beneficial state if at least one of the connected players cooperates 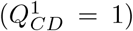, while transitions from state 2 occur only under mutual cooperation 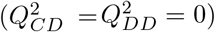. There is only one transition vector in this subclass,

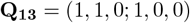

with ξ = (*k*^3^ − 2)/(6*k*^2^)
iv. The fourth subclass is a variant of the third, where coordination in a less beneficial state allows restoration of the more beneficial state 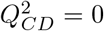 while 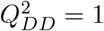. The respective transition vector is

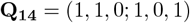

for which *ξ* = (*k*^3^ − 2)*/*(12*k*^2^), indicating the weakest beneficial effect of environmental transitions.

##### Second class: mutual cooperation rewarded only in the disadvantageous state

In the second class, mutual cooperation is rewarded only in the less beneficial state 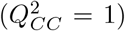, while cooperation in the more beneficial state does not ensure persistence in that state 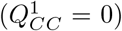. For these transition rules to promote cooperation, unilateral cooperation must be penalized in both states 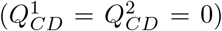. These transition vectors can be written as

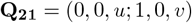

For these transition vectors, the beneficial effect of game transitions depends on the network degree, with *ξ* = (*k* − 3)*/*4 *>* 0 for *k >* 3. We interpret this as a ‘time-in’ mechanism, where the more beneficial state is accessible only transiently through coordination in the less beneficial state.

The above analysis shows that the beneficial effects of game transitions on cooperation in the absence of state knowledge depend critically on how mutual cooperation is rewarded across states and how unilateral cooperation is treated. In particular, transition rules that reward mutual cooperation and penalize unilateral defection in both states (class **Q**_**11**_) are most effective in promoting cooperation. By contrast, rewarding unilateral cooperation in either state 2 (subclass *Q*_12_) or state 1 (subclass *Q*_13_) weakens this effect. Notably, even within the class 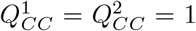, rewarding unilateral cooperation in *both* states 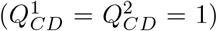 leads to *ξ* = 0, providing no advantage for cooperators (Fig. 4a). Our analytical predictions (indicated by the vertical red line in **Supplementary** Fig. S1) are in good agreement with the numerical results across all transition vectors.

#### Impact of state knowledge across transition rules

In this section, we identify those transition rules for which environmental state knowledge facilitates the spread of cooperation more easily compared to the case where knowledge of the states is unavailable. We classify all transition vectors according to the value of knowledge, *V*_*K*_ (Eq. (4)).

Transition rules with an absorbing state, that is, those for which no further transitions occur once a particular state is reached (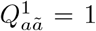 or 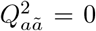 for all *a, ã*), are neutral with respect to state knowledge. There are 15 such cases. Since the system becomes trapped in a single state, knowledge of the environmental state does not influence evolutionary outcomes.

Across all 64 deterministic transition rules, we find that in 26 cases (Fig. 4b), access to environmental information substantially promotes cooperation relative to the no knowledge setting (see also **Supplementary** Fig. S2). In these cases, knowledge either lowers the critical benefit-to-cost ratio required for cooperation to exceed its neutral abundance or enables cooperation to emerge under conditions where state-independent behavior would otherwise inhibit it. Excluding the 15 neutral cases with absorbing states, we focus on the remaining 49 transition rules that allow behavior-dependent state transitions. Within this set, we identify several distinct classes in which state knowledge has beneficial, neutral, or detrimental effects.

##### First class: no reward for mutual cooperation in the less beneficial state

We first consider transition rules for which mutual cooperation in state 2 does not lead to state 1 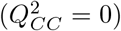. These rules can be written as

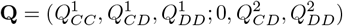

There are 32 such transition vectors. After excluding the absorbing cases 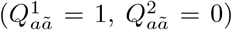, 21 transition rules remain that allow state transitions.

For 17 of these 21 cases (Fig. 4b), we find *V*_*K*_ *>* 0, indicating that state knowledge lowers the critical threshold for cooperation. The remaining four cases correspond to transition vectors of the form

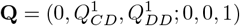

in which neither mutual cooperation nor unilateral cooperation in state 2 enables a transition to the beneficial state. In these cases, conditioning behavior on state knowledge hinders cooperation compared to the no-knowledge setting.

##### Second class: mutual cooperation rewarded in the less beneficial state

When mutual cooperation in state 2 leads to state 1 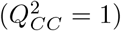, the effect of state knowledge depends sensitively on how mutual cooperation in the more beneficial state and unilateral cooperation in both states are rewarded or penalized. We first consider the subclass where 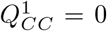. If unilateral defection in state 1 is penalized 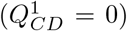, the impact of knowledge depends on whether unilateral cooperation is rewarded and mutual defection is penalized in the less beneficial state. These transition vectors can be written as

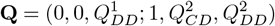

When 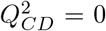, we identify four cases in which cooperation is more easily established in the absence of state knowledge. In contrast, when 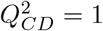, knowledge promotes cooperation provided that 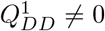 and 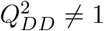. If both 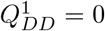 and 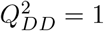, transitions between states occur independently of players’ actions, resulting in neutral outcomes with respect to knowledge.

Within the same class 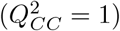, if unilateral cooperation in state 1 is rewarded 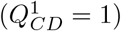, cooperation is favored more strongly when individuals do not condition their behavior on state information. This classification holds regardless of whether mutual cooperation in state 1 is rewarded or penalized. Excluding absorbing cases, this class includes 12 transition vectors of the form

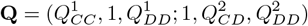

##### Third class: mutual cooperation rewarded in both states

Finally, when mutual cooperation is rewarded in both states (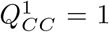 and 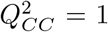) and unilateral defection in the more beneficial state leads to a less beneficial state 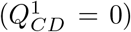, state knowledge promotes cooperation (*V*_*K*_ *>* 0) provided that transitions from state 2 to state 1 occur under either unilateral or mutual defection. These transition rules are given by

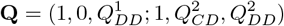

with 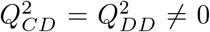, yielding six such cases. However, when transitions to state 1 from state 2 occur only under mutual cooperation, the outcome is neutral with respect to state knowledge.

Our systematic analysis across all deterministic transition rules reveals general patterns governing the role of environmental knowledge. When mutual cooperation in state 2 is not rewarded 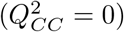, the absence of state knowledge hinders cooperation. When mutual cooperation is rewarded in at least one state, the effect of state knowledge depends strongly on how unilateral cooperation is treated. In particular, if unilateral cooperation in the more beneficial state is rewarded, cooperation can emerge more readily when players ignore state information. Among the 49 non-neutral transition rules, state knowledge promotes cooperation in 53% of cases, while the absence of knowledge is advantageous in 41%, with the remaining 6% yielding neutral outcomes (Fig. 4c). Notably, even among the transition rules where state-independent behavior proves detrimental to cooperation, state-dependent behavior promotes cooperation in 58% of those cases.

### Evolutionary dynamics under imperfect environmental state knowledge

So far, our analysis has considered two limiting cases: complete knowledge of the environmental state, where individuals condition their behavior on the current state, and no knowledge, where behavior is state-independent. In many real-world settings, however, information about the environment is imperfect, and individuals typically make decisions based on uncertain or imperfect knowledge. It is therefore important to understand how imperfect knowledge of the environmental state affects evolutionary dynamics under game transitions.

To model imperfect knowledge, we assume that a player may misperceive her true current state *j* with a probability *n*_*j*_ (and correctly with probability 1 − *n*_*j*_) [56]. When noise is symmetric, *n*_1_ = *n*_2_ = *n*; otherwise *n*_1_ ≠ *n*_2_. Under imperfect knowledge, strategies become effectively stochastic, as actions are conditioned on perceived rather than true states (see **Methods** section).

We focus on the impact of imperfect knowledge under the transition rule **Q**_**1**_ = (1, 0, 0; 1, 0, 0) and compare it to the complete-knowledge case (*n*_1_ = *n*_2_ = 0). Results for other transition rules are presented in **Supplementary** Fig. S6. Simulations on a random regular graph with degree *k* = 4 show that introducing small, symmetric noise in state knowledge (*n*_1_ = *n*_2_ = 0.1) does not alter the critical benefit-to-cost ratio (*b*_1_*/c*)^∗^ required for cooperation to exceed its neutral abundance ( 𝒞_*n*_ = 0.5, Fig. 5a). For the parameter values considered, the expected level of cooperation remains essentially unchanged compared to the complete-knowledge case, indicating that symmetric noise has a small effect under the transition vector **Q**_**1**_. In contrast, asymmetric noise (*n*_1_ ≠ *n*_2_) can have a non-trivial impact on cooperation. Figure 5b shows that when the more beneficial state is perceived with higher accuracy (*n*_1_ = 0.01) than the less beneficial state (*n*_2_ = 0.1), the critical benefit-to-cost ratio decreases substantially: from 2.5 in the complete-knowledge case to 1.9. Conversely, when state 2 is perceived more accurately (*n*_2_ = 0.01) than state 1 (*n*_1_ = 0.1; Fig. 5c), the threshold increases to 3.3, thereby hindering cooperation.

**Figure 5.**
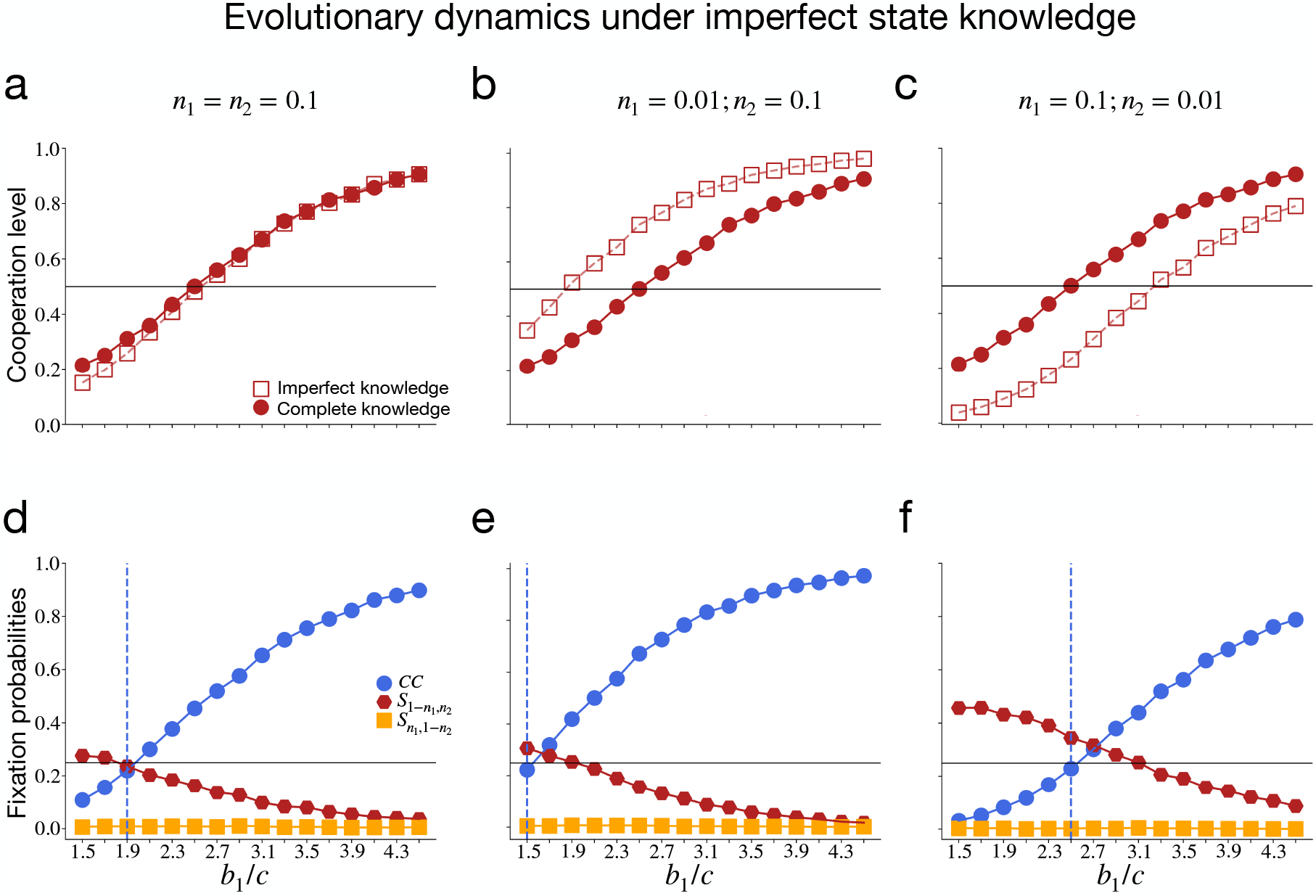
Evolutionary dynamics with imperfect knowledge of states under the transition vector Q_1_ = (1, 0, 0; 1, 0, 0). a-c To illustrate the effect of imperfect knowledge on evolutionary dynamics with game transitions, we calculate the cooperation level for different noise levels in state perception (*n*_1_, *n*_2_) and compare the outcome with the complete knowledge setting (*n*_1_ = *n*_2_ = 0). **a** Under a small symmetric noise (*n*_1_ = *n*_2_ = 0.1), the cooperation level remains unchanged between imperfect and complete-knowledge cases. **b**,**c** A small asymmetric noise, dependent on the value of *n*_1_, *n*_2_, can either promote (**b**) or have a detrimental effect (**c**) on cooperation, when compared with the complete-knowledge setting. **d-f** To further explore the impact of imperfect knowledge, we numerically compute the fixation probability of different cooperative strategies when individuals misperceive the true state due to noise. Across all values of *n*_1_ and *n*_2_ considered, only strategy *CC* is favored by selection. Dots show simulation results, and the vertical dashed lines show the analytical prediction for strategy *CC* to be favored by selection. In each simulation, all edges are in state 1 initially. Other parameter values used are the same as in Fig. 2.

To understand these patterns, we numerically compute fixation probabilities of different cooperative strategies under imperfect state knowledge (Fig. 5d–f). Across all levels of informational noise, unconditional cooperators (*CC*) remain the dominant strategy and are therefore the primary drivers of cooperation in the imperfect-knowledge regime. As shown in Fig. 5e, when knowledge about the more beneficial state is more accurate than that about the less beneficial state, the threshold required for the success of *CC*-strategy becomes less restrictive, with 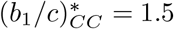. This reduced threshold (compared to the complete knowledge case, where 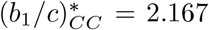 accounts for the higher level of cooperation observed under asymmetric state-information noise (*n*_1_ = 0.01, *n*_2_ = 0.1).

Complementing these numerical results, we analytically derive the condition under which the *CC*-strategy is favored by selection in the presence of imperfect state knowledge (**Supplementary Information** section 4). This analysis reveals that the beneficial effect of noise on cooperation under the transition vector **Q**_**1**_ becomes increasingly pronounced on graphs with large degree *k*. In particular, a small asymmetric noise, *n*_1_ *< n*_2_, can substantially lower the barrier for the success of *CC*. For example, when *k* = 100, and Δ*b* = 1, *c* = 1, the critical benefit-to-cost ratio, 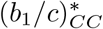, decreases from 50.17 in the complete knowledge case to 3.88 for *n*_1_ = 0.01, *n*_2_ = 0.1. These results demonstrate that imperfect environmental knowledge can fundamentally alter evolutionary outcomes. In particular, asymmetric noise in state perception can act as a mechanism that promotes cooperation, especially in highly connected networks, highlighting a non-trivial interplay between environmental feedback and the accuracy of state knowledge.

## Discussion

Our results show that the effect of environmental knowledge on cooperation is highly sensitive to the interplay between behavioral strategies and environmental feedback. When knowledge about the environment is lacking, environmental feedback, manifest as behavior-induced transitions between states, is not always beneficial for the spread of cooperation. Even in the minimal setting considered here, we find that access to state information can promote, have no effect on, or hinder cooperation. Which of these outcomes arises depends on the structure of the transition rule linking behavior to environmental change. Beyond the limiting cases of complete knowledge and lack of it, we also consider the possibility of individuals misperceiving the environmental state, thereby rendering their strategies effectively stochastic. Such imperfections can qualitatively alter evolutionary outcomes. Asymmetric noise in state perception can reduce the threshold for cooperation relative to both complete knowledge and no knowledge scenarios, thereby suggesting that uncertainty about the environmental state can, under certain conditions, facilitate cooperation.

Our primary focus on deterministic transition rules allowed for a complete classification of all transition structures of a given complexity and made it possible to identify general patterns in the effect of environmental feedback and state knowledge. To assess the robustness of our findings, we also examined a small set of stochastic transition rules 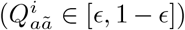 obtained by introducing noise (*ϵ*) in selected deterministic vectors. Across the three representative transition rules considered, our main results are largely robust to such perturbations. The exception is **Q**_**1**_ = (1, 0, 0; 1, 0, 0), which exhibits a higher sensitivity to stochasticity of state transitions than the others (**Supplementary** Fig. S5). These observations suggest that while deterministic transitions capture the essential mechanisms underlying the interplay between environmental feedback and information, stochastic transitions can introduce additional nuances in specific cases.

From a methodological perspective, we extend analytical approaches used for structured populations to incorporate environmental feedback with state-dependent behavior. Using pairwise and diffusion approximations under weak selection, we derive conditions for the success of cooperation (**Supplementary Information** sections 1, 2). We also formulate an alternative approach based on the sigma rule [55, 57] (**Supplementary Information** section 3), and where applicable, show that both methods yield consistent analytical results.

Our analysis shows that state knowledge can be profitable for individuals who exploit environmental shifts by switching to defection in the less beneficial state, provided subsequent environmental feedback facilitates a return to the more beneficial state. Then, such a conditional cooperator strategy (*CD*) can outperform unconditional defectors (*DD*) and is neutral with unconditional cooperators (*CC*) in state 1, thereby opening a pathway for the emergence of *CC* when *CD* is abundant. However, if environmental feedback strictly punishes any defection in state 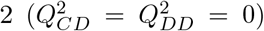, the advantage of the *CD* strategy disappears because of its inability to escape from the less beneficial state. This makes state-dependent behavior ineffective, leading to the appearance of a knowledge curse or neutral outcomes (**Supplementary** Fig. S2, S3).

The strategy space considered here is deliberately simple: individuals adopt pure strategies without memory, conditioning their behavior only on the current state. This setting may be particularly relevant for microbial systems embedded in structured environments such as biofilms, where interactions are local and behavioral responses are limited [22, 24, 58]. In such systems, individuals are unlikely to implement complex contingent strategies, and state-dependent but memory-less behavior may provide a natural description of their decision-making. At the same time, our framework can be interpreted as a limiting case of more complex strategies, allowing one to isolate the effect of environmental knowledge and feedback in structured populations.

Our framework can be generalized in several directions. First, we have assumed that the interaction network remains fixed while environmental states evolve. In many systems, however, interaction patterns may reorganize in response to environmental changes [59]. Extending the model to temporal or adaptive networks [60], where structure co-evolves with environmental states, may reveal additional feedback mechanisms. Second, in the present model, knowledge of the environmental state affects how individuals condition their actions, but does not expand the set of available actions. In more realistic settings, access to information may enable additional responses, such as rewarding or punishing behavior, depending on the state [61]. Third, extending the framework to networks with different topologies [62] would allow one to explore how structural heterogeneity interacts with environmental feedback and information.

Even though our results are closely parallel to those reported in stochastic games with memory-one strategies in well-mixed populations [47], there are important differences in the mechanisms by which cooperation evolves in these two scenarios. While the former relies on repeated interactions and complex memory-dependent strategies, the present work highlights how population structure interacts non-trivially with perceived environmental knowledge and feedback to shift the threshold for the evolution of cooperation. By demonstrating that similar qualitative behavior arises under both direct and network reciprocity, we provide a unified perspective on how environmental knowledge shapes the evolution of cooperation. Overall, our work shows that the effect of environmental information on cooperation is highly context dependent, even for simple memory-less strategies.

## Methods

### Game transitions with complete knowledge of the states

In our study, players can adopt pure state-dependent strategies when they are aware of the environmental states. For game transitions involving two states, the strategy space consists of four pure state-dependent strategies: **s**_**1**_ = (*C, C*), **s**_**2**_ = (*C, D*), **s**_**3**_ = (*D, C*), and **s**_**4**_ = (*D, D*). For the competition between *n*(*>* 2) strategies, selection can favor multiple strategies. In the mutation-selection process, a strategy **s** is favored by selection if it is more abundant than the average, 1*/n*, in the stationary distribution,

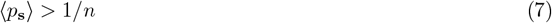

where *p*_**s**_ is the frequency of strategy **s**, and the angular brackets denote average over the stationary distribution. Without selection (*δ* = 0), all strategies have equal abundances 1*/n*; thus, for weak selection (*δ* ≪ 1), if a strategy **s** is more abundant than its neutral abundance, then one can say strategy **s** is favored by selection. In the limit of low mutation, the Eq. (7) translates as [55],

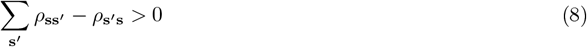

where, *ρ*_**ss**_*′* is the fixation probability of a strategy **s** in a population of **s**′ players. Equation 8 shows that under low mutation and weak selection, evolutionary dynamics in structured populations with multiple strategies can be fully characterized through the pairwise comparisons between fixation probabilities of strategies. Using Eq. (8), we calculate the analytical conditions for cooperative strategies **s**_1_, **s**_2_, and **s**_3_ to be favored by selection when individuals have complete knowledge of the state. The corresponding condition for each strategy can be written as follows:

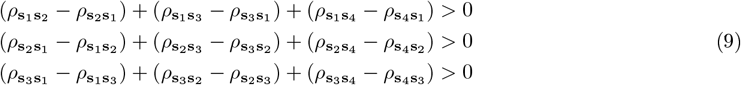

In the limit of low mutation, mutants (say, strategy **s**) arise only rarely in a resident population of strategy **s**′. When the mutant arises, it rapidly either goes to fixation or extinction in the population. The number of mutants in a finite population follows a one-step random walk. Under the diffusion approximation, this discrete stochastic dynamics can be approximated by a one-dimensional diffusion process in the mutant frequency, *p*_**s**_. At each time-step, the frequency of strategy **s** can either increases by 1*/N* (Δ*p*_**s**_ = 1*/N* ) or decreases by 1*/N* (Δ*p*_**s**_ = −1*/N* ). Thus, within the short time interval *t* → *t* + Δ*t*, the expected increment is,

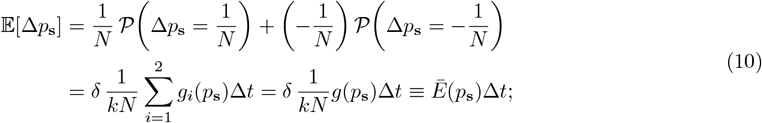

where, *g*_*i*_(*p*_**s**_) is a function of *p*_**s**_ and depends on the state *i*. The variance can be written as,

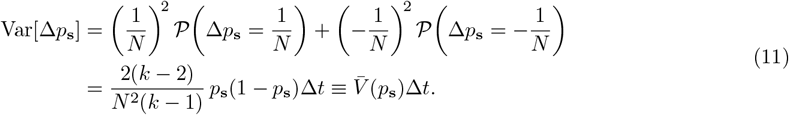

For the detailed calculation of *g*_*i*_(*p*_**s**_) and 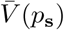, see **Supplementary Information**, section A.5. The fixation probability *ϕ*_**ss**_*′* (*y*) of strategy **s** in a resident population of **s**′−players, given an initial frequency *p*_**s**_(*t* = 0) = *y*, satisfies the backward Kolmogorov equation,

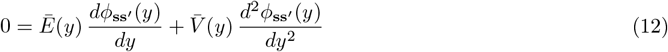

The solution of Eq. (12) is

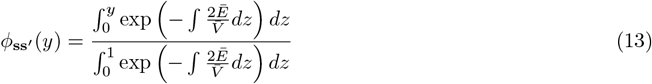

The fixation probability of *y* fraction of **s**′-player in a population of **s**-players is

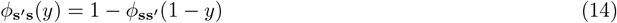

For sufficiently small *y* and *δ* ≪ 1 we have (for detailed calculation, see section 1 A.5 of the **Supplementary Information**),

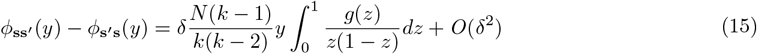

For a sufficiently large population and *y* = 1*/N*, we have

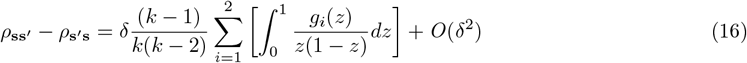

To calculate each condition in Eq. (9), we compute the difference of fixation probabilities *ρ*_**ss**_*′* − *ρ*_**s**_*′*_**s**_ for each pair strategies **s, s**′ ∈ { **s**_1_, **s**_2_, **s**_3_, **s**_4_ } using Eq. (16). See section 1A of the **Supplementary Information** for the detailed calculation of the difference in fixation probabilities for each pair of strategies under different game transition rules.

#### Calculation of *α*_1_ and *α*_2_

When state-dependent strategies (*CD* or *DC*) get fixed in the population, their contribution to the cooperation rate depends on the fraction of edges (*α*_1_ or *α*_2_) that remain in state (1 or 2) after equilibration. We therefore ran the simulation for an additional 5000 time steps after fixation and determined the fraction of edges in state 1 (state 2) when strategy *CD* (*DC*) takes over the population to obtain the probabilities *α*_1_ (*α*_2_).

### Game transitions with no knowledge of the states

When players are unaware of the environmental state, they adopt pure state-independent strategies. There are two pure state-independent strategies: **s**_**1**_ = (*C, C*), and **s**_**4**_ = (*D, D*). The set of state-independent strategies is therefore a subset of the state-dependent strategies. In the main text, in the setting without state knowledge, we denote the strategy (*C, C*) as strategy *C* and the strategy (*D, D*) as strategy *D*. The condition for the success of *C* over *D* is calculated using Eq. (16) as,

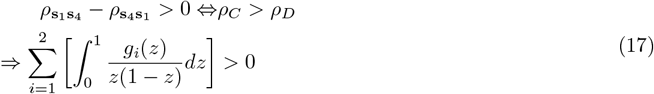

Here, we need to calculate the function *g*_*i*_(*z*) in each state *i* for the pair of strategies **s**_1_ and **s**_4_, and by substituting the corresponding *g*_*i*_(*z*) for each values of *i* in Eq. (17), we get the condition for strategy *C* to be favored by selection over *D*.

### Game transitions with imperfect knowledge of the states

In our framework, when players lack knowledge or have complete knowledge of the environmental states, they adopt pure strategies, cooperating or defecting with probability 1 in each state, according to their strategy. However, with imperfect state knowledge, a player misperceives her true current state *j* with a probability *n*_*j*_, which consequently can modify the player’s effective strategy. In this scenario, players use stochastic strategies to cooperate in each state.

Let 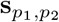 denote a state-dependent strategy with which, at each time step, a player cooperates with probability *p*_1_ in state 1 and with probability *p*_2_ in state 2. **s**_1,1_ thus corresponds to *CC*-strategy, **s**_1,0_ corresponds to *CD*-strategy, **s**_0,1_ corresponds to *DC*-strategy and **s**_0,0_ corresponds to *DD*-strategy. Due to noise in state perception, an individual with imperfect knowledge of the states cooperates with probability

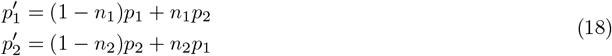

Thus, when individuals have imperfect knowledge of states, the 4 pure state-dependent strategies are effectively modified as follows: 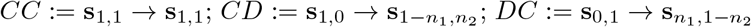 and *DD* := **s**_0,0_ **s**_0,0_.

In this scenario, the expected level of cooperation is defined as,

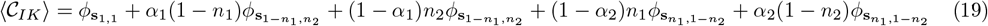

To understand the expression above, let’s consider the second term on the right-hand side. Here, *α*_1_ denotes the probability that an edge equilibrates in state 1 after strategy 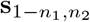 has taken over the population. The factor (1 − *n*_1_) represents the probability that an individual with strategy 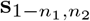 cooperates when in state 1, while 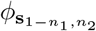 denotes the fixation probability of this strategy starting from an unbiased initial population state (1*/*4, 1*/*4, 1*/*4, 1*/*4). Similarly, all other terms can be explained as well. We use Eq. (19) to calculate the expected cooperation level in the imperfect knowledge scenario in Fig. 5.

## Acknowledgement

AB acknowledges the Kepler Computing Facility maintained by the Department of Physical Sciences at IISER Kolkata, and partial support from QUT Brisbane for a research visit during the preparation of this manuscript. MK acknowledges support from Australian Research Council Special Research Initiative, SRIEAS Grant SR200100005, Securing Antarctica’s Environmental Future, and Australian Research Council Discovery Early Career Researcher Award DE250101223. We thank Sagar Chakraborty, Nadiah Kristensen, Greg Kubitz, and Casper van Elteren for helpful discussions.

## Author contributions

A.B.: conceptualization, data curation, formal analysis, investigation, methodology, software, visualization, writing-original draft, writing-review and editing; M.K.: formal analysis, methodology, writing-original draft, writing-review and editing; S.S.: formal analysis, methodology, resources, supervision, writing-original draft, writing-review and editing.

## Supplementary Information

### Supplementary Information Text

We study game transitions between two states, where each state is represented by a game with a distinct payoff matrix. The payoff matrices associated with game 1 (**A**^(1)^) and game 2 (**A**^(2)^) are, respectively,

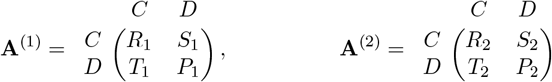

where each entry of the payoff matrix **A**^(*i*)^ corresponds to the payoff obtained by a player choosing the action indicated in the column when interacting with a co-player choosing the action indicated in the row of game *i* ∈ {1, 2}. For example, if a player cooperates (*C*) and her co-player defects (*D*) in game 1, the player gets a payoff 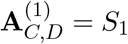 and her co-player receives a payoff 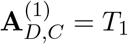.

Each individual is represented by a node on a regular graph of degree *k*, where the number of nodes is *N*, and the number of undirected edges is *kN/*2. An edge between two nodes denotes a one-shot game interaction between two individuals. The game transitions pattern on an edge is described by a transition vector,

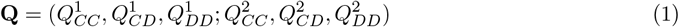

where each entry 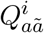 represents the probability that an edge transitions to state (game) 1 in the next time step. The probability depends on the actions *a* and *ã* of players connected by that edge in game *i* ∈ {1, 2} in the previous time-step. Here we assume, 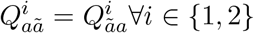. Due to game transitions across all edges, here, games played with different neighbors can be distinct. At each time step, each player interacts with all their neighbors and derives an accumulated payoff *π* from all interactions. This payoff is then interpreted as reproductive fitness, *f* = 1 − *δ* + *δπ*, where *δ* ≥ 0 is the selection strength. We study the effect of weak selection (*δ* ≪ 1) [1], which has been widely studied in evolutionary biology [2].

#### 1 Game transitions with complete knowledge of the states

In this study, when players have complete knowledge of the state they are currently in, they can adopt state-dependent strategies. When players adopt state-dependent strategies, their actions in different states (games) can be distinct. In our model, at each evolutionary time step, players interact only once; thus, their strategy does not take into account any past actions. For game transitions between 2 states, a state-dependent strategy **s**_**d**_ can be represented as **s**_**d**_ = (*a*_1_, *a*_2_), where *a*_1_, *a*_2_ ∈ { *C, D* } are actions in game 1 and game 2, respectively, of a player adopting strategy **s**_**d**_. Note that here we assume, according to a player’s strategy, they take an action *a*_*i*_ in game *i* with probability 1; such deterministic strategies are referred to as “pure strategies”. Since players can take either of the two actions *C* or *D* in each game, the set of pure state-dependent strategies consists of 4 strategies only,

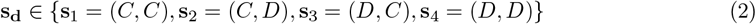

For simplicity, we refer to the **s**_1_ = (*C, C*) strategy as *CC* − strategy only and so on for all other strategies, for the rest of this article.

For the competition between *n*(*>* 2) strategies, selection can favor multiple strategies. Tarnita et al. [3][4] have shown that a strategy **s** is favored by selection if it is more abundant than the average, 1*/n*, in the stationary distribution,

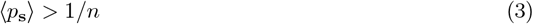

where *p*_**s**_ is the frequency of strategy **s**, and the angular brackets denote average over the stationary distribution. Without selection (*δ* = 0), all strategies have equal abundances 1*/n*; thus, for weak selection (*δ* ≪ 1), if a strategy **s** is more abundant than its neutral abundance, then one can say strategy **s** is favored by selection. In the limit of low mutation, the Eq. 3 translates as [4],

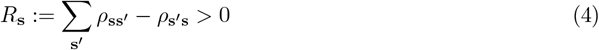

where, *ρ*_**ss**_*′* is the fixation probability of a strategy **s** in a population of **s**′ players. Eq. 4 implies that to study game transitions with complete knowledge of states (**s, s**′ ∈ { **s**_1_, **s**_2_, **s**_3_, **s**_4_ }) with several strategies (*n* = 4), it suffices to analyze ^4^*C*_2_ = 6 pairwise competition between strategies. Here, we want to derive the analytical conditions for cooperative strategies **s**_1_ = *CC*, **s**_2_ = *CD*, and **s**_3_ = *DC* to be favored by selection when individuals have complete knowledge of the state. Using Eq. 4, the conditions for each strategy to be favored by selection can be written as,

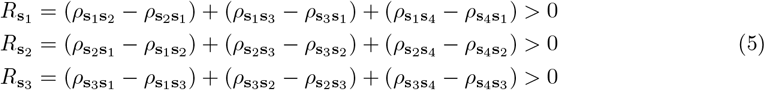

In the following, using a combination of pair approximation[5] and diffusion approximation[6], we systematically study all 6 pairwise competitions between strategies (*ρ*_**ss**_*′* − *ρ*_**s**_*′*_**s**_) to obtain the analytical condition for each of the three cases expressed in Eq. 5.

Again, we stress that there are *n* = 4 competing strategies in our main system with complete knowledge; however, due to the assumption of low mutation, the population spends most of its time in a homogeneous state. With the rare occurrence of a mutant, it either goes to fixation or gets extinct. Thus, in the limit of low mutation, we analytically calculate the pairwise fixation probabilities of different strategies to describe the whole system with multiple strategies.

Note that, in this paper, we have studied state-dependent strategies for game transitions between 2 states; however, the methodology developed here can easily be extended to game transitions between any *m* states. For game transitions between *m* states, a state-dependent strategy is written as **s** = (*a*_1_, *a*_2_, …, *a*_*m*_), where each *a*_*i*_ ∈ { *C, D* }, yielding *n* = 2^*m*^ competing state-dependent strategies in the population. For this general setup, one must study ^*n*^*C*_2_ pairwise competition between strategies in order to determine the selection conditions under which each state-dependent strategy is favored.

Below, we develop a general formalism to study pairwise competition between any two state-dependent strategies, say **s** and **s**′ (**s, s**′ ∈ {**s**_1_, **s**_2_, **s**_3_, **s**_4_}) for game transitions between 2 states. Each strategy is represented by a pair of actions: **s** = (*a*_1_, *a*_2_) and 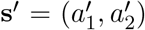. A player using strategy **s** chooses action *a*_1_ in game 1 and *a*_2_ in game 2, whereas a player using strategy **s**′ chooses 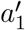 in game 1 and 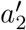 in game 2. We now introduce the following variables to characterize the evolving system:

*p*_**s**_: the frequency of **s**-players;

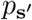 frequency of s_′_players;

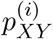: the frequency of edges in game *i* connecting two players using strategy *X* and *Y* ;

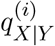 : the conditional probability to find a *X*-player given the adjacent node is occupied by a *Y* -player and the edge connecting these two players is in game *i*;

*p*_*XY*_ : the frequency of edges that connect a *X*-player and a *Y* -player;

*q*_*X*|*Y*_ : the conditional probability to find a *X*-player given that the adjacent node is occupied by a *Y* -player.

Here, both *X* and *Y* stand for **s** and **s**′ (*X, Y* ∈ { **s, s**′ }) and **s, s**′ are an arbitrary pair of state-dependent strategies defined by the action pairs, where, **s, s**′ ∈ { **s**_1_, **s**_2_, **s**_3_, **s**_4_ }. Then, we have the following constraints on our system,

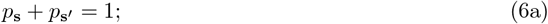

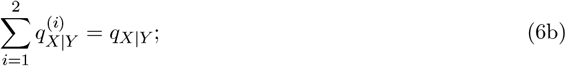

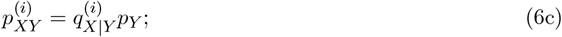

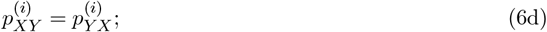

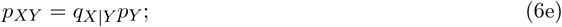

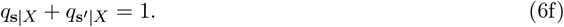

which imply that the dynamics between two strategies, **s** and **s**′, can be described using variables *p*_**s**_ and 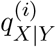 in pair approximation [7, 8].

#### A Death-Birth update

After all interactions and payoff accumulations, we use a death-birth update process where a player is selected for death at random and all the neighboring nodes compete to fill the vacant site with probability proportional to fitness [8]. Note that, in our model, selection acts only on strategies, but both players’ strategies and the game they play coevolve over time. Below, we start by investigating the change in the frequency of **s**−player.

##### A.1 Change in *p*_s_: Updating a s′-player

A player using strategy **s**′ is chosen randomly with probability *p*_**s**_*′* . It’s *k* neighbors compete for the empty site. Among *k* neighbors, let 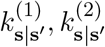 denote the number of neighbors, with strategy **s**, who play game 1, game 2, respectively, with the focal **s**′−player. Similarly, 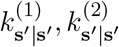 denotes the number of neighbors with strategy s′ who play game 1, game 2, respectively, with the focal **s**′ − player. The neighborhood configuration of the focal player can be represented as 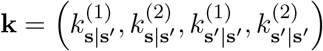, which satisfies the condition, 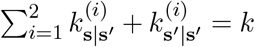. The probability of such a neighborhood configuration **k** around the focal player is,

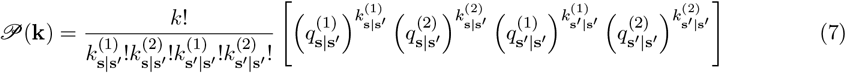

The fitness of each **s**-player who plays game *i* with the focal **s**′-player,

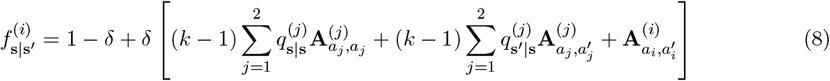

In the calculation of the fitness of an **s**−player who is a neighbor to the focal **s**′−player in game *i*, the **s**−player certainly receives a payoff 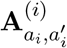, since the **s**−player uses action *a*_*i*_ and **s**′−player uses action 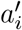 in game *i*. For the remaining *k* − 1 uncertain neighbors, **s** −player receives a payoff according to the conditional probabilities of finding other players (either **s** − or **s**′ − player) around the **s** − player and actions of players in each game *j*.

Similarly, the fitness of each **s**′-player who plays game *i* with the focal player,

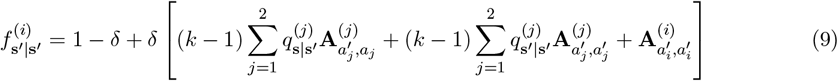

The probability that one of the **s**−players takes over the vacant site is,

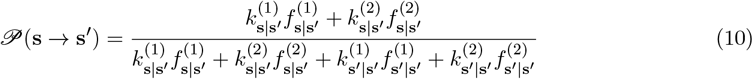

The probability that one of the **s**′−players takes over the vacant site is,

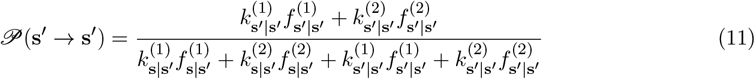

Therefore, *p*_**s**_ increases by 1*/N* with probability,

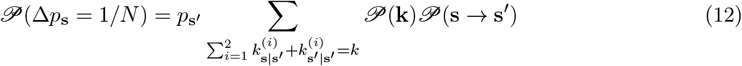

##### A.2 Change in *p*_s_: Updating a s−player

A player using strategy **s** is chosen randomly with probability *p*_**s**_. It’s *k* neighbors compete for the empty site. Among *k* neighbors, let 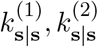 denote the number of neighbors who adopt strategy **s** and play game 1, game 2, respectively, with the focal **s**−player. Similarly, 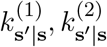 denotes the number of neighbors who adopt strategy s′ and play game 1, game 2, respectively, with the focal **s**− player. The neighborhood configuration around the focal **s**−player can be represented as 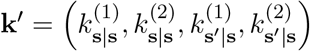 which satisfies the condition, 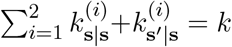. The probability of such a neighborhood configuration **k**′ around the focal player is,

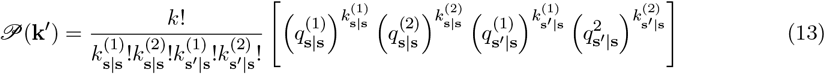

The fitness of each **s**-player who plays game *i* with the focal **s**-player,

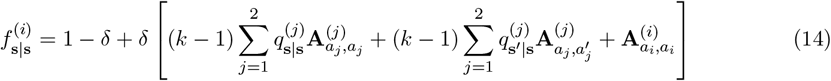

The fitness of each **s**′-player who plays game *i* with the focal player,

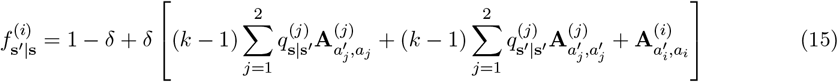

The probability that one of the **s**−players takes over the vacant site is,

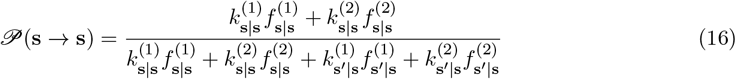

The probability that one of the **s**′−players takes over the vacant site is,

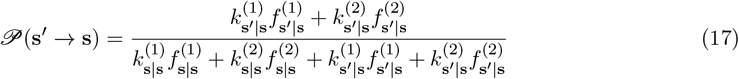

Therefore, *p*_**s**_ decreases by 1*/N* with probability,

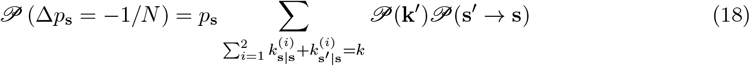

##### A.3 Change in *p*_**s**_

If one replacement occurs in one unit of time, at each time step, the change in the frequency of players using strategy **s** takes into account the probability of increase and decrease in frequency of the **s** − player by 1*/N* . Using Eqs. 12 and 18, we have,

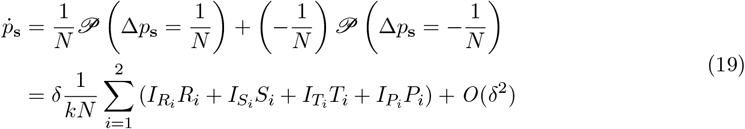

where the coefficients 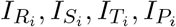 depends on the pair of strategies **s** and **s**′ considered. In sections A.6-A.11, we will obtain the expression for each coefficient for competition between all 6 pairs of strategies.

##### A.4 Edge dynamics

We know from Eqs. 6, the competition between two strategies can be described by *p*_**s**_ and 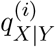 . So far, we have studied the dynamics of *p*_**s**_. Below, we study the dynamics concerning 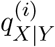.

###### A.4.1 Change of ss−edges in game 1

When a random player *m* is chosen to die, the type of edge between *m* and its immediate neighbors changes. However, 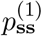 can change in the remaining edges as well, since at each time step, all games across all edges update, potentially changing the edge type. Note that, change in 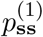 is different from the change in *p*_**s**_. When an **s**−player dies and is replaced by an **s**−player or a **s**′−player dies and is replaced by a **s**′−player, *p*_**s**_ doesn’t change; however, 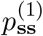 can change due to game transitions along the edges. Change in 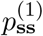 has two parts: (i) switching of edges connecting the focal (dead) player and its immediate neighbors to a different game state and (ii) switching of all other edges to a different game state. Let’s first consider the changes due to the first part.

###### Increase of ss−edges in game 1

We first consider a case in which a random **s**′−player is chosen to die. We take the neighborhood configuration, 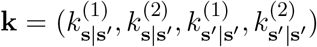. The number of **ss**-edges in game 1 increases by 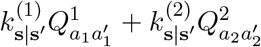, when a **s**′−player is replaced by an **s**-player. The first part corresponds to the case where the new **s**-player connects to the previous **s** − player’s neighbors who were playing game 1 using strategy **s**, and in the current time step, these edges remain in game 1 with probability 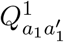, (since, **s**−player uses action *a*_1_ and **s**′−player uses action 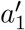 in game 1), thus increasing the number of **ss**−edges in game 1 by 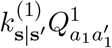 . Similarly, the second part corresponds to the case where the new **s**-player connects to the dead **s**′−player’s neighbors **s** in game 2 and in the current time step, these edges transition to game 1 with probability 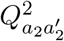, (since, **s**−player uses action *a*_2_ and the dead **s**′−player uses action 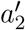 in game 2), thus increasing the number of **ss**−edges in game 1 by 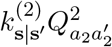. Since the total number of undirected edges in the system is *kN/*2, the frequency of **ss**−edges in game 1, 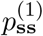 is increased by 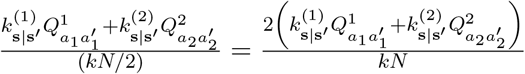 when a random **s**′ − player is chosen to die. For the neighborhood configuration **k**, the probability of the above event is,

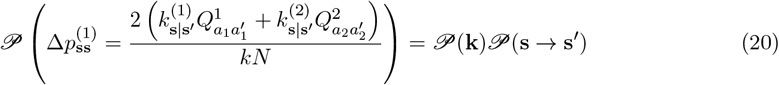

Next, we consider that a random **s**−player is chosen to die. We take the neighborhood configuration, 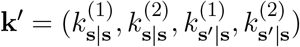. The number of **ss**−edges in game 1 increases by 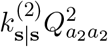 when an **s**−player is replaced by another **s**-player−the new **s**-player connects to previous **s**−player’s neighbors **s** who were playing game 2, and in the current time step they transition to game 1 with probability 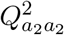 (since, both players using strategy **s** takes action *a*_2_ in game 2), thus increasing the number of **ss**−edges by 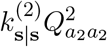. For the neighbor configuration **k**′, the probability of this event,

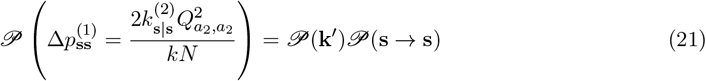

###### Decrease of ss−edges in game 1

When a random **s**−player dies and is replaced by another **s**−player, the number of **ss**−edges can decrease due to the transition of **ss**−edges from game 1 in the previous time step to game 2 in the current time step with probability 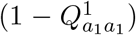. Consequently, decreasing 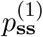 by 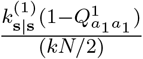 . For the neighbor configuration **k**′, the probability of this event,

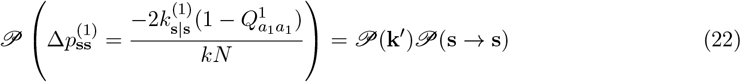

The number of **ss**-edges in game 1 decreases by 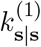 when the focal **s**-player is replaced by a **s**′-player. The dead **s** -player disconnects from the previous neighbors **s** who were playing game 1 in the previous time step, thus decreasing **ss**−edges in game 1 by 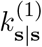. For the neighborhood configuration **k**′, the probability of this event,

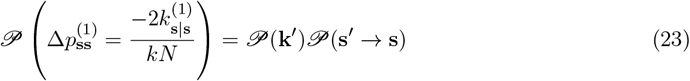

###### Increase of ss−edges in game 1

Let’s first consider that a random **s**′−player is chosen to die. In this case, regardless of which player replaces the focal **s**′−player, the number of **ss**−edges in game 1 can increase due to the transition of all **ss**−edges in game 2 in the previous time step to game 1 in the current time step with probability 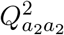 (since, players adopting strategy **s** uses action *a*_2_ in game 2). Thus, the number of **ss**−edges increases by 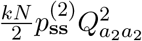. For the same neighborhood configuration around **s**′−player, 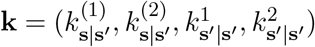, the probability of the above event is,

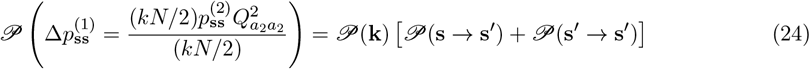

Next, we consider that an **s**−player is randomly chosen to die. The number of all **ss**−edges in game 2 in the previous time step is 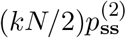. Among these _(_**ss**−edges, the num_)_ber of edges in game 2 outside the nearest neighborhood of the focal **s**−player is 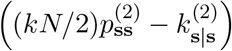. Irrespective of which player replaces the focal **s**−player, these edges transition to game 1 with probability 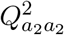. For neighborhood configuration **k**′, the probability of this event,

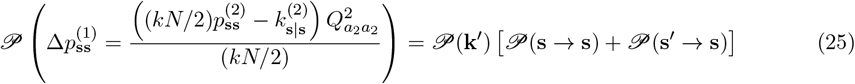

###### Decrease of ss−edges in game 1

Similar to the above, when a random **s**′−player is chosen to die, the frequency of **ss**−edges decreases due to the switching of **ss**−edges from game 1 to game 2. The number of **ss**−edges decreases by 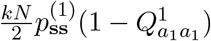. For the same neighborhood configuration **k** around **s**′−player, the probability of the above event is,

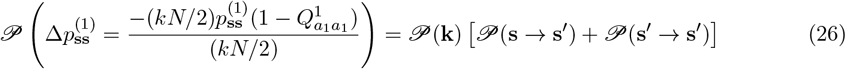

When an **s**−player is chosen to die, the numbe(r of **ss**−edges decr)eases due to switching of all other edges except in the immediate neighborhood is 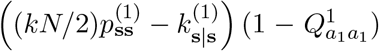 For neighborhood configuration **k**′ around **s**−player, the probability of this event,

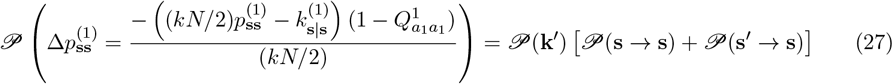

Using Eqs 20-27 the time derivative of 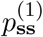 can be written as,

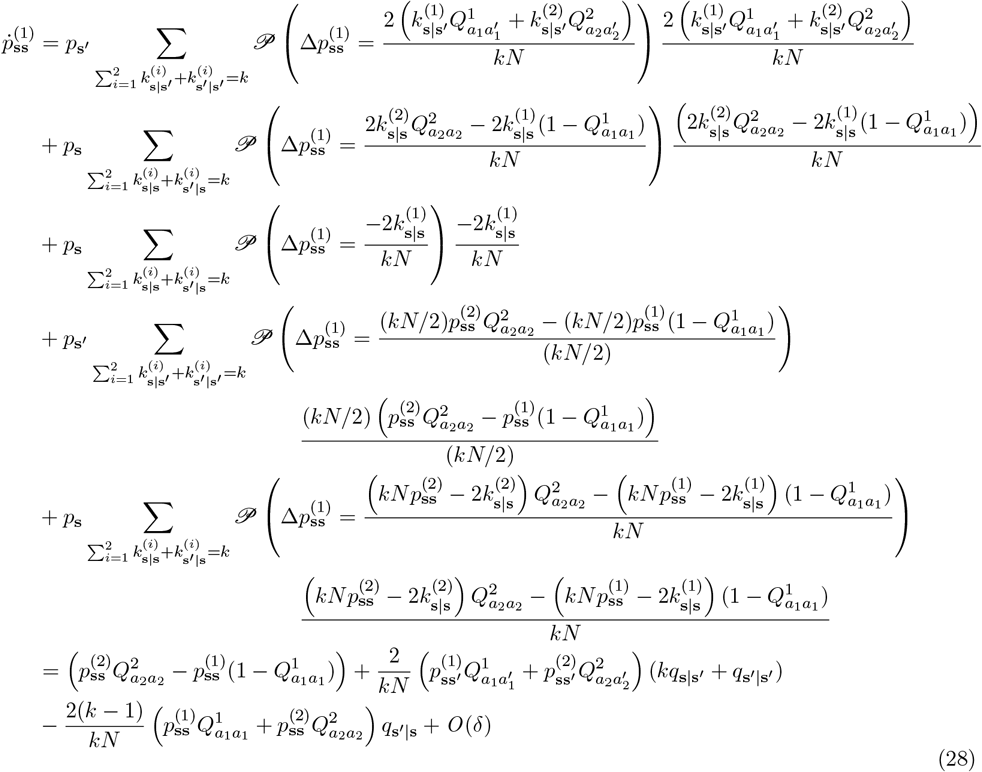

##### A.4.2 Change of ss−edges in game 2

Similar to changes of **ss** − edges in game 2, the change in 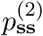 has two parts: (i) Inter-game switching of edges connecting the focal player and its immediate neighbors and (ii) Inter-game switching of all the other edges. Let’s first consider the changes due to the first part.

###### Increase of ss−edges in game 2

When a **s**′−player is selected to die, and its neighborhood configuration is **k**, the increase in 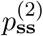 due to edges between the focal **s**′−player and it’s nearest neighbors is

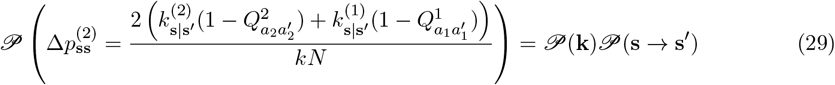

Note that here the probability of an **ss**′−edge to remain in game state 2 is 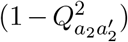 (since, **s**−player uses action (*a*_2_) and the **s**′−player uses action 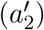 in game 2), and probability to transition to game state 2 from game 1 is 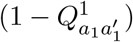.

When an **s**−player dies and it’s neighborhood configuration is **k**′, the increase in 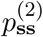 due to edges between the focal **s**−player and it’s nearest neighbors is,

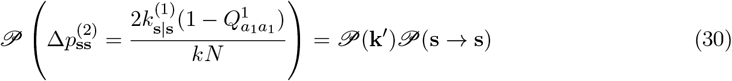

###### Decrease of ss−edges in game 2

When an **s**−player is randomly chosen to die and is replaced by another **s**−player, for the same neighborhood configuration **k**′, the decrease in 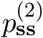 due to switching of edges between the focal **s**−player and it’s nearest neighbors is,

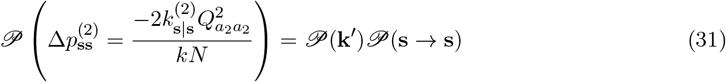

When the focal (dead) **s**−player is replaced by a **s**′−player, for the neighborhood configuration **k**, the decrease in 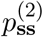 due to switching of edges between the focal **s**−player and it’s nearest neighbors is,

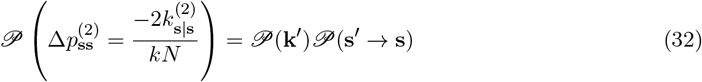

###### Increase of ss−edges in game 2

When a random **s**′−player is chosen to die, and its neighborhood configuration is **k**, the increase in 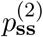 due to switching of all other edges except in the immediate neighborhood of the focal player is,

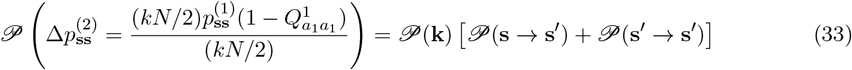

When a random **s**−player dies, and its neighborhood configuration is **k**′, the increase in 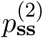 due to switching of all other edges except in the immediate neighborhood of the focal player is,

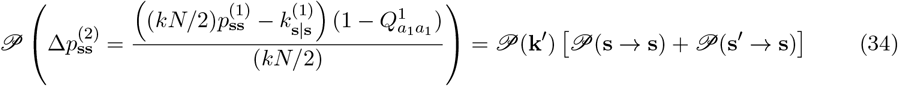

###### Decrease of ss−edges in game 2

When a random **s**′−player is chosen to die, and its neighborhood configuration is **k**, the decrease in 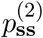 due to switching of all other edges except in the immediate neighborhood of the focal player is,

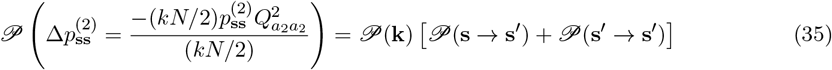

When a random **s**−player dies, and its neighborhood configuration is **k**′, the decrease in 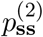 due to switching of all other edges except in the immediate neighborhood of the focal player is,

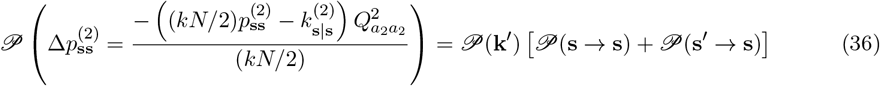

Analogously, using Eqs 29-36 we obtain the time derivative of 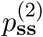

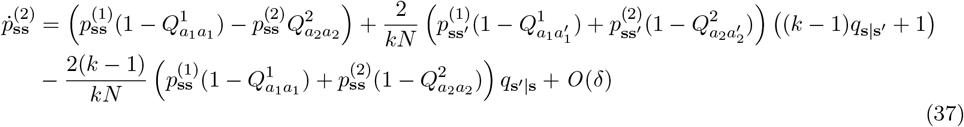

##### A.4.3 Separation of different time scales

Using Eqs 28 and 37 we can write,

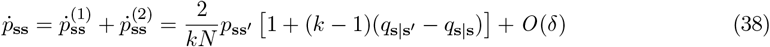

and using, *q*_**s**|**s**_ = *p*_**ss**_*/p*_**s**_ we can write 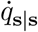 in terms of 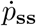 as follows,

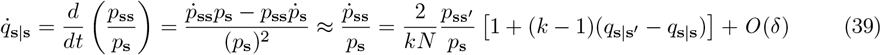

Note that due to weak selection limit (*δ* ≪ 1), in Eq. 39 we considered only the leading order term in the Taylor expansion (which is 𝒪(*δ*^0^), which is non-zero, and ignored all other higher order terms. In the above calculation, we have also considered 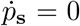, as the first non-zero term in the expression of 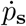 (Eq. 19) is of 𝒪(*δ*^1^), which has been ignored while calculating Eq. 39.

Comparing Eq. 19 and Eq. 39, we observe that a change in *p*_**s**_ happens at 𝒪(*δ*^1^), while changes in *q*_**s**|**s**_ happen at 𝒪(*δ*^0^). Thus, for weak selection (*δ* ≪ 1), *q*_**s**|**s**_ equilibrates on a faster time-scale than *p*_**s**_. This allows us to solve 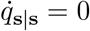 to get the equilibrium value of 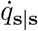 in terms of *p*_**s**_ which then becomes the only independent variable governing the evolutionary dynamics of the system.

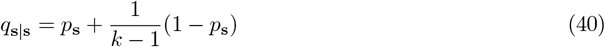

Using Eq. 6a-6f and Eq. 40, we see that *p*_**ss**_*′, q*_**s**|**s**_, *q*_**s**_*′*_|**s**_, *q*_**s**|**s**_*′, q*_**s**_*′*_|**s**_*′* can be written in terms of *p*_**s**_ as follows,

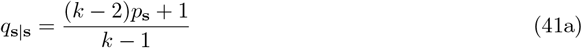

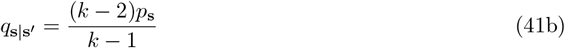

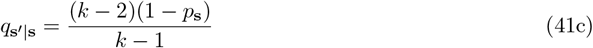

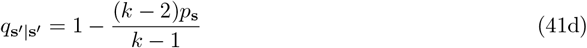

Note that the identity in Eq. 40 and Eqs. 41a-41d is true for all pairs of state-dependent strategies **s** and **s**′ ∈ {**s**_1_, **s**_2_, **s**_3_, **s**_4_}.

##### A.4.4 Change of ss′ − edges in game 1 due to switching of edges connecting the focal player and nearest neighbors

Here, we first calculate the net change in **ss**′−edges when a **s**′−player is randomly chosen to die. For the neighborhood configuration **k** (as considered earlier) around the **s**′−player, the probability that the configuration **k** is found around the **s**′−player and the net change in **ss**′−edges when an **s**−player replaces the **s**′−player

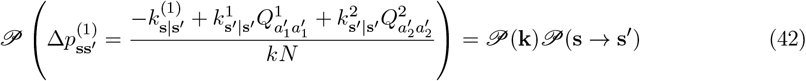

and when a **s**′−player replace the focal **s**′−player

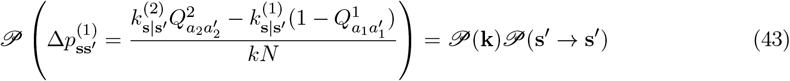

When an **s**−player is chosen to die, the probability of finding the neighborhood configuration **k**′ around the **s**−player and the change in **ss**′−edges in game 1 due to the event of an **s**−player replacing the focal **s**−player

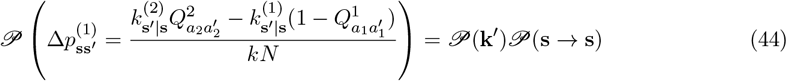

and **s**′−player replacing the focal **s**−player

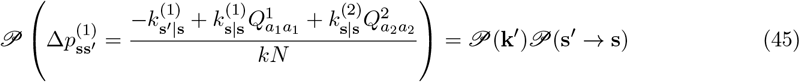

Note that using Eq. 6d, here we are considering 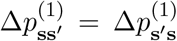, thus each **ss**′−edge is effectively counted in both directions. To avoid double-counting, the change in the number of **ss**′ − edges is therefore normalized by *kN* (the total number of ordered half-edges) rather than by *kN/*2, which counts undirected edges only once.

###### Change of ss′−edges in game 1 due to switching of all other edges

When a **s**′−player is randomly chosen to die, under the neighborhood configuration **k** around the focal (dead) player, the change in **ss**′−edges in game 1 due to switching of all other edges except the edges connecting the focal **s**′−player and its nearest neighbors is

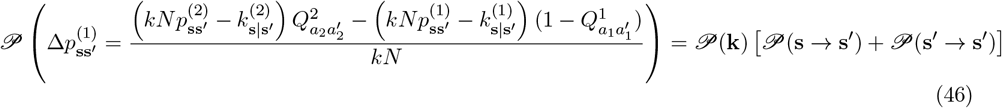

When an **s**−player is randomly chosen to die, under the neighborhood configuration **k**′ around the focal (dead) player, the change in **ss**′−edges in game 1 due to switching of all other edges except the edges connecting the focal **s**−player and its nearest neighbors is

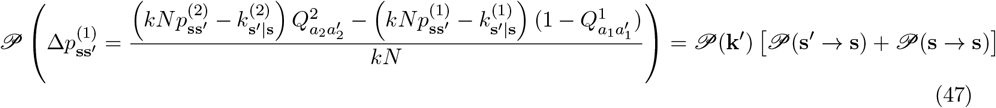

Analogously, using 42-47 we have the time derivative of 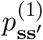

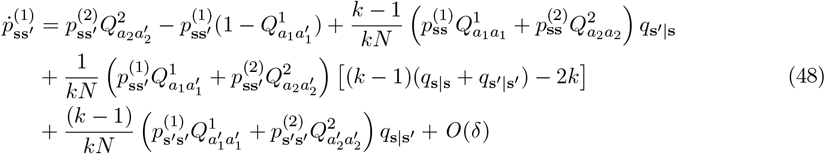

##### A.4.5 Change of s′s′ − edges in game 1 due to switching of edges connecting the focal player and nearest neighbors

We first consider the net change in **s**′**s**′−edges when a **s**′−player is randomly chosen to die. For the neighborhood configuration **k** (as considered earlier) around the **s**′−player, the probability that the configuration **k** is found around the **s**′−player and the net change in **s**′**s**′−edges when an **s**−player replace the focal **s**′−player

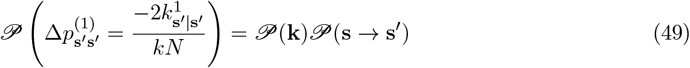

and when a **s**′−player replace the focal **s**′−player

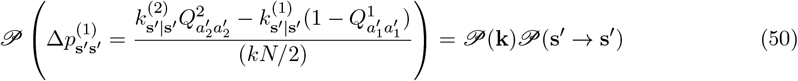

When an **s**−player is chosen to die, the probability of finding the neighborhood configuration **k**′ around the **s**−player and the change in **s**′**s**′−edges in game 1 due to the event of a **s**′−player replacing the focal **s**−player

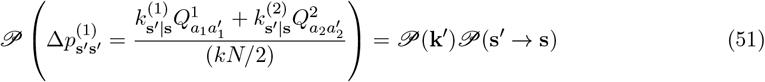

Note that when the randomly chosen focal (dead) player is an **s** − player, there is no **s**′**s**′ − edge in the neighborhood of the focal player. Thus, when an **s** − player replaces the focal **s** − player, there is no change in the number of **s**′**s**′ − edges in game 1 due to the switching of edges connecting the focal player and nearest neighbors.

###### Change of s′s′−edges in game 1 due to switching of all other edges

When a **s**′−player is randomly chosen to die, under the neighborhood configuration **k** around the focal (dead) player, the change in **s**′**s**′−edges in game 1 due to switching of all other edges except the edges connecting the focal **s**′−player and its nearest neighbors is

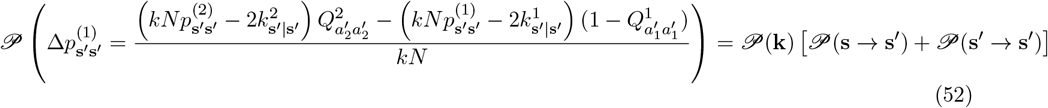

When an **s**−player is randomly chosen to die, under the neighborhood configuration **k**′ around the focal (dead) player, the change in **s**′**s**′−edges in game 1 due to switching of all other edges except the edges connecting the focal **s**−player and its nearest neighbors is

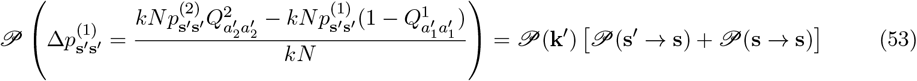

Analogously, using 49-53 we have the time derivative of 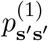

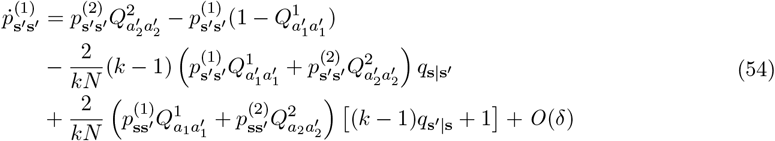

Using Eqs. 41a-41d and 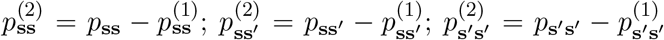, we rewrite Eqs. 28,48, and 54 as following,

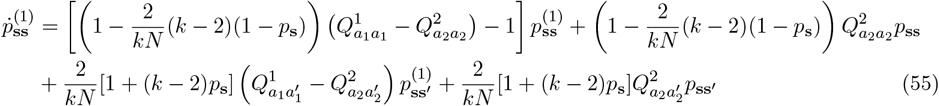

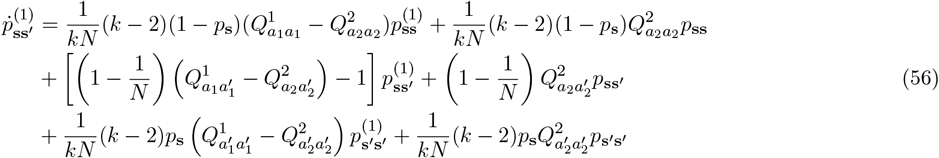

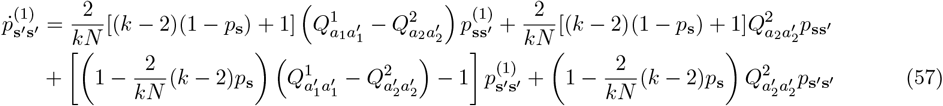

The edge dynamics in game 1, described by Eqs. 55,56, and 57 can be rewritten in a matrix form as following,

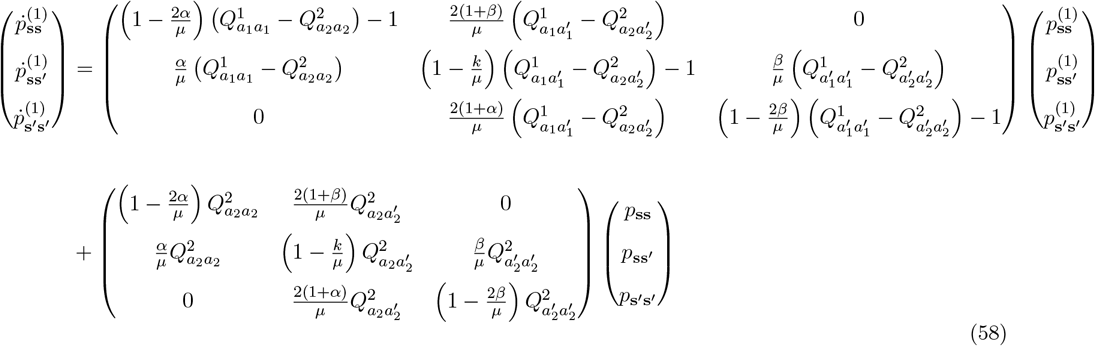

where, *α* = (*k* − 2)(1 − *p*_**s**_), *β* = (*k* − 2)*p*_**s**_, and *µ* = *kN* . Let’s denote two column vectors as, 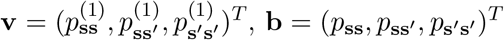 and **I** is a 3 × 3 identity matrix. Eq. 58 can be written in the vector form

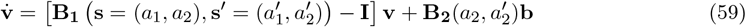

where,

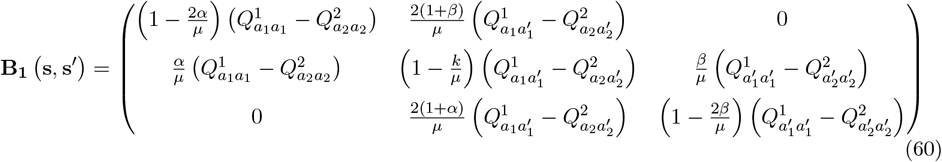

and

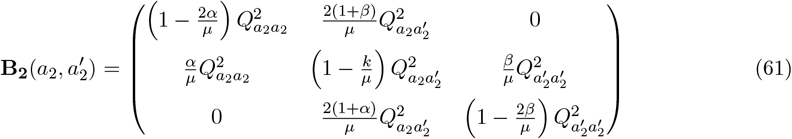

The solution of the linear system described by Eq. 59 is given by

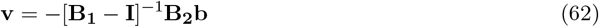

For 0 *< p*_**s**_ *<* 1, Eq. 59 admits a unique equilibrium solution given by Eq. 62, and the system always converges to the equilibrium point irrespective of the initial game individuals are playing. However, if the det(**B**_**1**_ − **I**) = 0 then the solution given by Eq. 62 doesn’t exist. In such cases, the system can admit multiple solutions, and the initial conditions determine the final outcome. A systematic analysis of all 64 deterministic transition vectors across all strategy pairs shows that the system of Eq. 59 is sensitive to initial conditions only under those transition vectors that have one absorbing state (15 cases). Here, once either game state is reached, it cannot be left, as 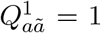 or 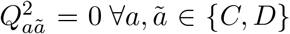. Within our framework of game transitions, these transition vectors are not meaningful as they don’t allow transitions between states, and individuals play only one of the two games indefinitely. Consequently, state-dependent behavior under complete knowledge is also irrelevant for these 15 deterministic transition vectors. For all remaining deterministic transition structures, the solution given by Eq. 62 always exists for all pairs of strategies.

Eq. 62 shows that 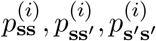, for each *i* ∈ { 1, 2 } can be expressed in terms of *p*_**ss**_, *p*_**ss**_*′, p*_**s**_*′*_**s**_*′*, which in turn can be written solely in terms of *p*_**s**_ using Eqs. 41a–41d. These quantities depend on the pair of specific strategies **s, s**′ ∈ { **s**_1_, **s**_2_, **s**_3_, **s**_4_ } and on the underlying game transition pattern described in Eq. 1.

#### A.5 Diffusion approximation

Assuming Eq. 40 always holds for a pair of strategies **s** and **s**′, we study a one-dimensional diffusion process with respect to a random variable *p*_**s**_. Within the short time interval *t* → *t* + Δ*t* we have,

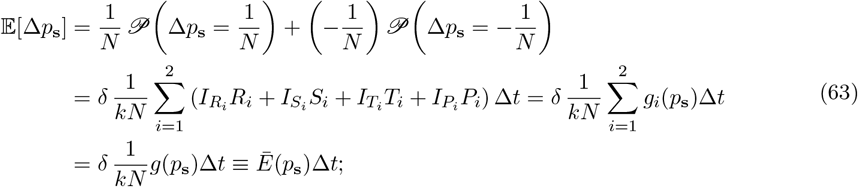

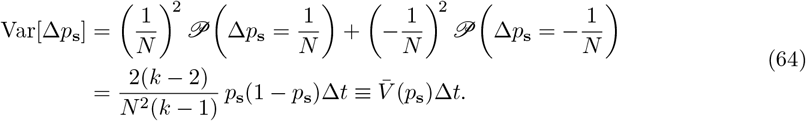

The fixation probability *ϕ*_**ss**_*′* (*y*) of strategy **s** in a resident population of **s**′ − players, given an initial frequency *p*_**s**_(*t* = 0) = *y*, satisfies the following differential equation,

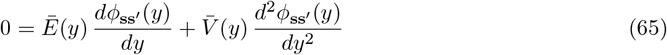

The solution of Eq. 65 is

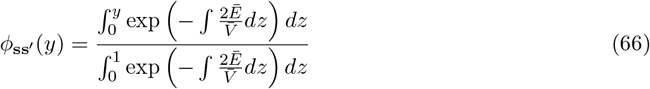

where, for *δ* ≪ 1,

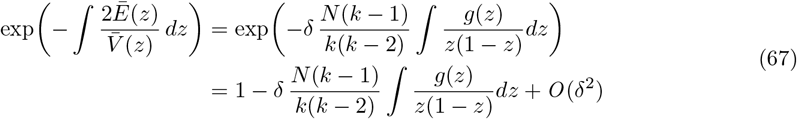

The fixation probability of *y* fraction of **s**-players in a population of **s**′-players is,

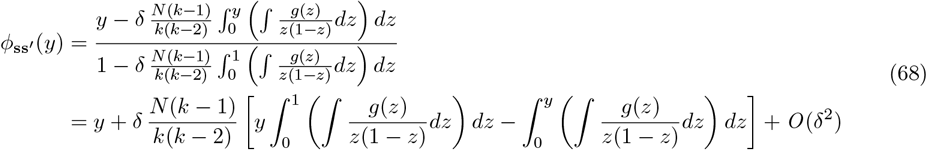

The fixation probability of *y* fraction of **s**′-player in a population of **s**-players is

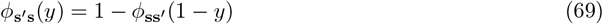

The difference in fixation probabilities is

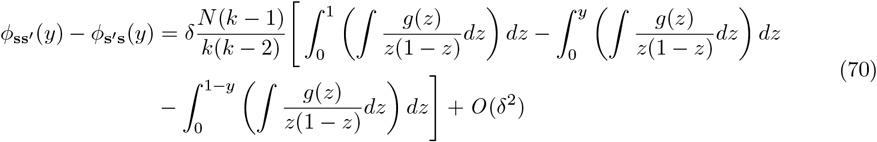

For sufficiently small *y*, we have,

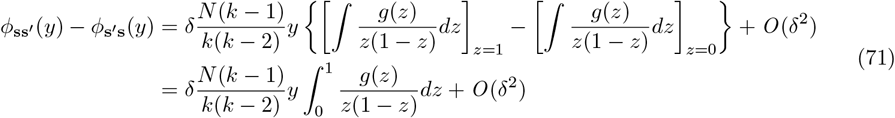

For a sufficiently large population and *y* = 1*/N*, we have

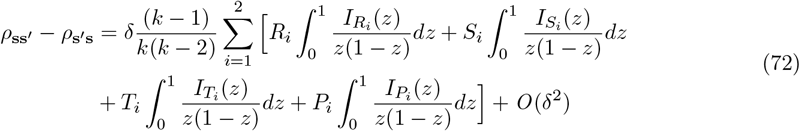

To calculate the conditions in Eq. 5, first we need to write 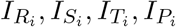 for all *i* ∈ {1, 2} in terms of *p*_**s**_ for pairs of strategies **s, s**′ ∈ {**s**_1_, **s**_2_, **s**_3_, **s**_4_ }. Note that Eq. 72 holds for competition between any pair of strategies and across all update rules, not only the death-birth update. Below, we systematically study the competition between all possible pairs of state-dependent strategies.

#### A.6 Competition between *CC* and *CD* strategy

Recall that our generalized state-dependent strategies **s** and **s**′ are defined by their action pairs: **s** = (*a*_1_, *a*_2_), 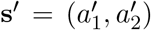. In the specific pairwise competition between the strategies *CC* and *CD*, we have, **s** = **s**_1_ = (*C, C*) and **s**′ = **s**_2_ = (*C, D*). Thus, for this case, *a*_1_ = *C, a*_2_ = *C*, 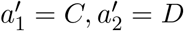 . The dynamics of 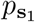due to the interaction between **s**_1_ and **s**_2_ then follows the same format as Eq. 19 with,

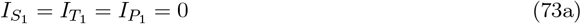

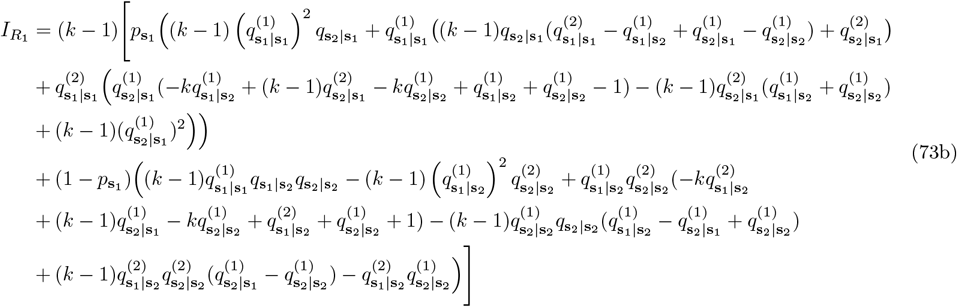

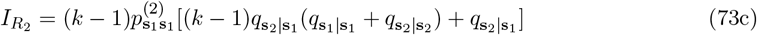

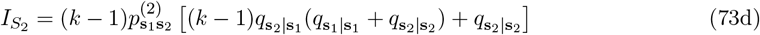

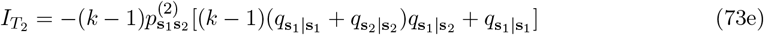

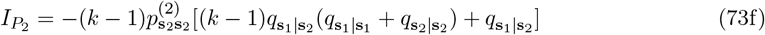

Note that Eqs. 73a-73f are in terms of *q* _*X*|*Y*_, 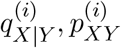 for all *X, Y* ∈ {**s**_1_, **s**_2_}. Using Eqs. 41a-41d we can write *q*_*X*|*Y*_ in terms of 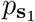 only. Next, we calculate 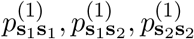 using Eq. 62 with the corresponding matrices,

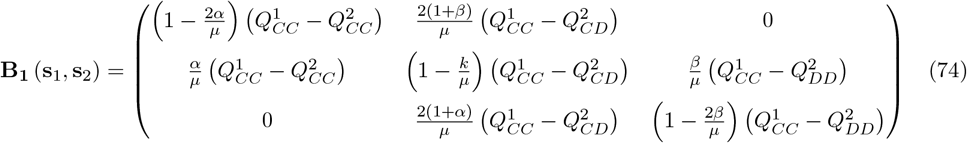

and

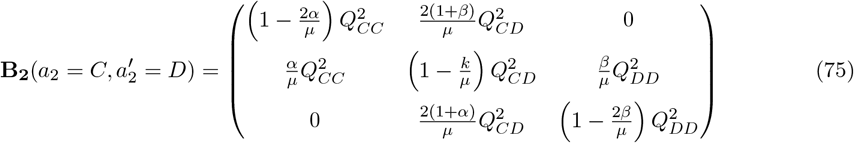

Using matrices **B**_**1**_ (**s**_1_, **s**_2_) and **B**_**2**_(*a*_2_ = *C*, 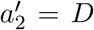), solving Eq. 62, substituting the solutions into Eqs. 73a-73f, we get 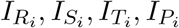 in terms of 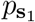 only. Then, inserting 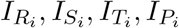 in Eq. 72, we obtain the difference in fixation probabilities of one strategy in a population of the other, 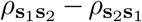 .

#### A.7 Competition between *CC* and *DC* strategy

Analogously, in the specific competition between the strategies *CC* and *DC*, we have, **s** = **s**_1_ = (*C, C*) and **s**′ = **s**_3_ = (*D, C*). Thus, for this case, *a*_1_ = *C, a*_2_ = *C*, 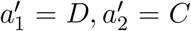. The dynamics of 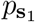due to the interaction between **s**_1_ and **s**_3_ then follows the same format as Eq. 19 with,

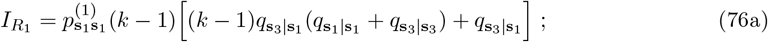

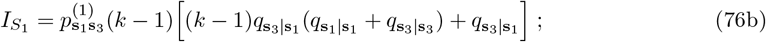

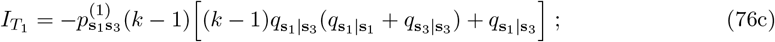

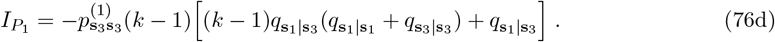

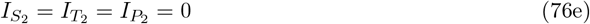

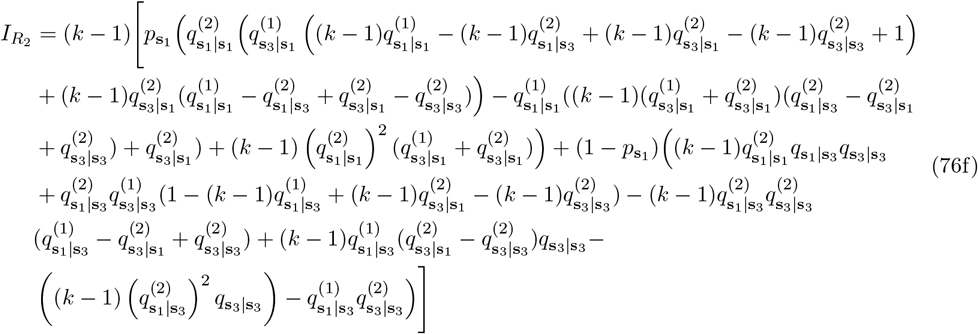

Next, we calculate 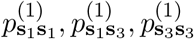 using Eq. 62 with the corresponding matrices,

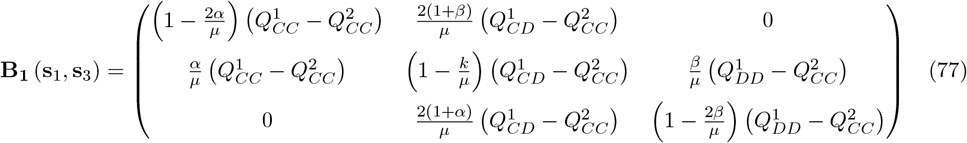

and

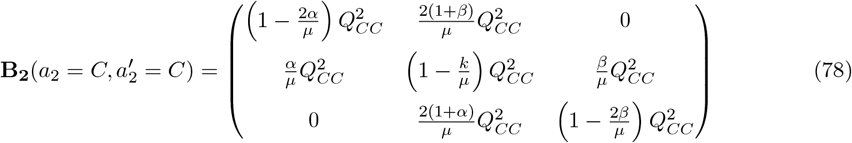

Using matrices **B**_**1**_ (**s**_1_, **s**_3_) and **B**_**2**_(*a*_2_ = *C*, 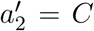), solving Eq. 62, substituting the solutions into Eqs. 76a-76f, we get 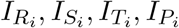 in terms of 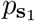only. Then, inserting 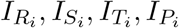 in Eq. 72, we obtain the difference in fixation probabilities of one strategy in a population of the other, 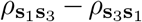 .

#### A.8 Competition between *CC* and *DD* strategy

Analogously, in the specific competition between the strategies *CC* and *DD*, we have, **s** = **s**_1_ = (*C, C*) and **s**′ = **s**_4_ = (*D, D*). Thus, for this case, *a*_1_ = *C, a*_2_ = *C*, 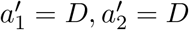. The dynamics of 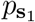 under the interaction between **s**_1_ and **s**_4_ then follows the same format as Eq. 19 with,

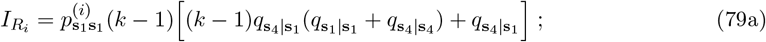

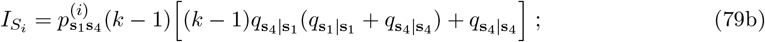

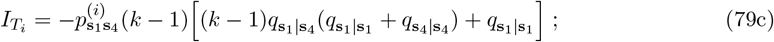

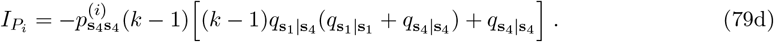

Next, we calculate 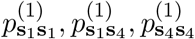 using Eq. 62 with the corresponding matrices,

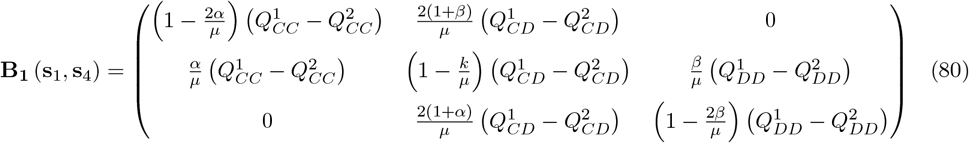

and

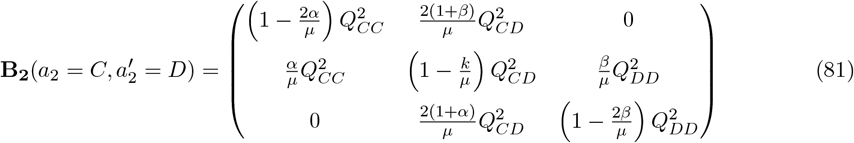

Using matrices **B**_**1**_ (**s**_1_, **s**_4_) and **B**_**2**_(*a*_2_ = *C*, 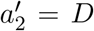), solving Eq. 62, substituting the solutions into Eqs. 79a-79d, we get 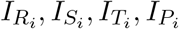 in terms of 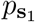 only. Then, inserting 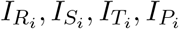 in Eq. 72, we obtain the difference in fixation probabilities of one strategy in a population of the other, 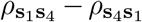 .

#### A.9 Competition between *CD* and *DC* strategy

Analogously, in the specific competition between the strategies *CD* and *DC*, we have, **s** = **s**_2_ = (*C, D*) and **s**′ = **s**_3_ = (*D, C*). Thus, for this case, *a*_1_ = *C, a*_2_ = *D*, 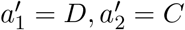. The dynamics of 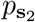 under the interaction between **s**_2_ and **s**_3_ then follows the same format as Eq. 19 with,

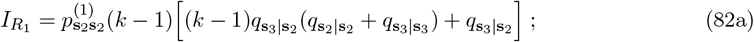

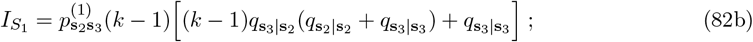

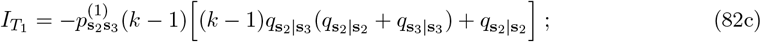

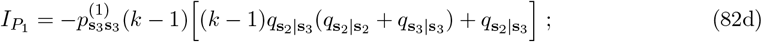

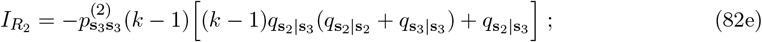

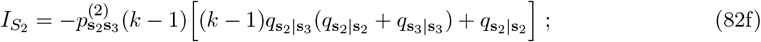

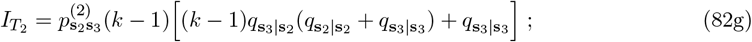

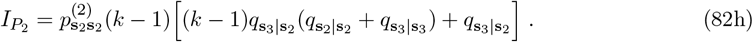

Next, we calculate 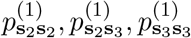 using Eq. 62 with the corresponding matrices,

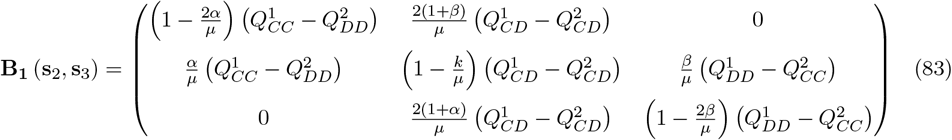

and

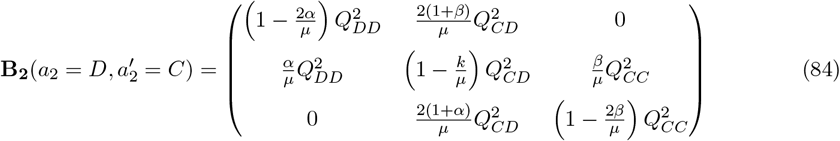

Using matrices **B**_**1**_ (**s**_2_, **s**_3_) and **B**_**2**_(*a*_2_ = *D*, 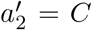), solving Eq. 62, substituting the solutions into Eqs. 82a-82h, we get 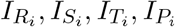 in terms of 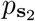 only. Then, inserting 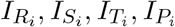 in Eq. 72, we obtain the difference in fixation probabilities of one strategy in a population of the other, 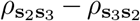 .

#### A.10 Competition between *CD* and *DD* strategy

Analogously, in the specific competition between the strategies *CD* and *DD*, we have, **s** = **s**_2_ = (*C, D*) and **s**′ = **s**_4_ = (*D, D*). Thus, for this case, *a*_1_ = *C, a*_2_ = *D*, 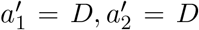. The dynamics of 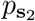 under the interaction between **s**_2_ and **s**_4_ then follows the same format as Eq. 19 with,

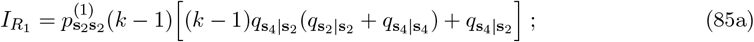

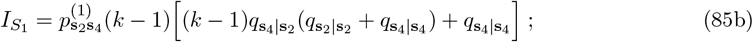

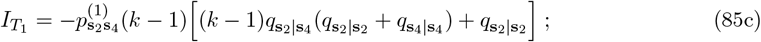

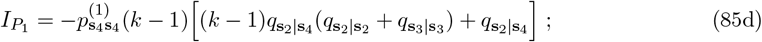

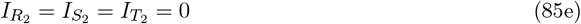

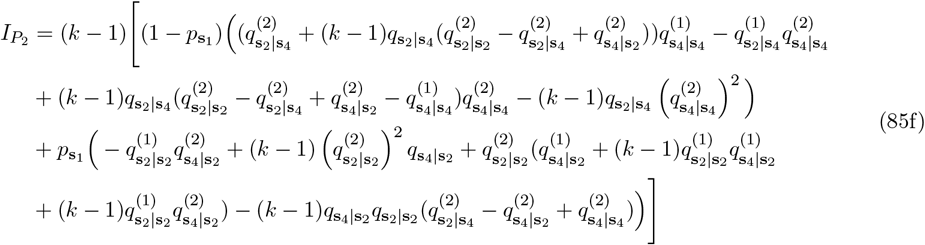

Next, we calculate 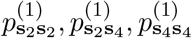 using Eq. 62 with the corresponding matrices,

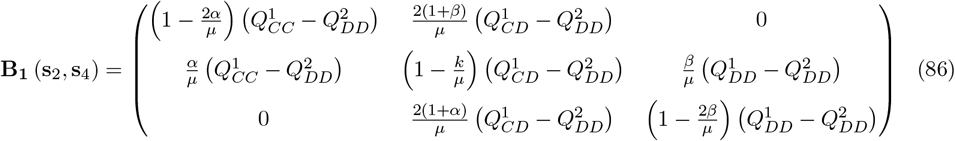

and

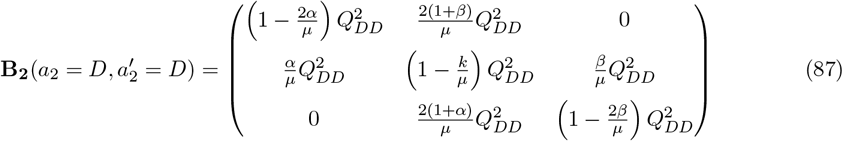

Using matrices **B**_**1**_ (**s**_2_, **s**_4_) and **B**_**2**_(*a*_2_ = *D*, 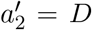), solving Eq. 62, substituting the solutions into Eqs. 85a-85f, we get 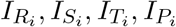 in terms of 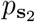 only. Then, inserting 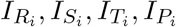 in Eq. 72, we obtain the difference in fixation probabilities of one strategy in a population of the other, 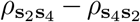 .

#### A.11 Competition between *DC* and *DD* strategy

Analogously, in the specific competition between the strategies *DC* and *DD*, we have, **s** = **s**_3_ = (*D, C*) and **s**′ = **s**_4_ = (*D, D*). Thus, for this case, *a*_1_ = *D, a*_2_ = *C*, 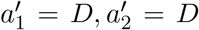. The dynamics of *p*_**s**_3 under the interaction between **s**_3_ and **s**_4_ then follows the same format as Eq. 19 with,

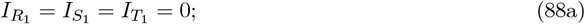

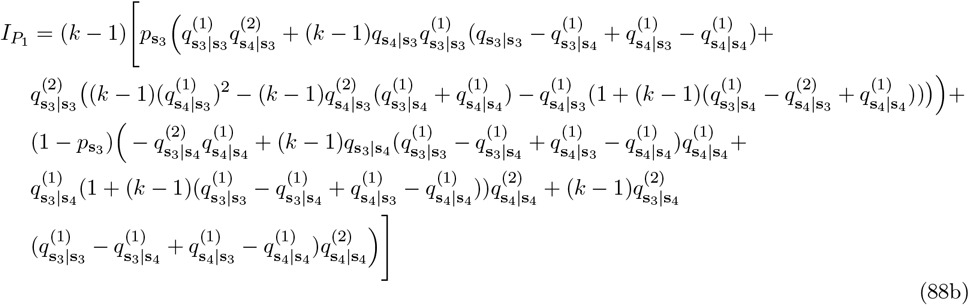

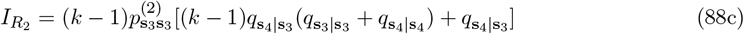

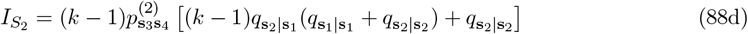

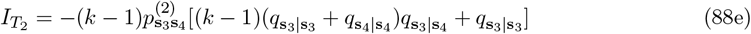

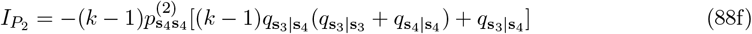

Next, we calculate 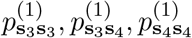 using Eq. 62 with the corresponding matrices,

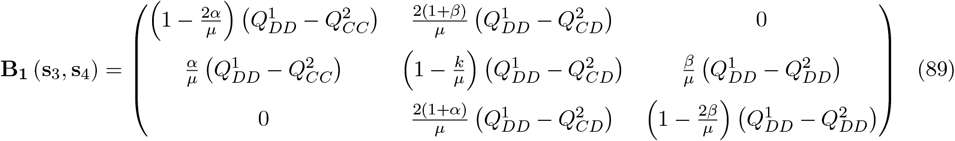

and

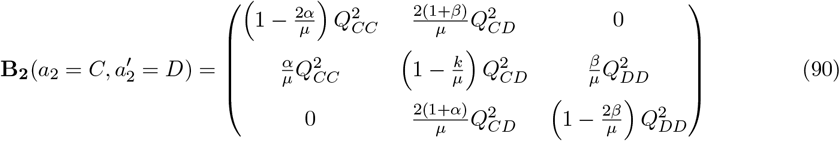

Using matrices **B**_**1**_ (**s**_3_, **s**_4_) and **B**_**2**_(*a*_2_ = *C*, 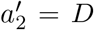), solving Eq. 62, substituting the solutions into Eqs. 88a-88f, we get 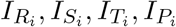 in terms of 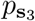only. Then, inserting 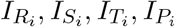 in Eq. 72, we obtain the difference in fixation probabilities of one strategy in a population of the other, 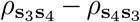 .

### B Analysis of 3 examples of game transition dynamics given in the main text

After having provided the general method to calculate *ρ*_**ss**_*′* −*ρ*_**s**_*′*_**s**_ for all **s, s**′ ∈ {**s**_1_, **s**_3_, **s**_3_, **s**_4_} under an arbitrary transition vector **Q**, in this section, we calculate the critical benefit-to-cost ratio required for the strategies *CC* and *CD* to be favored by selection for three representative game transition vectors discussed in the main text. Note that in the complete knowledge scenario, *DC* is also a game 2 cooperative strategy; however, our numerical simulations in the main text show that for these three representative examples, the *DC*−strategy is never favored by selection for any *b*_1_*/c* value. Therefore, we do not compute the selection condition for the *DC*−strategy.

In the main text, we consider donation game (which is a form of the Prisoners’ Dilemma game) in both state 1 and state 2, where,

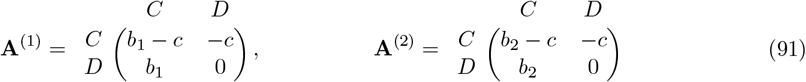

Since *P*_*i*_ = 0, we do not calculate *I*_*Pi*_ separately in the examples below.

#### B.1 Q_1_ = (1, 0, 0; 1, 0, 0)

The first example we consider is the deterministic transition with transition structure **Q**_**1**_ = (1, 0, 0; 1, 0, 0). Su et al. [9] have shown that, under this transition vector, game transitions can promote cooperation in the absence of state knowledge. Below, we will show how the knowledge of states affects the conditions for the emergence of different cooperative state-dependent strategies for the transition vector **Q**_**1**_.

##### B.1.1 Calculation of 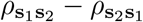

Following the methodology described in section A.6, we first calculate 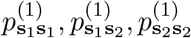 using the matrices **B**_**1**_ and **B**_2_ (using Eqs. 74 and 75) for the transition vector **Q**_**1**_ as follows,

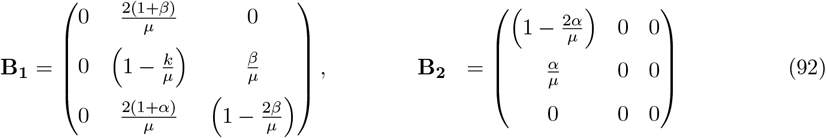

Inserting **B**_**1**_, **B**_**2**_ in Eq. 62 and using the large population limit *N* → ∞ we get,

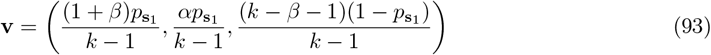

Using Eqs. 41a-41d, one can show that 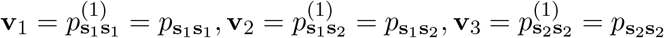. Thus under this game transition vector **Q**_**1**_, when *CC* and *CD* players interact, all edges always remain in game 1. Using Eq. 93 and the identity 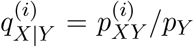 we write 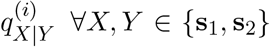 in terms of 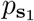 only. Substituting 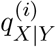 and Eq. 93 in Eqs. 73a-73f, we find that

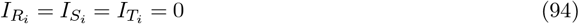

Then, inserting 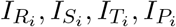 in Eq. 72, we obtain

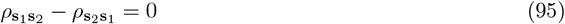

Under the game transition vector **Q**_1_, *CC* and *CD* players are neutral to each other.

##### B.1.2 Calculation of 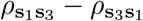

For interaction between **s**_1_ and **s**_3_ players, using Eqs. 77 and 78, the matrices **B**_**1**_ and **B**_2_ for the transition vector **Q**_**1**_ are,

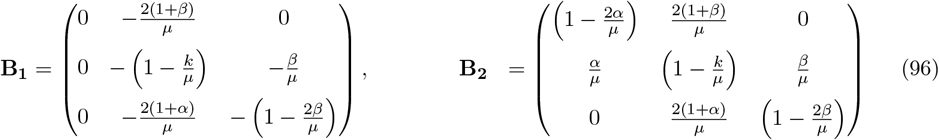

Inserting **B**_**1**_, **B**_**2**_ in Eq. 62 and using the large population limit *N* → ∞ we get,

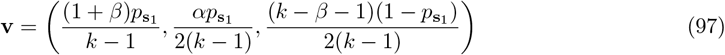

Analogously, substituting Eq. 97 in Eqs. 76a-76e, we find that

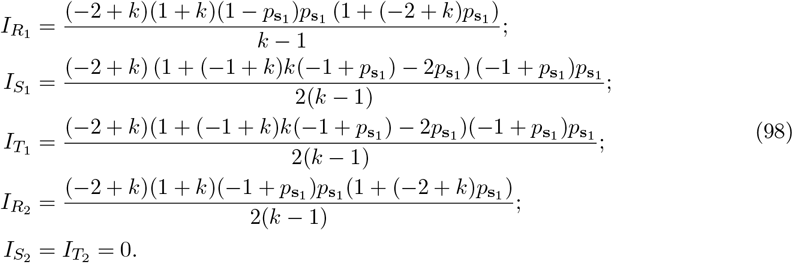

Then, inserting 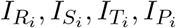 and donation game payoff matrix (Eq. 91) in Eq. 72, we obtain

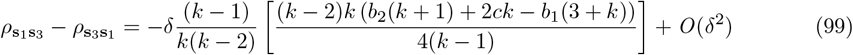

##### B.1.3 Calculation of 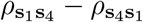

For interaction between **s**_1_ and **s**_4_ players, using Eqs. 80 and 81, the matrices **B**_**1**_ and **B**_2_ for the transition vector **Q**_**1**_ are,

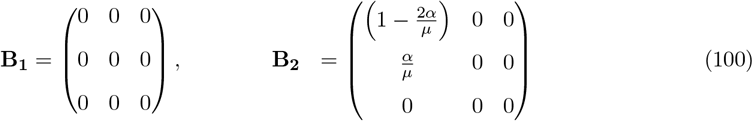

Inserting **B**_**1**_, **B**_**2**_ in Eq. 62 and using the large population limit *N* → ∞ we get,

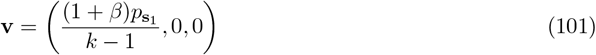

Analogously, substituting Eq. 101 in Eqs. 79a-79d, we find that

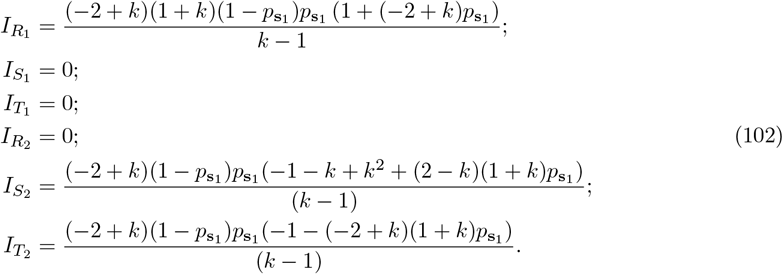

Then, inserting 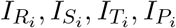 and donation game payoff matrix in Eq. 72, we obtain

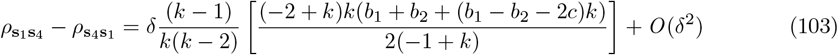

Using Eqs. 95,99, and 103, we get the condition for the *CC* strategy to be favored by selection under the game transition rule **Q**_**1**_,

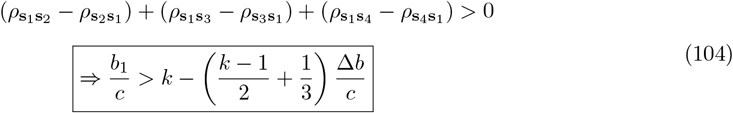

where, Δ*b* = *b*_1_ − *b*_2_. Eq. 104 corresponds to Eq. 6 in the main text with 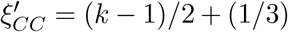 for the transition vector **Q**_**1**_.

##### B.1.4 Calculation of 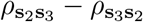

For interaction between **s**_2_ and **s**_3_ players, using Eqs. 83 and 84, the matrices **B**_**1**_ and **B**_2_ for the transition vector **Q**_**1**_ are,

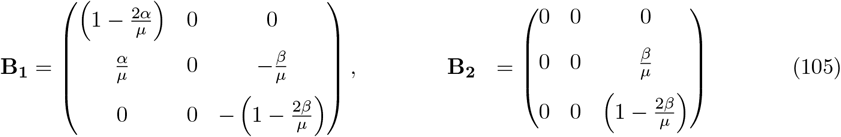

Inserting **B**_**1**_, **B**_**2**_ in Eq. 62 and using the large population limit *N* → ∞ we get,

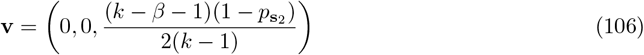

Analogously, substituting Eq. 106 in Eqs. 82a-82h, we find that

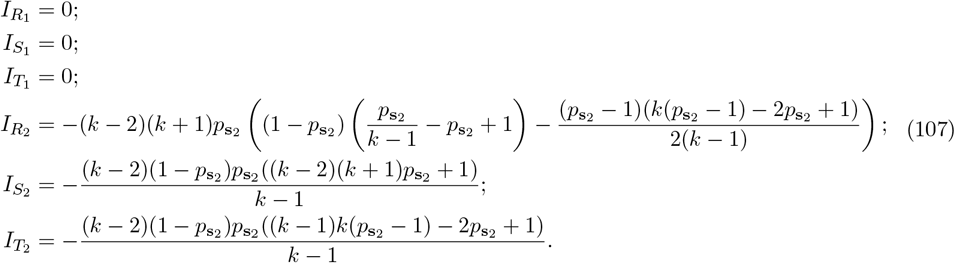

Then, inserting 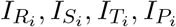 and donation game payoff matrix in Eq. 72, we obtain

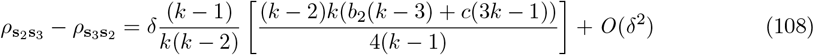

##### B.1.5 Calculation of 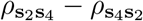

For interaction between **s**_2_ and **s**_4_ players, using Eqs. 86 and 87, the matrices **B**_**1**_ and **B**_2_ for the transition vector **Q**_**1**_ are,

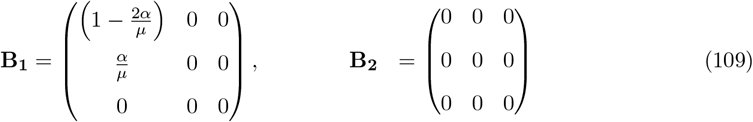

Inserting **B**_**1**_, **B**_**2**_ in Eq. 62 and using the large population limit *N* → ∞ we get,

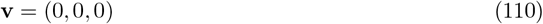

Thus, under this game transition vector **Q**_**1**_, when *CD* and *DD* players interact, all edges always remain in game 2. Analogously, substituting Eq. 110 in Eqs. 85a-85f, we find that

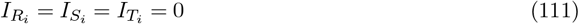

Then, inserting 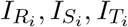 and donation game payoff matrix in Eq. 72, we obtain

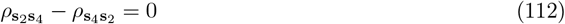

Under the game transition vector **Q**_1_, *CD* and *DD* players are neutral to each other.

Using Eqs. 95,108, and 112, we get the condition for the *CD* strategy to be favored by selection under the game transition rule **Q**_**1**_,

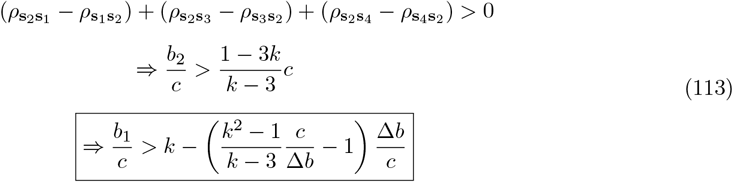

where, Δ*b* = *b* − *b* . Eq. 113 corresponds to Eq. 6 in the main text with 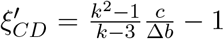 for the transition vector **Q**_**1**_.

#### B.2 Q_**2**_ = (1, 0, 0; 1, 1, 1)

Here we will show how the knowledge of game states affects the conditions for the emergence of different cooperative state-dependent strategies for the transition vector **Q**_**2**_.

##### B.2.1 Calculation of 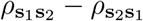

For interaction between **s**_1_ and **s**_2_ players, using Eqs. 74 and 75 the matrices **B**_**1**_ and **B**_2_ for the transition vector **Q**_**2**_ are,

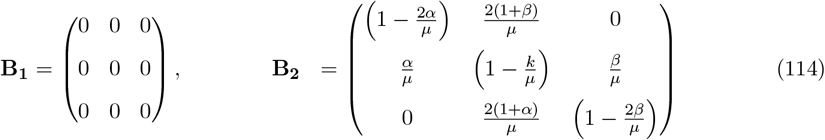

Inserting **B**_**1**_, **B**_**2**_ in Eq. 62 and using the large population limit *N* → ∞ we get,

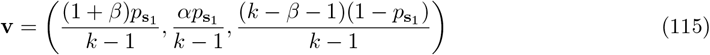

Analogously, substituting Eq. 115 in Eqs. 73a-73f, we find that

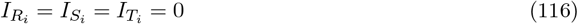

Then, inserting 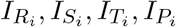 in Eq. 72, we obtain

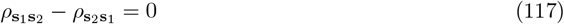

Under the game transition vector **Q**_2_, *CC* and *CD* players are neutral to each other.

##### B.2.2 Calculation of 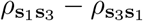

For interaction between **s**_1_ and **s**_3_ players, using Eqs. 77 and 78, the matrices **B**_**1**_ and **B**_2_ for the transition vector **Q**_**2**_ are,

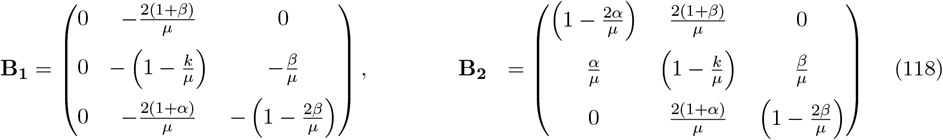

Inserting **B**_**1**_, **B**_**2**_ in Eq. 62 and using the large population limit *N* → ∞ we get,

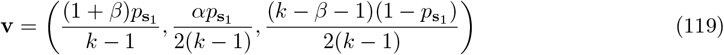

Analogously, substituting Eq. 119 in Eqs. 76a-76e, we find that

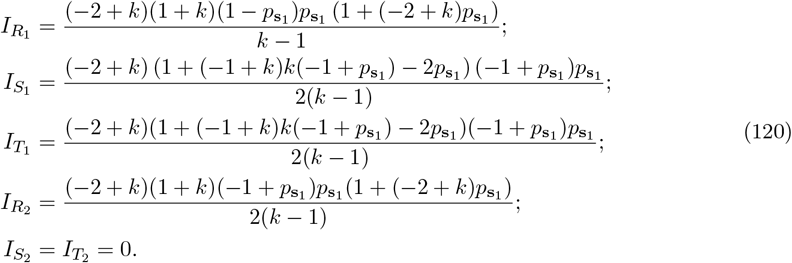

Then, inserting 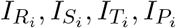and using donation game payoff matrix in Eq. 72, we obtain

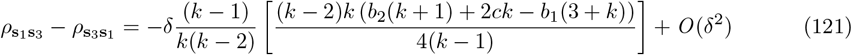

##### B.2.3 Calculation of 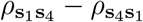

For interaction between **s**_1_ and **s**_4_ players, using Eqs. 80 and 93, the matrices **B**_**1**_ and **B**_2_ for the transition vector **Q**_**2**_, are,

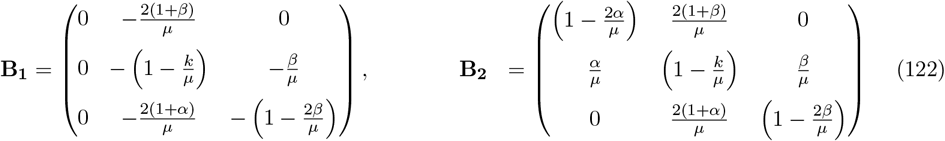

Inserting **B**_**1**_, **B**_**2**_ in Eq. 62 and using the large population limit *N* → ∞ we get,

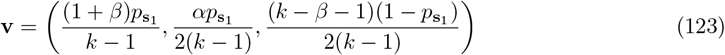

Analogously, substituting Eq. 123 in Eqs. 79a-79d, we find that

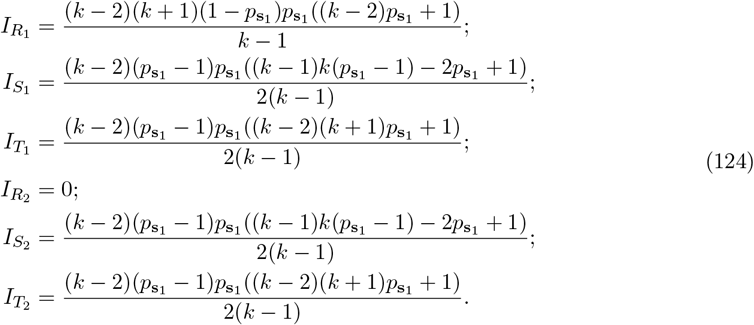

Then, inserting 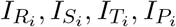 and donation game payoff matrix in Eq. 72, we obtain

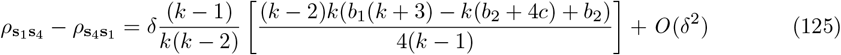

For 0 *< y <* 1, using Eqs. 117,121, and 125, we get the condition for the *CC* strategy to be favored by selection under the game transition rule **Q**_**2**_,

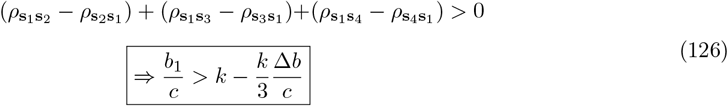

where, Δ*b* = *b*_1_ − *b*_2_. Eq. 126 corresponds to Eq. 6 in the main text with 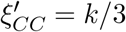 for the transition vector **Q**_**2**_.

##### B.2.4 Calculation of 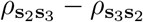

For interaction between **s**_2_ and **s**_3_ players, using Eqs. 83 and 84, the matrices **B**_**1**_ and **B**_2_ for the transition vector **Q**_**2**_ are,

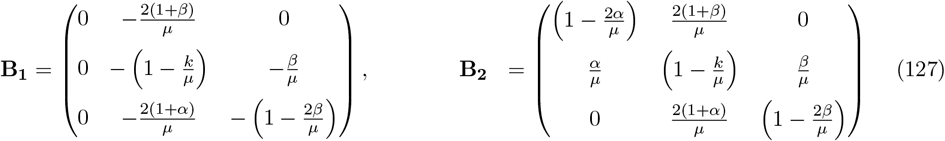

Inserting **B**_**1**_, **B**_**2**_ in Eq. 62 and using the large population limit *N* → ∞ we get,

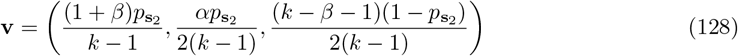

Analogously, substituting Eq. 128 in Eqs. 82a-82h, we find that

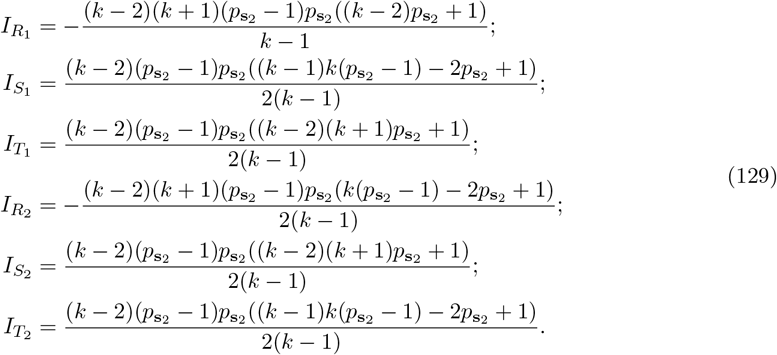

Then, inserting 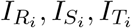 and donation game payoff matrix in Eq. 72, we obtain

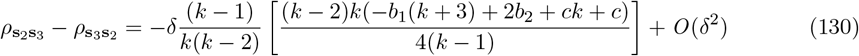

##### B.2.5 Calculation of 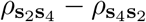

For interaction between **s**_2_ and **s**_4_ players, using Eqs. 86 and 87, the matrices **B**_**1**_ and **B**_2_ for the transition vector **Q**_**2**_ are,

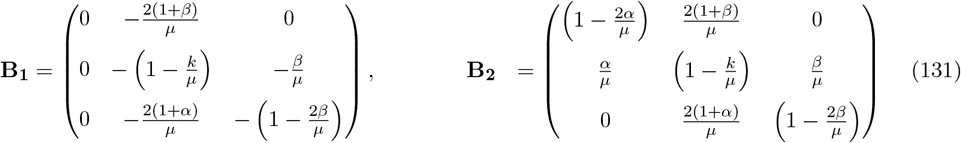

Inserting **B**_**1**_, **B**_**2**_ in Eq. 62 and using the large population limit *N* → ∞ we get,

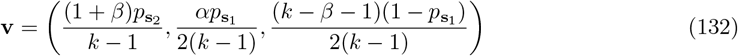

Analogously, substituting Eq. 132 in Eqs. 85a-85f, we find that

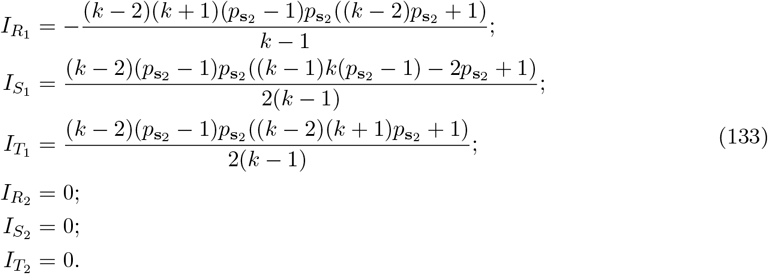

Then, inserting 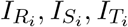 and donation game payoff matrix in Eq. 72, we obtain

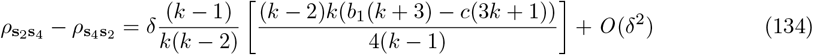

Using Eqs. 117,130, and 134, we get the condition for *CD* strategy to be favored by selection under game transition rule **Q**_2_,

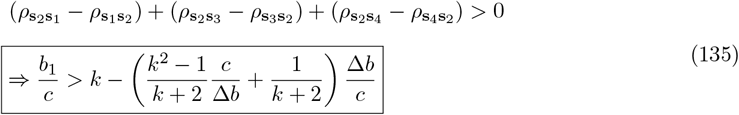

where, Δ*b* = *b* − *b* . Eq. 135 corresponds to Eq. 6 in the main text with 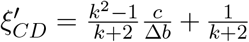 for the transition vector **Q**_**2**_.

##### B.3 Q_3_ = (1, 1, 0; 1, 0, 0)

Here we will show how the knowledge of game states affects the conditions for the emergence of different cooperative state-dependent strategies for the transition vector **Q**_**3**_.

###### B.3.1 Calculation of 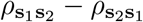

For interaction between **s**_1_ and **s**_2_ players, using Eqs. 74 and 75 the matrices **B**_**1**_ and **B**_2_ for the transition vector **Q**_**3**_ are,

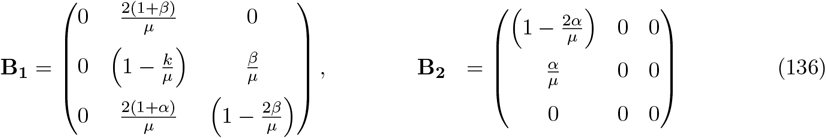

Inserting **B**_**1**_, **B**_**2**_ in Eq. 62 and using the large population limit *N* → ∞ we get,

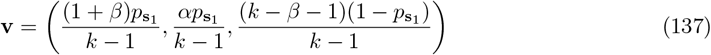

Analogously, substituting Eq. 137 in Eqs. 73a-73f, we find that

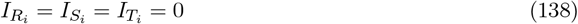

Then, inserting 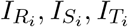 in Eq. 72, we obtain

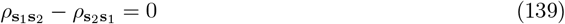

Under the game transition vector **Q**_3_, *CC* and *CD* players are neutral to each other.

###### B.3.2 Calculation of 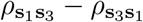

For interaction between **s**_1_ and **s**_3_ players, using Eqs. 77 and 78, the matrices **B**_**1**_ and **B**_2_ for the transition vector **Q**_**3**_ are,

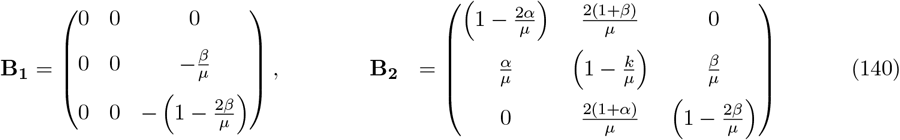

Inserting **B**_**1**_, **B**_**2**_ in Eq. 62 and using the large population limit *N* → ∞ we get,

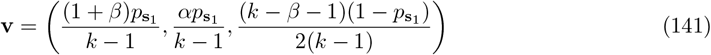

Analogously, substituting Eq. 141 in Eqs. 76a-76e, we find that

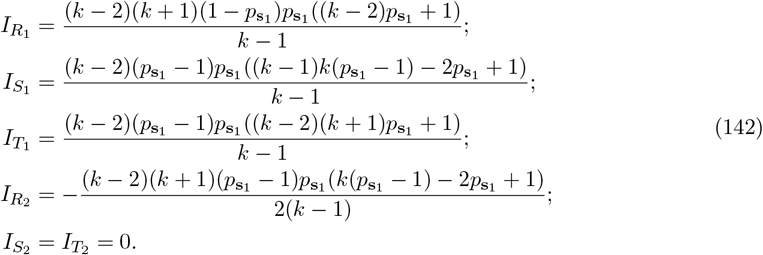

Then, inserting 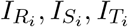 and donation game payoff matrix in Eq. 72, we obtain

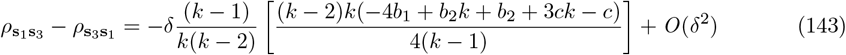

###### B.3.3 Calculation of 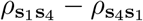

For interaction between **s**_1_ and **s**_4_ players, using Eqs. 80 and 81, the matrices **B**_**1**_ and **B**_2_ for the transition vector **Q**_**3**_ are,

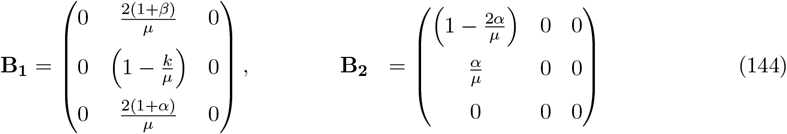

Inserting **B**_**1**_, **B**_**2**_ in Eq. 62 and using the large population limit *N* → ∞ we get,

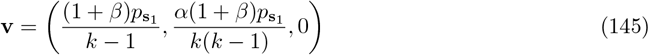

Analogously, substituting Eq. 145 in Eqs. 79a-79d, we find that

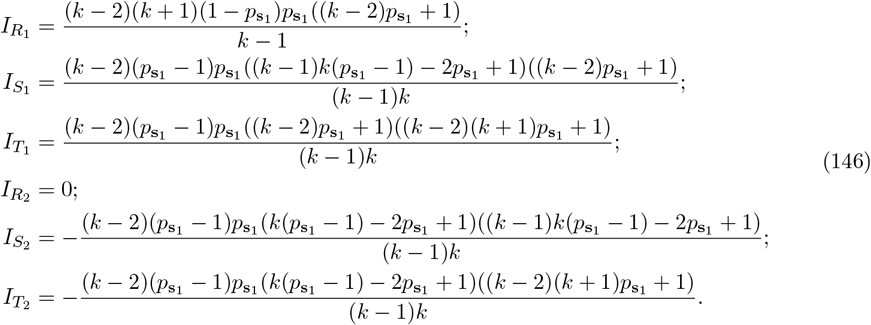

Then, inserting 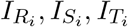 and donation game payoff matrix in Eq. 72, we obtain

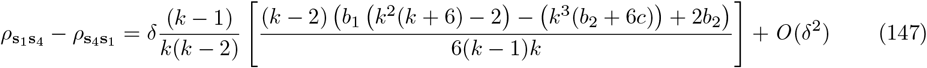

Using Eqs. 139,143, and 147, we get the condition for the *CC* strategy to be favored by selection under the game transition rule **Q**_**3**_,

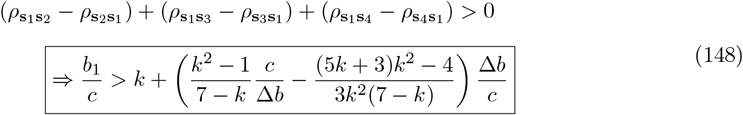

where, Δ*b* = *b* −*b* . Eq. 148 corresponds to Eq. 6 in the main text with 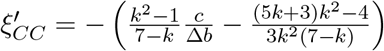 for the transition vector **Q**_**3**_.

###### B.3.4 Calculation of 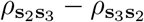

For interaction between **s**_2_ and **s**_3_ players, using Eqs. 83 and 84, the matrices **B**_**1**_ and **B**_2_ for the transition vector **Q**_**3**_ are,

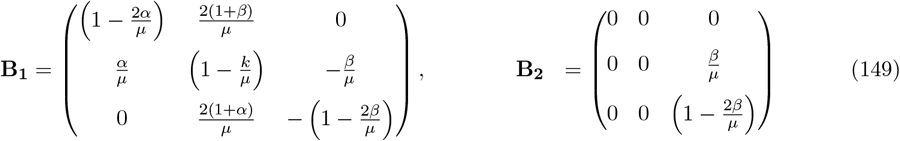

Inserting **B**_**1**_, **B**_**2**_ in Eq. 62 and using the large population limit *N* → ∞ we get,

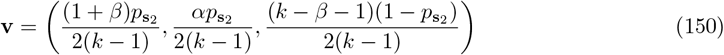

Analogously, substituting Eq. 150 in Eqs. 82a-82h, we find that

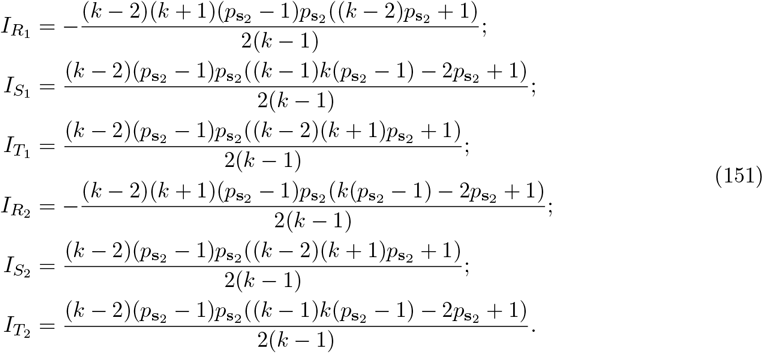

Then, inserting 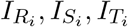 and donation game payoff matrix in Eq. 72, we obtain

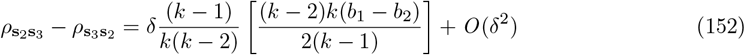

###### B.3.5 Calculation of 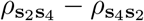

For interaction between **s**_2_ and **s**_4_ players, using Eqs. 86 and 87, the matrices **B**_**1**_ and **B**_2_ for the transition vector **Q**_**3**_ are,

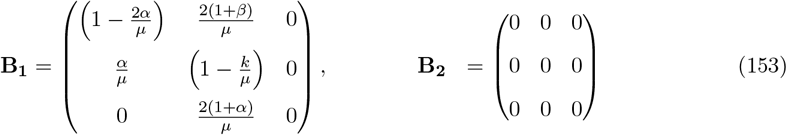

Inserting **B**_**1**_, **B**_**2**_ in Eq. 62 and using the large population limit *N* → ∞ we get,

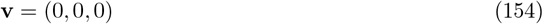

Thus, under this game transition vector **Q**_**3**_, when *CD* and *DD* players interact, all edges always remain in game 2. Analogously, substituting Eq. 154 in Eqs. 85a-85f, we find that

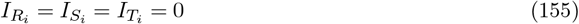

Then, inserting 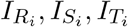 and donation game payoff matrix in Eq. 72, we obtain

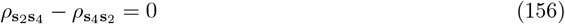

Under the game transition vector **Q**_3_, *CD* and *DD* players are neutral to each other.

For 0 *< y <* 1, using Eqs. 139,152, and 156, we get the condition for the *CD* strategy to be favored by selection under the game transition rule **Q**_**3**_,

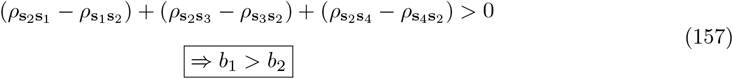

#### 2 Game transitions without knowledge of the states

In this section, we assume that players have no information about the state (game) they are currently in. Consequently, they adopt state-independent strategies, using a uniform action across all states. As in section 1, players interact with their neighbors only once, and their strategy does not take past actions into account. For transitions between two game states, a state-independent strategy is represented by a single action *a*, written as **s**_**I**_ = (*a*), where *a* ∈ {*C, D*} . A player using strategy **s**_*I*_ therefore uses the same action in both game 1 and game 2. Since a state-independent strategy corresponds to the special case of a state-dependent strategy with *a*_1_ = *a*_2_ = *a*, it follows that the set of state-dependent strategies (Eq. 2) contains the state-independent strategies as a strict subset. Because each player may choose either *C* or *D*, the set of pure state-independent strategies consists of 2 strategies only, 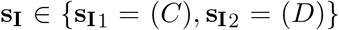 . As **s**_**I**_ ⊂ **s**_**d**_, we note that, 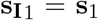 and 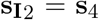 (see Eq. 2).

Similar to section 1, here we want to analytically derive the condition for cooperators (strategy **s**_1_) to be favored by selection over defection (strategy **s**_4_) for different transition rules given by Eq. 1. Thus, for game transitions without knowledge of the game states, selection favors cooperators relative to defectors if [2],

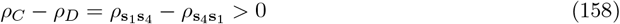

The competition between *C*(= **s**_1_) and *D*(= **s**_4_) strategies has already been studied in section A.8 for an arbitrary transition vector **Q**. Following the methodologies developed in section A.8 and using the donation game payoff matrices (Eq. 91), one can show [9] for any arbitrary transition vector **Q**,

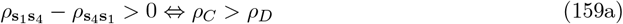

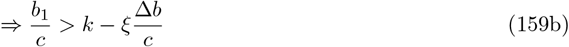

where

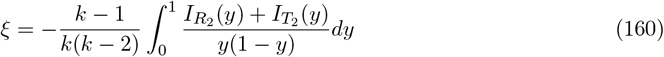

for death-birth updating. Since the value of 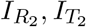 depends on the specific transition rule, the *ξ* value captures the underlying game transition pattern. Below, we calculate the value of *ξ* for a few examples of transition vectors considered in the main text.

##### 2.1 Q_**1**_ = (1, 0, 0; 1, 0, 0)

Eq. 103 shows the difference in fixation probabilities when there are only 2 competing strategies **s**_1_ and **s**_4_ in the population for the transition vector **Q**_**1**_. Substituting Eq. 103 in Eq. 158, we get the condition for cooperators to be favored by selection in the absence of state knowledge, for the transition vector **Q**_**1**_,

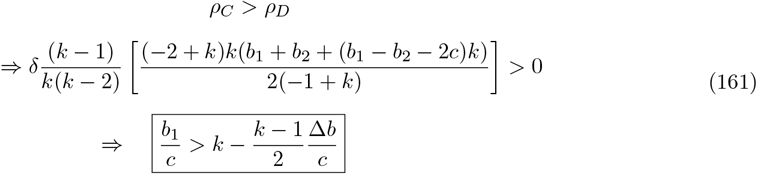

Eq. 161 corresponds to Eq. 5 in the main text with *ξ* = (*k* − 1)*/*2 for the transition vector **Q**_**1**_.

##### 2.2 Q_**2**_ = (1, 0, 0; 1, 1, 1)

Eq. 125 shows the difference in fixation probabilities when there are only 2 competing strategies **s**_1_ and **s**_4_ in the population for the transition vector **Q**_**2**_. Substituting Eq. 125 in Eq. 158, we get the condition for cooperators to be favored by selection in the absence of state knowledge, for the transition vector **Q**_**2**_,

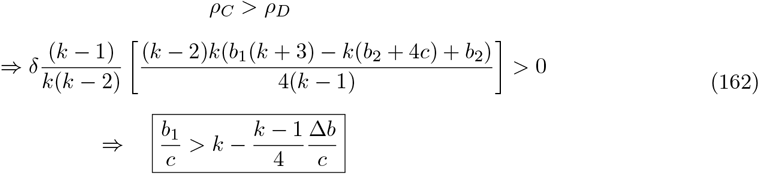

Eq. 162 corresponds to Eq. 5 in the main text with *ξ* = (*k* − 1)*/*4 for the transition vector **Q**_**2**_.

##### 2.3 Q_**3**_ = (1, 1, 0; 1, 0, 0)

Eq. 147 shows the difference in fixation probabilities when there are only 2 competing strategies **s**_1_ and **s**_4_ in the population for the transition vector **Q**_**3**_. Substituting Eq. 147 in Eq. 158, we get the condition for cooperators to be favored by selection in the absence of state knowledge, for the transition vector **Q**_**3**_,

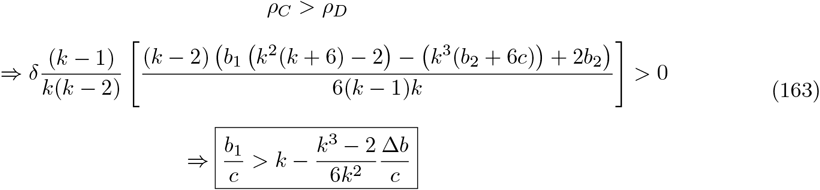

Eq. 163 corresponds to Eq. 5 in the main text with *ξ* = (*k*^3^ − 2)*/*6*k*^2^ for the transition vector **Q**_**3**_.

In the following, Table 1 shows the value of *ξ* for all 64 deterministic transition vectors. To calculate *ξ* values for each transition vector **Q** in table 1, we follow the same method described in section A.8 except for **Q**_**a**_ = (1, 1, 1; 0, 0, 0). This transition vector doesn’t allow any transition between states. For our initial condition of all edges in state 1, the population remains permanently in game 1. Thus, the critical (*b*_1_*/c*)^∗^ value remains the same as the threshold obtained for a single game, which is given by the degree *k* of the network.

#### 3 An alternate method to study game transitions

In this section, we introduce an alternative approach to studying evolutionary dynamics with game transitions. In our model, as the game played between the pair of players changes with time in a manner that depends on their actions, the individual strategies can also be updated following a death-birth process. However, the game being played across an edge evolves much faster than individuals’ strategies, because at each time step only one of *N* individuals updates her strategy, whereas the game updates across all *kN/*2 edges. Therefore, for a sufficiently large population size *N*, we can assume two individuals connected by an edge act “as if” they are effectively playing an infinitely repeated game (over several time steps) before their strategies are updated. Under this assumption, we can use the framework of repeated games to calculate each player’s effective payoff along an edge, under the effect of game transitions. To calculate the effective payoff using the Markov chain method, here, we represent a state-dependent strategy with its probability of cooperation in each state. Let 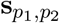 denote a state-dependent strategy with which, at each time step, a player cooperates with probability *p*_1_ in game 1 and with probability *p*_2_ in game 2. **s**_1,1_ thus corresponds to *CC*(= **s**_1_) strategy, **s**_1,0_ corresponds to *CD*(= **s**_2_) strategy, **s**_0,1_ corresponds to *DC*(= **s**_3_) strategy and **s**_0,0_ corresponds to *DD*(= **s**_4_) strategy (see Eq. 2). For a state-independent strategy, a player chooses cooperation with the same probability *p*_1_ = *p*_2_ = *p* in both states. Let **s**_*p*_ denote a state-independent strategy with which, at each time step, a player cooperates with probability *p* in all states. Thus, **s**_1_ corresponds to pure cooperators *C* and **s**_0_ corresponds to pure defectors *D*. Note that pure cooperators (**s**_1_) and defectors (**s**_0_) are present in the set of state-dependent strategies as well, which earlier we denoted as *CC*(= **s**_1_) and *DD*(= **s**_4_) respectively. In the following, we show how to calculate the payoff between two state-dependent strategies connected by an edge. Since any state-independent strategy can be deduced from a state-dependent strategy with *p*_1_ = *p*_2_ = *p* in both states, the same method also applies to state-independent strategies.

##### 3.1 Calculation of payoffs

Given player 1’s strategy 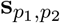 and player 2’s strategy 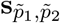, the payoff between the two strategies can be calculated using the Markov chain method [10]. The states of the Markov chain correspond to eight possible outcomes *ω* = (*i, a, ã*) of a given round. Here, *i* ∈ {1, 2} corresponds to the game state, and *a, ã* ∈ {*C, D*} are player 1’s and player 2’s actions, respectively, in state *i*. The transition probability to move from state *ω* = (*i, a, ã*) in one round to *w*′ = (*i*′, *a*′, *ã*′) in the next round is written as,

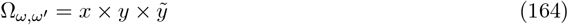

The first factor,

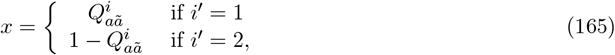

reflects the probability of moving from state *i* to state *i*′, given the players’ actions. The other two factors are,

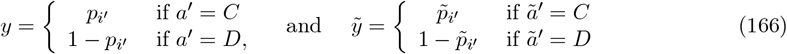

This leads to an 8 ×8 transition matrix 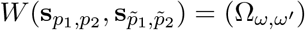. This transition matrix has a unique left eigen vector 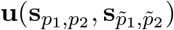, which can be found by solving the linear equation 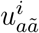. The entries of this eigenvector give the frequency with which players observe the outcome *ω* = (*i, a, ã*) over the course of the game. When both states are represented by donation games (Eq. 91), for a given transition vector **Q**, we can therefore compute the expected payoff of the players as

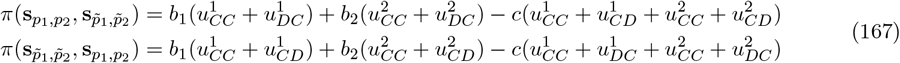

##### 3.2 Game transitions without knowledge of the states

When players do not possess the knowledge of the states, they adopt state-independent strategies, **s**_*p*_. In this scenario, there are only two pure strategies competing in the population, cooperators (*C*) and defectors (*D*). In the following, using the alternate method, we show when a pure cooperator (**s**_1_) strategy is favored by selection over a defector (**s**_0_) strategy.

###### Payoff between two cooperators

When two **s**_1_ players interact along an edge over several time-steps, using Eqs. 164-166 the unique eigen vector **u**(**s**_1_, **s**_1_) can be written as,

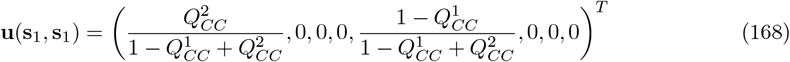

Substituting Eq. 168 in Eq. 167, we calculate the payoff of each cooperators 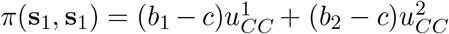, where 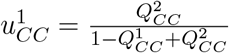, and 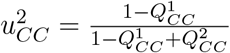.

###### Payoff between a cooperator and a defector

When an edge connects a **s**_1_ and a **s**_0_ player and they interact over several time-steps, using Eqs. 164-166 the unique eigen vector **u**(**s**_1_, **s**_0_) can be written as,

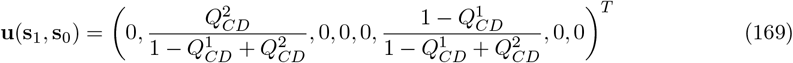

Analogously, using Eq. 169, the payoff of the cooperator and the defector is 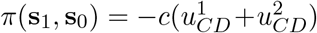, and 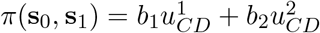, respectively, where 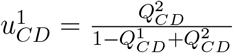, and 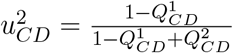.

###### Payoff between two defectors

When two **s**_0_ players interact along an edge over several time-steps, using Eqs. 164-166 the unique eigen vector **u**(**s**_0_, **s**_0_) can be written as,

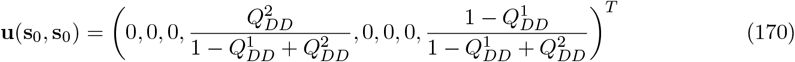

For donation games in both states, two defectors always receive a payoff of 0.

Thus, our model of game transitions on networks is equivalent to a situation where all players “as if” are playing a single game with an “effective” payoff matrix,

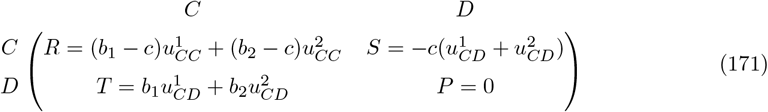

where, values of 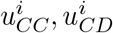 are according to Eqs. 168, 169. Note that Eq. 171 is true not only for death-birth updating but for other update rules as well, such as pairwise comparison, imitation update, etc.

Since evolutionary dynamics with game transitions without knowledge of the states can be effectively thought of as a game between cooperators and defectors with a single modified payoff matrix given by Eq. 171, we can use the well-established results for a single game to determine the conditions for the success of cooperators over defectors. Below, we use the “Sigma rule” [3] to find the condition for cooperators to be favored relative to defectors, *ρ*_*C*_ *> ρ*_*D*_ (Eq. 159a).

###### 3.2.1 Analysis based on Sigma rule

For interaction between 2 strategies, *A* and *B*, governed by a single payoff matrix,

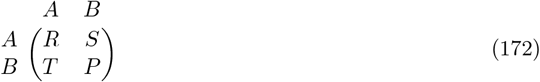

Tarnita et al. [3] have shown that in the limit of weak selection, the condition for strategy *A* to be favored over strategy *B* is equivalent to

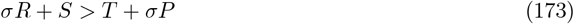

which is termed as a “sigma rule”. The parameter *σ* depends on the population structure and the update rule, but is independent of the payoff values. For a random regular graph and under the death-birth update rule, *σ* = (*k* + 1)*/*(*k*− 1).

Substituting the effective payoff matrix (Eq. 171) in Eq. 173 with the corresponding *R, S, T, P* values; for death-birth updating, the condition for cooperators (*C*) favored by selection over defectors (*D*), follows again the same form as Eq. 159b with

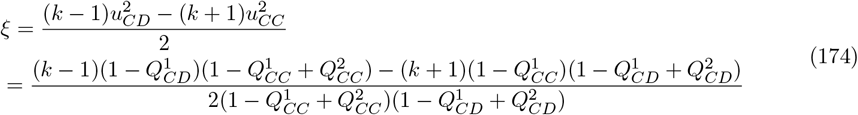

which was obtained by using 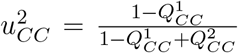 and 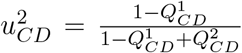 from Eqs. 168 and 169, respectively.

However, note that for the competition between pure strategies **s**_1_ and **s**_0_, for the following classes of deterministic game transitions characterized by the specific values of 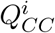 and 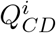,

1. 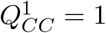, and 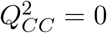;
2. 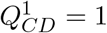, and 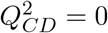;

the value of *ξ* given by Eq. 174 blows up. For these two classes of transition vectors, the corresponding Markov chain is not ergodic, and the long-time stationary distribution **u**(**s**_1_, **s**_1_) (**u**(**s**_1_, **s**_0_)) for class 1 (class 2) depends on the initial state. There is another class of game transitions characterized by 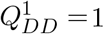, and 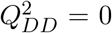 for which the Markov chain is also not ergodic; however, since two defectors always receive a payoff of 0, it doesn’t affect the *ξ* value. To overcome this, for the analytical calculations, we assume that players’ strategies are subject to execution errors with a small probability *ε*. Under this assumption, an intended act of cooperation may be mistakenly implemented as defection, and vice versa, with probability *ε* [11]. For *ε >* 0, a player’s strategy **s**_*p*_ effectively becomes **s**_(1−*ε*)*p*+*ε*(1−*p*)_. To this end, a pure cooperator strategy **s**_1_ becomes **s**_1−*ϵ*_ and a pure defector strategy **s**_0_ becomes **s**_*ε*_. Given the error probability, except **Q** = (1, 1, 1; 0, 0, 0), the Markov chain is ergodic for all deterministic transition vectors and the stationary distribution **u** is independent of the initial state. The case corresponding to **Q** = (1, 1, 1; 0, 0, 0) is not relevant either because transitions between games are not possible for such a transition vector, and the game remains in profitable state 1 for all times for our initial condition of all edges initially in game 1. Therefore, for the above-mentioned two classes of game transitions, we study competition between **s**_1−*ε*_ and **s**_*ε*_ following the same method described in sections 3.1-3.2.1 to calculate the value of *ξ*.

The *ξ* value calculated for game transitions without knowledge of states using the alternative method (section 3.2) matches exactly with values given in Table 1 for all but 6 of the 64 deterministic transition vectors. For the following six transition vectors: **(i) Q** = (0, 1, 0; 0, 0, 1), **(ii) Q** = (0, 1, 0; 1, 0, 0), **(iii) Q** = (0, 1, 1; 0, 0, 1), **(iv) Q** = (0, 1, 1; 1, 0, 1), **(v) Q** = (1, 1, 0; 1, 0, 0), and **(vi) Q** = (1, 1, 0; 1, 0, 1) our assumption of execution errors breaks down and the analytical value of *ξ* using the alternate method doesn’t exactly match with the results found using the basic calculation described in section 2. Nevertheless, the alternative method provides a simple framework for calculating the critical threshold (*b/c*)^∗^ for scenarios with and without state knowledge.

###### 3.2.2 Intuitive explanation based on the alternate method

Here, we study the game transition vector **Q**_**1**_ using the alternative method proposed in section 3.2 to understand how game transitions promote cooperation (in the absence of knowledge about states) when compared to a single game.

###### Q_**1**_ = (1, 0, 0; 1, 0, 0)

When two **s**_1_ players interact, the unique eigen vector **u**(**s**_1_, **s**_1_) for the transition vector **Q**_**1**_ can be written using Eq. 168,

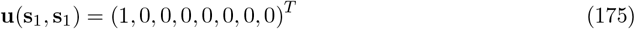

The payoff to each cooperator is *π*(**s**_1_, **s**_1_) = *b*_1_ − *c*.

When a **s**_1_ and a **s**_0_ player interacts, the unique eigen vector **u**(**s**_1_, **s**_0_) for the transition vector **Q**_**1**_ can be written using Eq. 169,

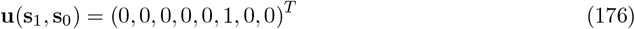

The payoff to the cooperator, *π*(**s**_1_, **s**_0_) = −*c* and the defector receives *π*(**s**_0_, **s**_1_) = *b*_2_.

The two defectors always receive a payoff of 0.

The effective payoff matrix (Eq. 171) between cooperators and defectors under the game transition vector **Q**_**1**_ becomes

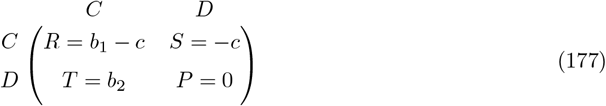

Here, *b*_1_ (*b*_2_) is the benefit of cooperation in game 1 (game 2), and we assume game 1 is more valuable than game 2 (*b*_1_ *> b*_2_). The payoff matrix (Eq. 177) shows that with game transitions between two donation games under the transition structure **Q**_**1**_, where only mutual cooperation leads to the more valuable state, evolution proceeds “as if” cooperators and defectors are playing an asymmetric game. Compared to playing only game 1 (where *R* = *b*_1_ −*c, S* = −*c, T* = *b*_1_, *P* = 0), unilateral defection brings a lesser payoff to the defector by a factor of Δ*b* = *b*_1_ −*b*_2_, whereas mutual cooperation brings the same payoff *b*_1_ − *c* to each cooperator (Eq. 177). Since *b*_1_ *> b*_2_, game transition gives cooperators an advantage over defectors, and as is evident, the advantage grows with the difference between the two games (Δ*b*).

Substituting Eq. 177 in Eq. 173, one can again show that the barrier to be overcome for cooperation to spread can be reduced by a factor 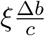, where *ξ* = (*k* −1)*/*2 for **Q**_**1**_. This matches exactly with the expression for *ξ* obtained from Eq. 161 using the basic method described in Section 1.

##### 3.3 Game transitions with complete knowledge of the states

When players possess complete knowledge of the state they are currently in, they adopt state-dependent strategies, 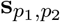 . In this scenario, four pure state-dependent strategies compete in the population: *CC* = **s**_1,1_, *CD* = **s**_1,0_, *DC* = **s**_0,1_, and *DD* = **s**_0,0_. In the following, using the alternate method, for a class of game transitions, we calculate the conditions under which **s**_1,1_ and **s**_1,0_ strategies are favored by selection when players have knowledge of the current state.

###### 3.3.1 Sigma rule for multiple strategies in structured population

For a game between *n* strategies, the payoff for an interaction between two strategies is given by an *n*× *n* payoff matrix, Π = [*π*_*ij*_], where *π*_*ij*_ is the payoff to the player using strategy *i* against a player using strategy *j*. For the competition between multiple strategies (*n >* 2), under weak selection (*δ* ≪ 1) and low mutation limit, Tarnita et al. [4] have shown that a strategy *l* is favored by selection if,

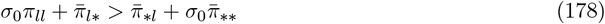

where 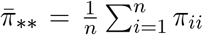, average payoff when both players use the same strategy; 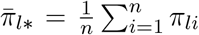 average payoff to strategy *l*; 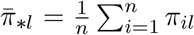, average payoff when playing against strategy *l*. Under a low mutation limit, for regular graphs and death-birth updating, *σ*_0_ = (*k* + 1)*/*(*k* − 1) is the same as *σ* defined in Eq. 173.

Below, using Eq. 178, we derive the conditions under which the strategies *CC*(= **s**_1,1_) and *CD*(= **s**_1,0_) are favored by selection in our model with *n* = 4 interacting state-dependent strategies.

###### Payoff calculation for competition between *CC* and *CC*-players

When two **s**_1,1_ players interact along an edge over several time-steps, using Eqs. 164-166 the unique eigen vector **u**(**s**_1,1_, **s**_1,1_) can be written as,

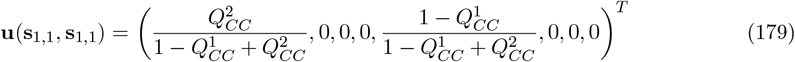

Substituting Eq. 179 in Eq. 167, we calculate the payoff of each *CC*-players as 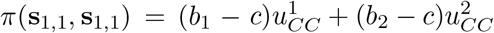, where 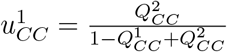, and 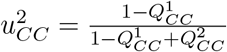.

###### Payoff calculation for competition between *CC* and *CD*-players

When a **s**_1,1_ player interacts with a **s**_1,0_ player along an edge over several time-steps, using Eqs. 164-166 the unique eigen vector **u**(**s**_1,1_, **s**_1,0_) can be written as,

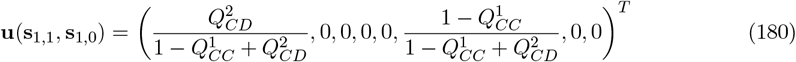

Substituting Eq. 180 in Eq. 167, we calculate the payoff of *CC*-player and *CD*-player as 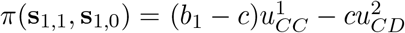 and 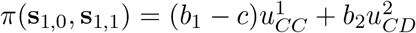, respectively, where 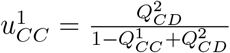, and 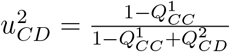.

###### Payoff calculation for competition between *CC* and *DC*-players

When a **s**_1,1_ player interacts with a **s**_0,1_ player along an edge over several time-steps, using Eqs. 164-166 the unique eigen vector **u**(**s**_1,1_, **s**_0,1_) can be written as,

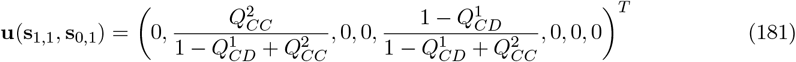

Substituting Eq. 181 in Eq. 167, we calculate the payoff of *CC*-player and *DC*-player as 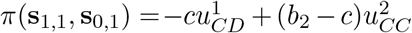 and 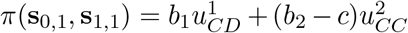, respectively, where 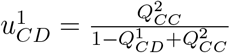 and 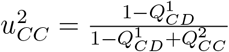.

###### Payoff calculation for competition between *CC* and *DD*-players

When a **s**_1,1_ player interacts with a **s**_0,0_ player along an edge over several time-steps, using Eqs. 164-166 the unique eigen vector **u**(**s**_1,1_, **s**_0,0_) can be written as,

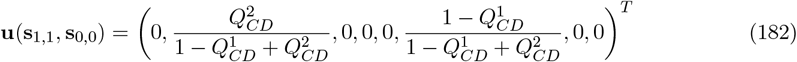

Substituting Eq. 182 in Eq. 167, we calculate the payoff of *CC*-player and *DD*-player as 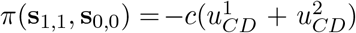 and 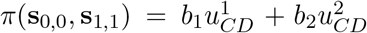, respectively, where 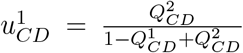, and 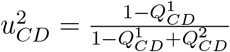.

###### Payoff calculation for competition between *CD* and *CD*-players

When two **s**_1,0_ players interact along an edge over several time-steps, using Eqs. 164-166 the unique eigen vector **u**(**s**_1,0_, **s**_1,0_) can be written as,

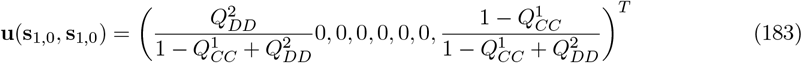

Substituting Eq. 183 in Eq. 167, we calculate the payoff of each *CD*-players 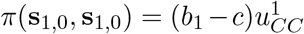, where 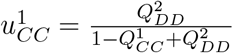.

###### Payoff calculation for competition between *CD* and *DC*-players

When a **s**_1,0_ player interacts with a **s**_0,1_ player along an edge over several time-steps, using Eqs. 164-166 the unique eigen vector **u**(**s**_1,0_, **s**_0,1_) can be written as,

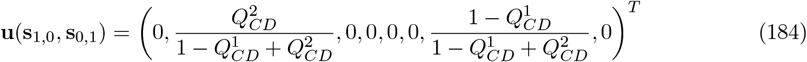

Substituting Eq. 184 in Eq. 167, we calculate the payoff of *CD*-player and *DC*-player as 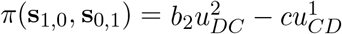 and 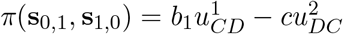, respectively, where 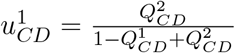, and 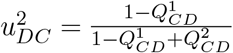.

###### Payoff calculation for competition between *CD* and *DD*-players

When a **s**_1,0_ player interacts with a **s**_0,0_ player along an edge over several time-steps, using Eqs. 164-166 the unique eigen vector **u**(**s**_1,0_, **s**_0,0_) can be written as,

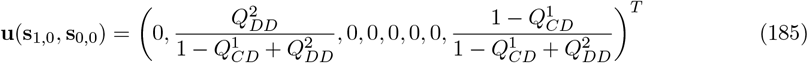

Substituting Eq. 185 in Eq. 167, we calculate the payoff of *CD*-player and *DD*-player as 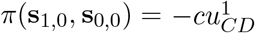 and 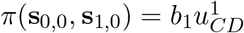, respectively, where 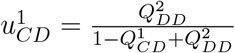.

###### Payoff calculation for competition between *DC* and *DC*-players

When two **s**_0,1_ players interact along an edge over several time-steps, using Eqs. 164-166 the unique eigen vector **u**(**s**_0,1_, **s**_0,1_) can be written as,

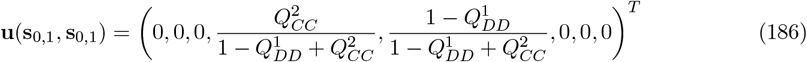

Substituting Eq. 186 in Eq. 167, we calculate the payoff of each *DC*-players 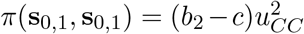, where 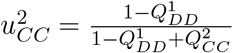.

###### Payoff calculation for competition between *DD* and *DD*-players

When two **s**_0,0_ players interact along an edge, both of the defectors receive a payoff of 0 in donation games.

In order to illustrate the usefulness of this alternative method, we show how to obtain the condition for the *CC* and *CD* strategies to be favored by selection using the example of the time-out game (**Q**_**2**_) used in the main text.

###### 3.3.2 Q_2_ = (1, 0, 0; 1, 1, 1)

Using Eq. 178, we can calculate the condition for a strategy *CC*(= **s**_1,1_) to be favored by selection when individuals adopt pure state-dependent strategies in the presence of complete knowledge of the states. Thus, strategy *CC* is favored by selection if,

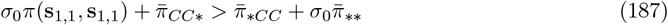

where,

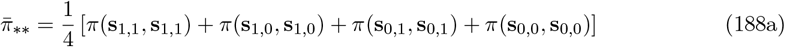

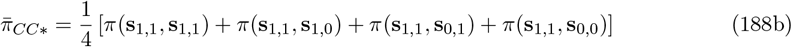

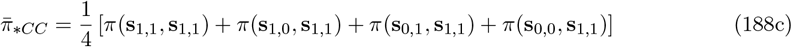

Using Eqs. 179-186 we calculate each payoff values for the transition vector **Q**_**2**_.

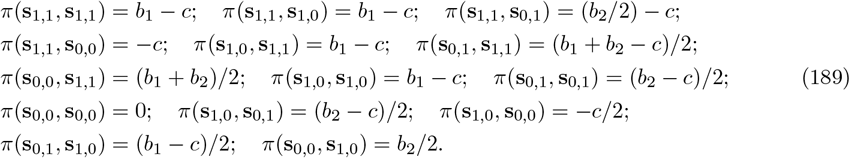

Substituting Eq. 189 in Eqs. 188a-188c, we get

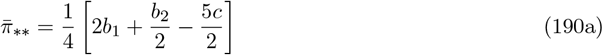

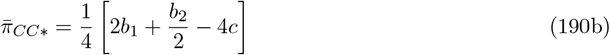

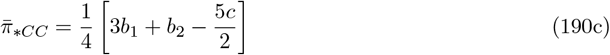

Putting Eqs. 190a-190c in the sigma rule (Eq. 187) and after doing some algebra, the condition for strategy *CC* to be favored by selection under death-birth updating is

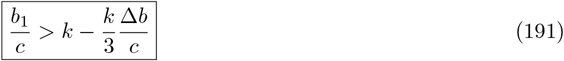

The above condition matches exactly with Eq. 126, calculated using the method described in section Eq. 191 also corresponds to Eq. 6 in the main text with 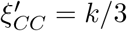 for the transition vector **Q**_**2**_.

Analogously, strategy *CD* is favored by selection if,

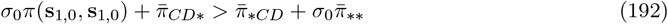

where,

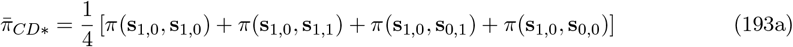

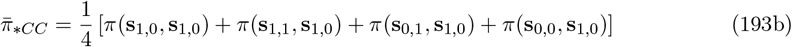

Again, substituting Eqs. 189 in Eqs. 188a,193a-193b, we get

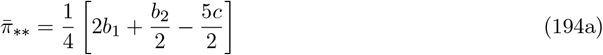

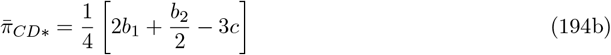

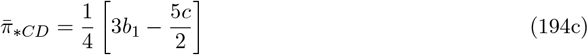

Putting Eqs. 194a-194c in the sigma rule (Eq. 192), and after doing some algebra, we get the condition for strategy *CD* to be favored by selection under death-birth updating is

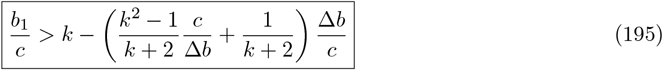

The above condition matches exactly with Eq. 135, calculated using the method described in section 1. Eq. 195 also corresponds to Eq. 6 in the main text with 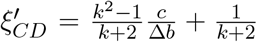 for the transition vector **Q**_**2**_.

The alternative method described in Section 3.3 for analyzing game transitions with knowledge of the states applies only to a specific class of deterministic transition structures that satisfy the following constraints.

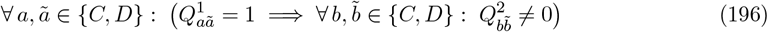

For competition between pure state-dependent strategies, if a transition vector does not satisfy the constraint in Eq. 196, the resulting Markov chain is non-ergodic, and the invariant distribution between any two strategies is no longer unique. Because of this constraint, the alternative method applies only to a limited set of deterministic transition vectors. However, for arbitrary stochastic transition vectors 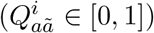 and stochastic strategies (*p*_1_, *p*_2_ ∈ [0, 1]), the alternative method provides a simpler way to compute the selection condition for any state-dependent strategies compared with the basic method described in Section 1.

### 4 Evolutionary dynamics with imperfect environmental state knowledge

In our model with imperfect knowledge of the environmental states, players can adopt one of the four possible strategies, *CC*(= **s**_1,1_), 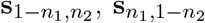, and *DD*(= **s**_0,0_). A player with strategy 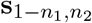 at each time step, cooperates with probability 1 − *n*_1_ in state 1 and with probability *n*_2_ in state 2. We want to understand how noise in state perception (*n*_1_ for state 1, and *n*_2_ for state 2) affects the evolutionary dynamics of cooperation. Here, we find the condition for the *CC* strategy to be favored by selection under the natural transition vector **Q**_**1**_ = (1, 0, 0; 1, 0, 0) when individuals have imperfect knowledge of states. We will follow the method described in section 3.3 with our modified strategies to calculate the critical threshold for the success of *CC*. Using the sigma rule for multiple strategies in the structured population, strategy *CC* is favored by selection under the death-birth update if,

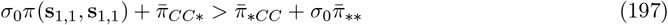

where,

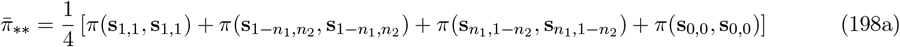

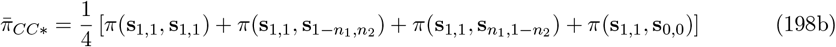

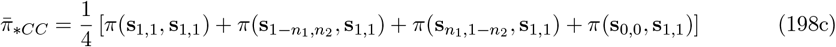

Using Eq. 167, we calculate the payoff between each strategy as follows,

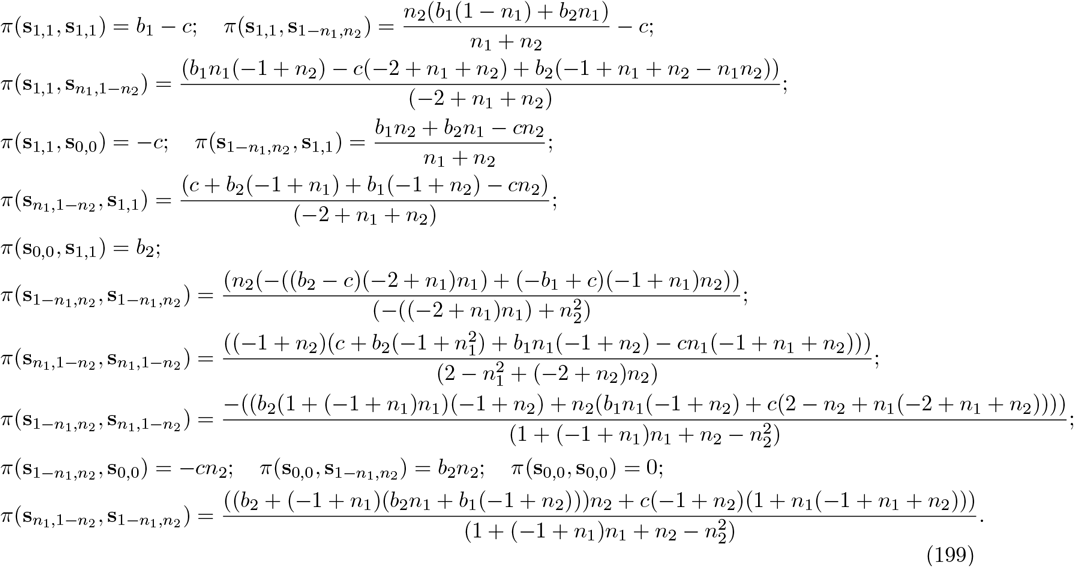

Substituting Eq. 199 into Eqs. 198a-198c and then using the sigma rule (Eq. 197), we get,

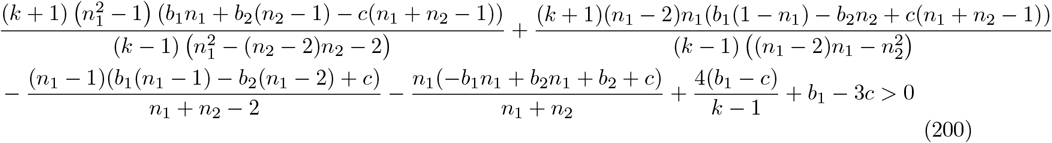

Eq. 200 gives the selection condition for the *CC*−strategy under the transition vector **Q**_**1**_ for arbitrary values of noise in state perception (*n*_1_, *n*_2_ ∈ (0, 1]). Note that we recover our old expression (Eq. 104) for the *CC*−strategy to be favored by selection under complete state knowledge in the sequential limit *n*_1_ → 0 followed by *n*_2_ → 0; reversing the order of limits yields a different expression.

## 5 Supplementary figures

**Figure S1.**
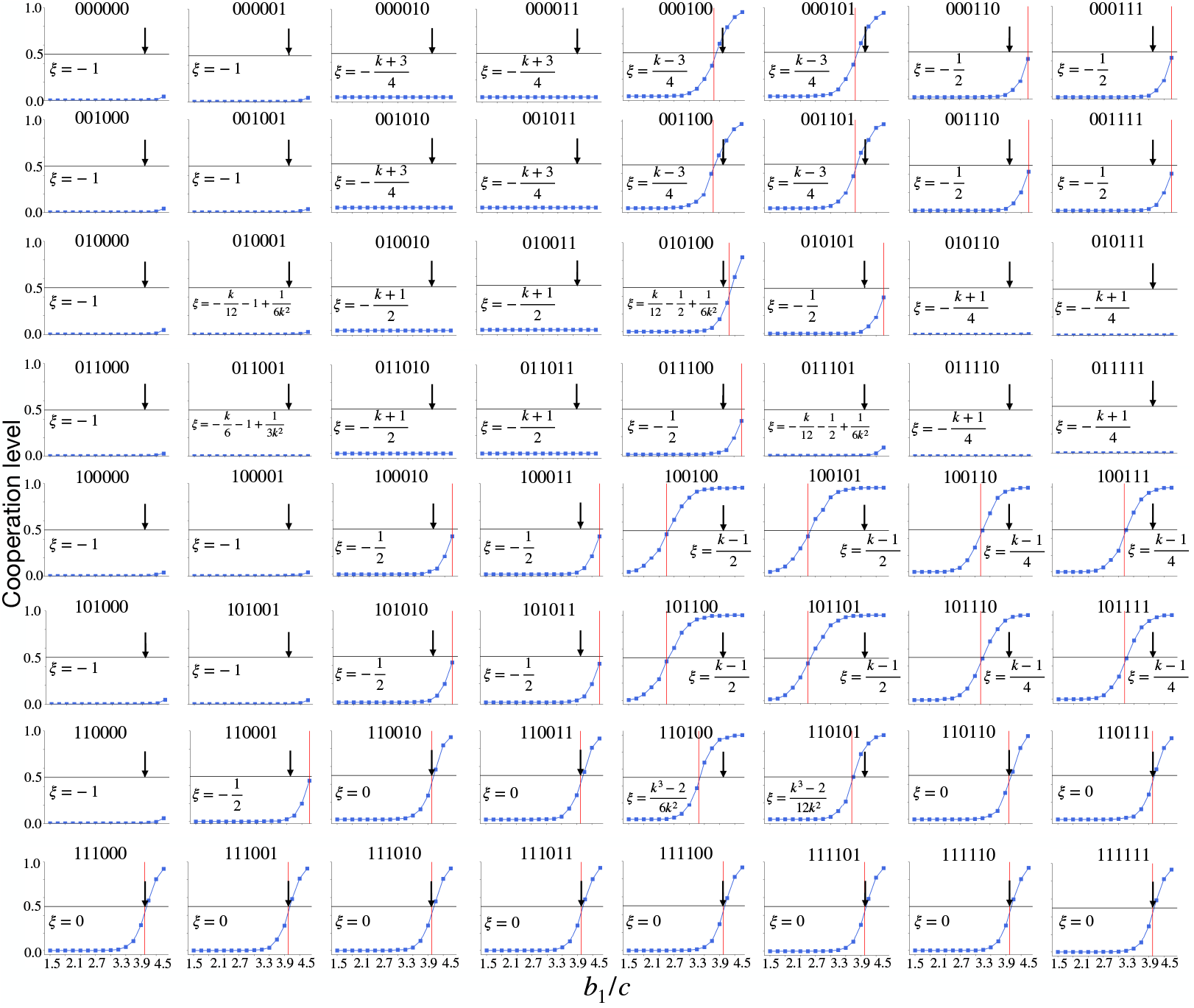
Evolutionary dynamics with game transitions for all 64 transition rules in the no knowledge scenario under death-birth update: Transition between 2 donation games is considered with game 1 being more beneficial than game 2 (*b*_1_ *> b*_2_). The critical benefit-to-cost ratio, (*b*_1_*/c*)^∗^, for the cooperation level to exceed 0.5, obtained from the theoretical analysis, is denoted by the vertical red line. Note that the analytical value of *ξ* (*>* 0, = 0, *<* 0), which is used to determine (*b*_1_*/c*)^∗^, indicates positive, neutral, or negative impact of game transitions on the evolution of cooperation. The blue dots show the cooperation level for different (*b*_1_*/c*) values, obtained from simulations. The black arrow corresponds to the critical benefit-to-cost ratio ((*b*_1_*/c*)^∗^ = *k*) for game 1 only, without any transitions. Each simulation starts with an equal number of *C* and *D* players playing game 1 initially and runs until the population reaches fixation (all *C* or all *D*), with each point averaged over 10^4^ independent trials. Parameter values: *N* = 500, *k* = 4, *δ* = 0.01, *c* = 1, and *b*_2_ = *b*_1_ − 1.

**Figure S2.**
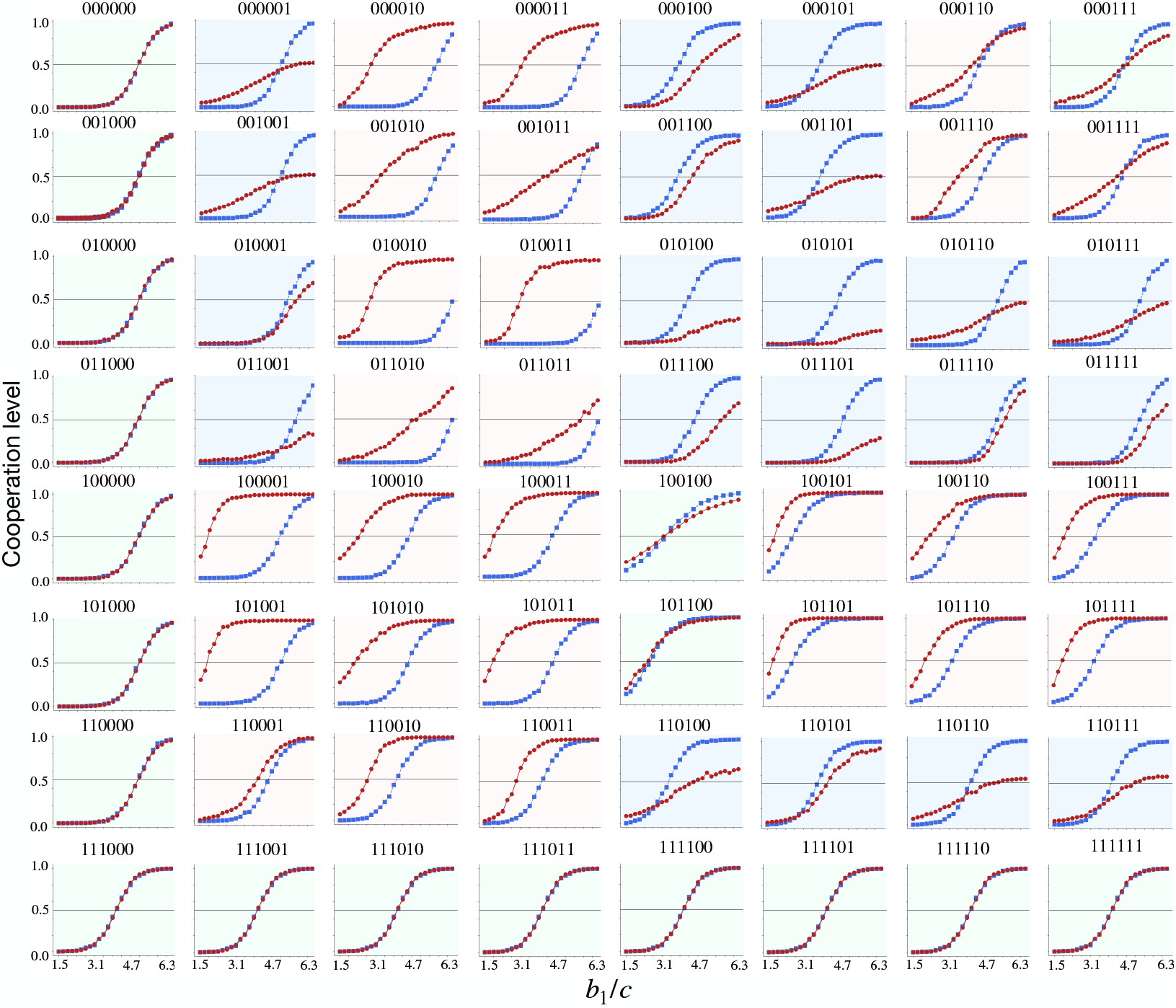
Comparison between complete and no knowledge scenario on the evolutionary dynamics with game transitions for all deterministic transition rules under death-birth update: The figures show cooperation levels for the games with (red lines) and without state knowledge (blue lines) as a function of the benefit-to-cost ratio in game 1 while keeping the difference of benefit of two games Δ*b* fixed. Different background colors are used to indicate the qualitative behavior for all the 2^6^ = 64 deterministic transition rules: a pale red background denotes instances where state-dependent behavior (complete knowledge) outperforms state-independent behavior (no knowledge scenario), a pale blue background denotes cases where state-independent behavior allows cooperation to dominate for a lower threshold of *b*_1_*/c*, and the pale green background denotes cases with neutral outcomes. All other parameter values are the same as Fig. S1.

**Figure S3.**
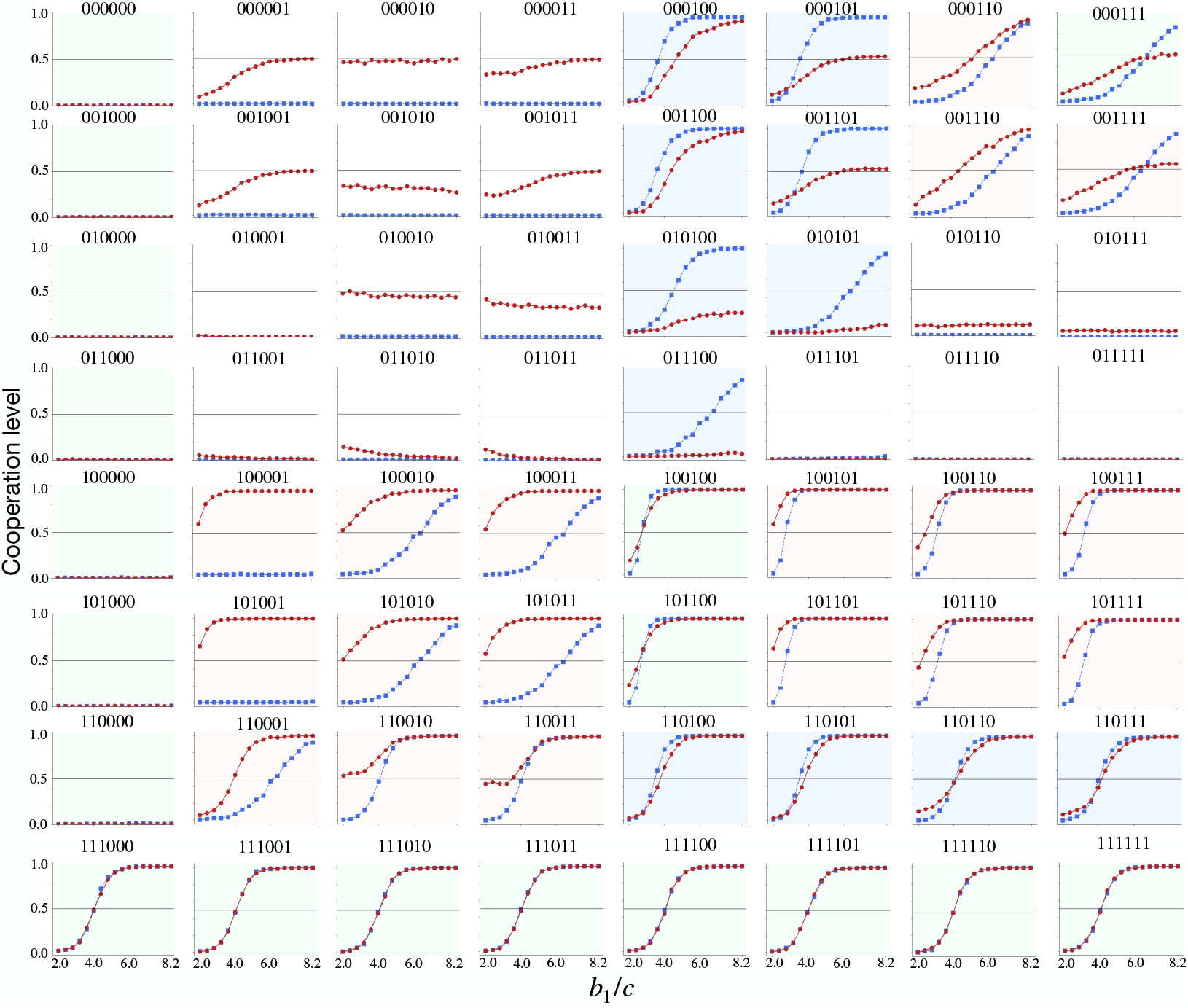
Analysis of all deterministic transition rules for fixed benefit in game 2 (*b*_2_). We plot cooperation levels for the games with state knowledge (red lines) and without knowledge (blue lines) as a function of the benefit-to-cost ratio in game 1, keeping *b*_2_ fixed while Δ*b* increases due to increasing *b*_1_. We use the same color scheme for the background as before (pale red: benefit conferred by knowledge about the state, pale blue: benefit of ignorance, and pale green: neutral). When *b*_2_ is fixed to a small value, cooperation can not evolve if mutual cooperation is not rewarded in any state, or if its rewarded only in the less beneficial state, while unilateral defection is rewarded (panels with a white background). In such situations, the benefit of cooperation in game 2 is too small for the evolution of cooperation with or without state knowledge. For all other transition rules, the results are consistent with Fig. S2. We used *b*_2_ = 1.8. All other parameter values are the same as Fig. S1.

**Figure S4.**
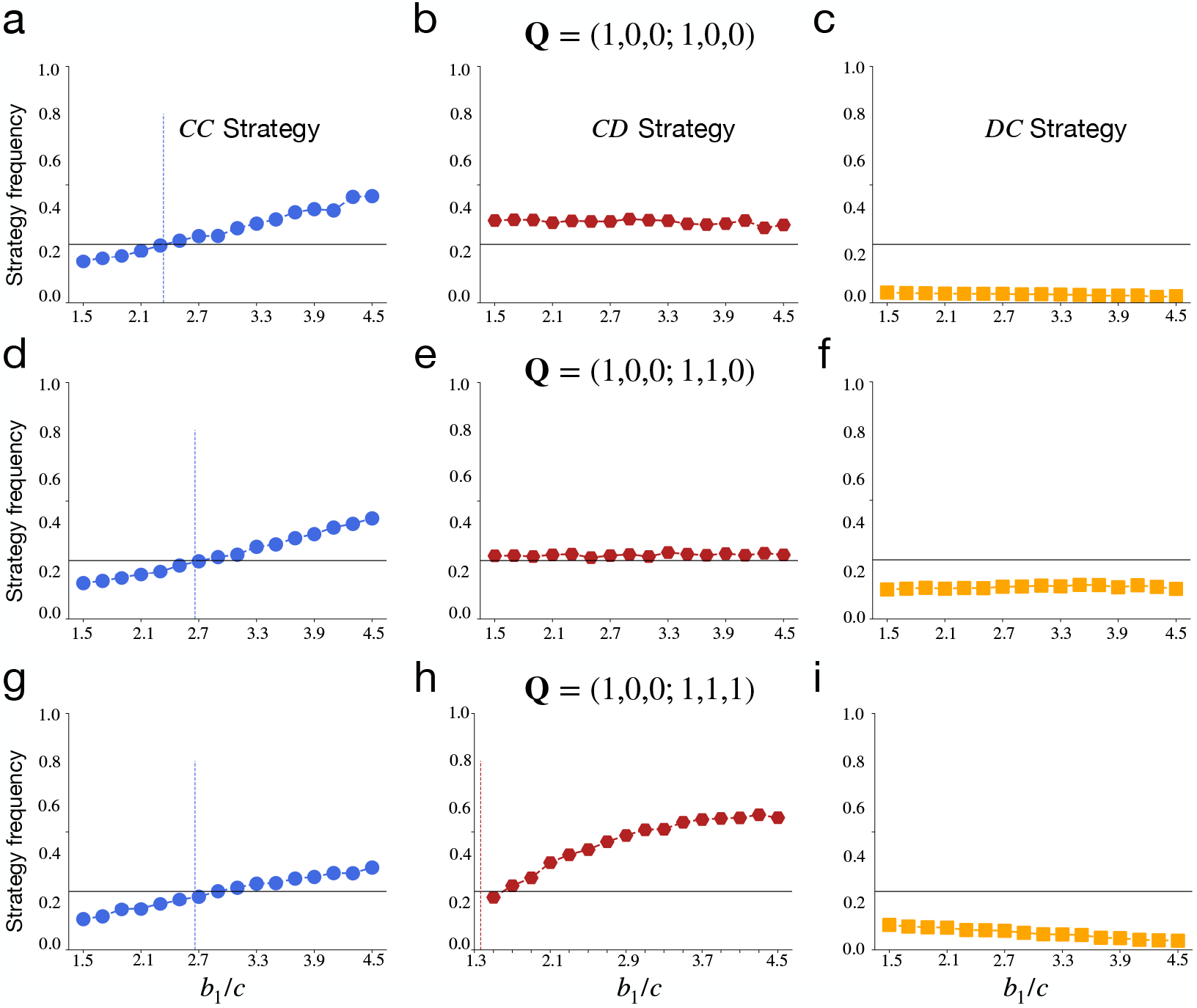
Evolutionary dynamics in the presence of rare mutations: We study the death-birth update on random regular graphs of degree *k*. With probability 1 −*µ*, the empty site is occupied by the randomly selected neighbor’s offspring. With probability *µ*, the empty site is occupied by one of the 4 available strategies *CC, CD, DC*, or *DD* with equal probability. The frequency of a strategy **s** in the stationary distribution, ⟨*p*_**s**_⟩, is used to measure the success of that strategy. A strategy **s** is favored by selection if ⟨*p*_**s**_⟩ *>* 1*/*4. We obtain each data point by averaging ⟨*p*_**s**_⟩ over 500 independent trials. For each trial, ⟨*p*_**s**_⟩ is obtained by time-averaging the fraction of strategy **s** present in the population in the last 2 × 10^7^ time steps of the total 4 × 10^7^ steps. The points of intersection between the dots and the horizontal lines mark the critical benefit-to-cost ratios 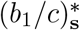 for a strategy **s** to be favored by selection ( ⟨*p*_**s**_⟩ *>* 1*/*4). Vertical dashed lines denote the analytical value of 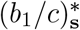 ; their absence in some cases means the analytical threshold lies outside the displayed range of *b*_1_*/c* values. Other parameters are: *N* = 100, *k* = 4, *δ* = 0.01, *µ* = 10^−4^, *b*_2_ = *b*_1_ − 1, *c* = 1.

**Figure S5.**
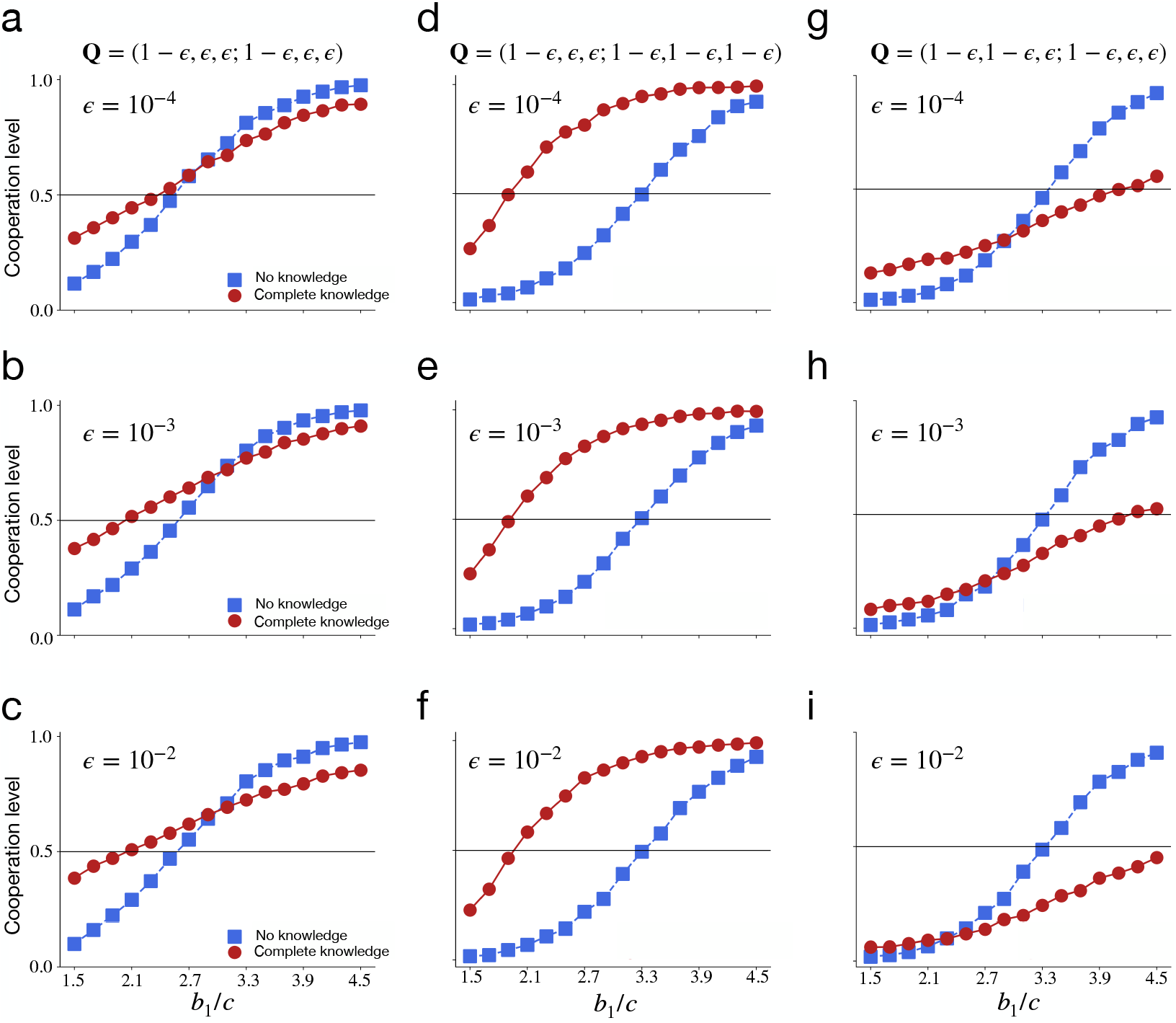
Evolutionary dynamics under stochastic transition vectors: Each column corresponds to a stochastic version of the three transition vectors **Q**_**1**_, **Q**_**2**_, and **Q**_**3**_ considered in the main text. Here, transitions occur between states with a finite probability of *ϵ* or 1 −*ϵ* compared to their deterministic parts (where *ϵ* = 0). Each row corresponds to different values of *ϵ*. The plots show how the cooperation level changes with increasing *b*_1_*/c* ratio. The stochastic transition vectors are shown at the top of each column. For the transition vectors **Q**_**2**_ = (1, 0, 0; 1, 1, 1) (panels **d, e, f** ) and **Q**_**3**_ = (1, 1, 0; 1, 0, 0) (panels **g, h, i**), introducing stochasticity in state transition probability does not alter the evolutionary outcomes relative to its analogous deterministic part. However, for the transition vector **Q**_**1**_ = (1, 0, 0; 1, 0, 0) (panels **a, b, c**), stochasticity in state transition promotes cooperation by lowering the critical threshold in the complete knowledge scenario relative to the no knowledge scenario. Parameters used are the same as in Fig. S1.

**Figure S6.**
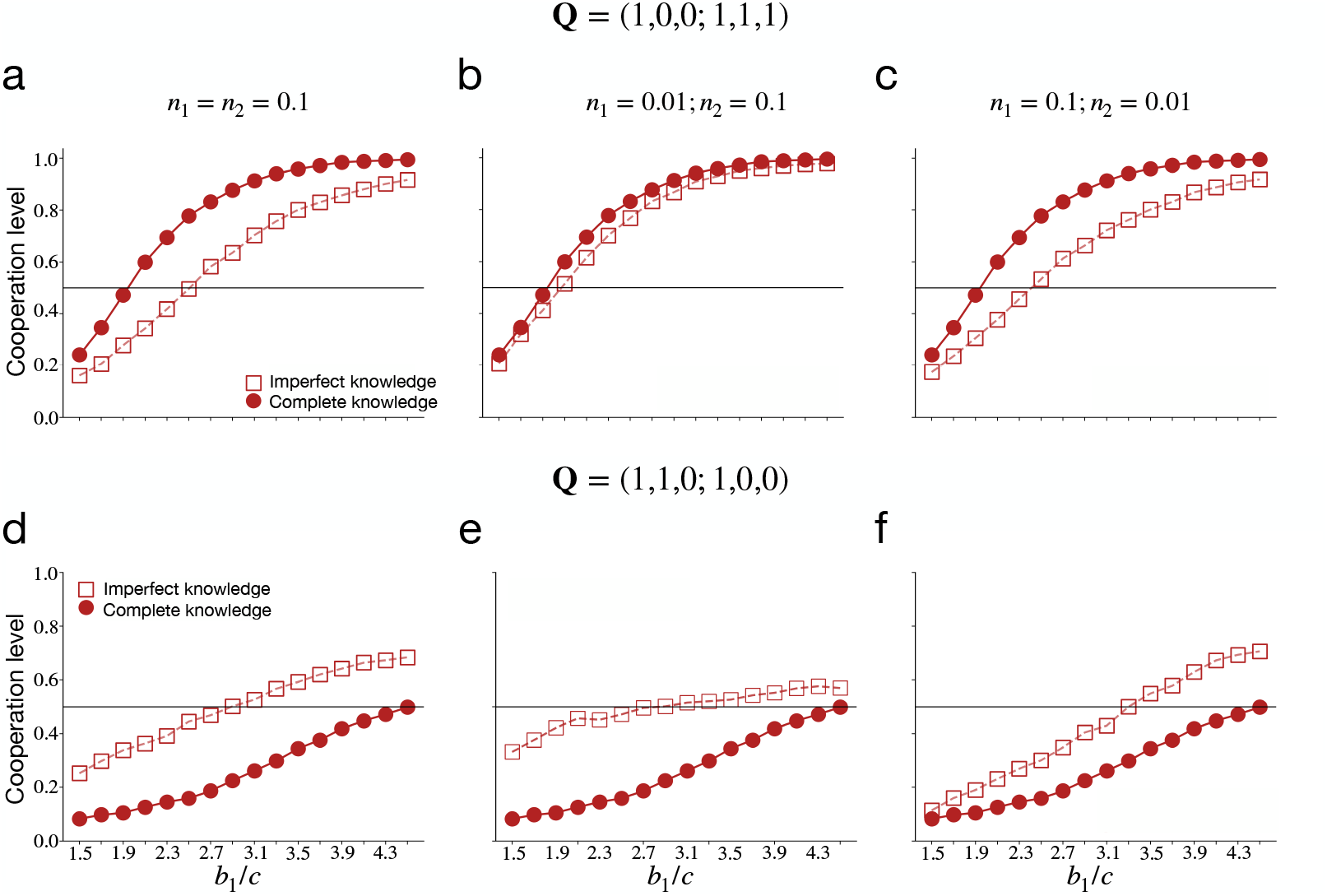
Evolutionary dynamics with imperfect state knowledge. The top (**a, b, c**) and bottom (**d, e, f** ) panels show the results for two different transition rules. Noise levels in the bottom panels **d, e, f** are the same as those shown in the top panels **a, b, c**, respectively. Parameter values are the same as in Fig. S1.

